# Micro-Meta App: an interactive software tool to facilitate the collection of microscopy metadata based on community-driven specifications

**DOI:** 10.1101/2021.05.31.446382

**Authors:** Alex Rigano, Shannon Ehmsen, Serkan Utku Ozturk, Joel Ryan, Alexander Balashov, Mathias Hammer, Koray Kirli, Karl Bellve, Ulrike Boehm, Claire M. Brown, James J. Chambers, Robert A. Coleman, Andrea Cosolo, Orestis Faklaris, Kevin Fogarty, Thomas Guilbert, Anna B. Hamacher, Michelle S. Itano, Daniel P. Keeley, Susanne Kunis, Judith Lacoste, Alex Laude, Willa Ma, Marco Marcello, Paula Montero-Llopis, Glyn Nelson, Roland Nitschke, Jaime A. Pimentel, Stefanie Weidtkamp-Peters, Peter J. Park, Burak Alver, David Grunwald, Caterina Strambio-De-Castillia

## Abstract

For the information content of microscopy images to be appropriately interpreted, reproduced, and meet FAIR (Findable Accessible Interoperable and Reusable) principles, they should be accompanied by detailed descriptions of microscope hardware, image acquisition settings, image pixel and dimensional structure, and instrument performance. Nonetheless, the thorough documentation of imaging experiments is significantly impaired by the lack of community-sanctioned easy-to-use software tools to facilitate the extraction and collection of relevant microscopy metadata. Here we present **Micro-Meta App**, an intuitive open-source software designed to tackle these issues that was developed in the context of nascent global bioimaging community organizations, including **B**io**I**maging **N**orth **A**merica (BINA) and **QUA**lity Assessment and **REP**roducibility in **Li**ght **Mi**croscopy (QUAREP-LiMi), whose goal is to improve reproducibility, data quality and sharing value for imaging experiments. The App provides a user-friendly interface for building comprehensive descriptions of the conditions utilized to produce individual microscopy datasets as specified by the recently proposed 4DN-BINA-OME tiered-system of Microscopy Metadata model. To achieve this goal the App provides a visual guide for a microscope-user to: 1) interactively build diagrammatic representations of hardware configurations of given microscopes that can be easily reused and shared with colleagues needing to document similar instruments. 2) Automatically extracts relevant metadata from image files and facilitates the collection of missing image acquisition settings and calibration metrics associated with a given experiment. 3) Output all collected Microscopy Metadata to interoperable files that can be used for documenting imaging experiments and shared with the community. In addition to significantly lowering the burden of quality assurance, the visual nature of Micro-Meta App makes it particularly suited for training users that have limited knowledge of the intricacies of light microscopy experiments. To ensure wide-adoption by microscope-users with different needs Micro-Meta App closely interoperates with **MethodsJ2** and **OMERO.mde**, two complementary tools described in parallel manuscripts.

## Background

The establishment of community-driven, shared documentation and quality control specifications for light microscopy would allow to appropriately document imaging experiments, minimize errors and quantify any residual uncertainty associated with each step of the procedure (*1–6*). In addition to providing essential information about the provenance (i.e., origin, lineage) (*7, 8*) of microscopy results, this would make it possible to faithfully interpret scientific claims, facilitate comparisons within and between experiments, foster reproducibility, and maximize the likelihood that data can be re-used by other scientists for further insight (*5, 6, 9, 10*). First and foremost, such information would serve to facilitate the compilation of accurate Methods sections for publications that utilize the quantitative power of microscopy experiments to answer scientific questions (*11–13*). Furthermore, it would provide clear guidance to the manufacturers of microscopy instruments, hardware components, and processing software about what information the scientific community requires to ensure scientific rigor so that they can be automatically provided during acquisition and written in the headers of image files. Last but not least, machine-actionable versions of the same information (*14*) could be provided alongside image datasets on the growing number of public image data resources (*3*) that allow the deposition of raw image data associated with scientific manuscripts, a promise to emulate for light microscopy the successful path that has led to community standards in the field of genomics (*15–19*) (e.g., the IDR (*20*), EMPIAR (*21*), and Bioimage Archive (*22*) hosted at the EMBL – EBI; the European Movincell (*23*); the Japanese SSBD hosted by RIKEN (*24*); and, in the USA, the NIH-funded Cell Image Library (*25, 26*), BRAIN initiative’s imaging resources (*27*), the Allen Cell Explorer (*28*), and the Human Cell Atlas (*29–32*)).

In order to promote the development of shared community-driven Microscopy Metadata standards, the NIH funded 4D Nucleome (4DN) (*33, 34*) and the Chan Zuckerberg Initiative (CZI) funded BioImaging North America (BINA) Quality Control and Data Management Working Group (QC-DM-WG) (*35*) have recently proposed the 4DN-BINA-OME (NBO) a tiered-system for Microscopy Metadata specifications (*36–39*). The 4DN-BINA-OME specifications lay the foundations for upcoming community-sanctioned standards being developed in the context of the Metadata Working Group (WG7) of the QUAlity Assessment and REProducibility for Instrument and Images in Light Microscopy (QUAREP-LiMi) initiative (quarep.org) (*4, 40*). Their purpose is to provide a scalable, interoperable and Open Microscopy Environment (OME) (*41–43*) Next-Generation File Format (NGFF) (*44*) compatible framework for light microscopy metadata guiding scientists as to what provenance metadata and calibration metrics should be provided to ensure the reproducibility of different categories of imaging experiments.

Despite their value in indicating a path forward, guidelines, specifications, and standards on their own lack the one essential feature that would make them actionable by experimental scientists faced with the challenge of producing well-documented, high-quality, reproducible and re-usable datasets: namely easy-to-use software tools or even better-automated pipelines to extract all available metadata from microscope configuration and image data files.

While some advances have been proposed, such as OMERO.forms (*45*), PyOmeroUpload (*46*) and MethodsJ (*5*), these tools only offer limited functionalities, are not integrated with community standards and are not per se future proof. To provide a way forward, in this and in two related manuscripts, we present a suite of three interoperable software tools (Supplemental Figure 1) that were developed to provide highly complementary, intuitive approaches for the bench-side collection of Image Metadata, with particular reference to Experimental Metadata and Microscopy Metadata (*37, 38*). In two related manuscripts, we describe: 1) **OMERO.mde**, which is highly integrated with the widely used OMERO image data management repository and emphasizes the development of flexible, nascent specifications for experimental metadata (*47–49*); and 2) **MethodsJ2** (*50*), which is designed as an ImageJ plugin and emphasizes the consolidation of metadata from multiple sources and automatic generation of Methods sections of scientific publications.

In this manuscript, we present **Micro-Meta App** (Figure 1), which works both as a stand-alone app on the user’s desktop and as an integrated resource in third party web data portals. It offers a visual guide to navigate through the different steps required for the rigorous documentation of imaging experiments (Figures 2–4) as sanctioned by community specifications such as the 4DN-BINA-OME (NBO) Microscopy Metadata specifications that were recently put forth to extend the OME Data Model (*36–38, 51*).

**Figure 1.**
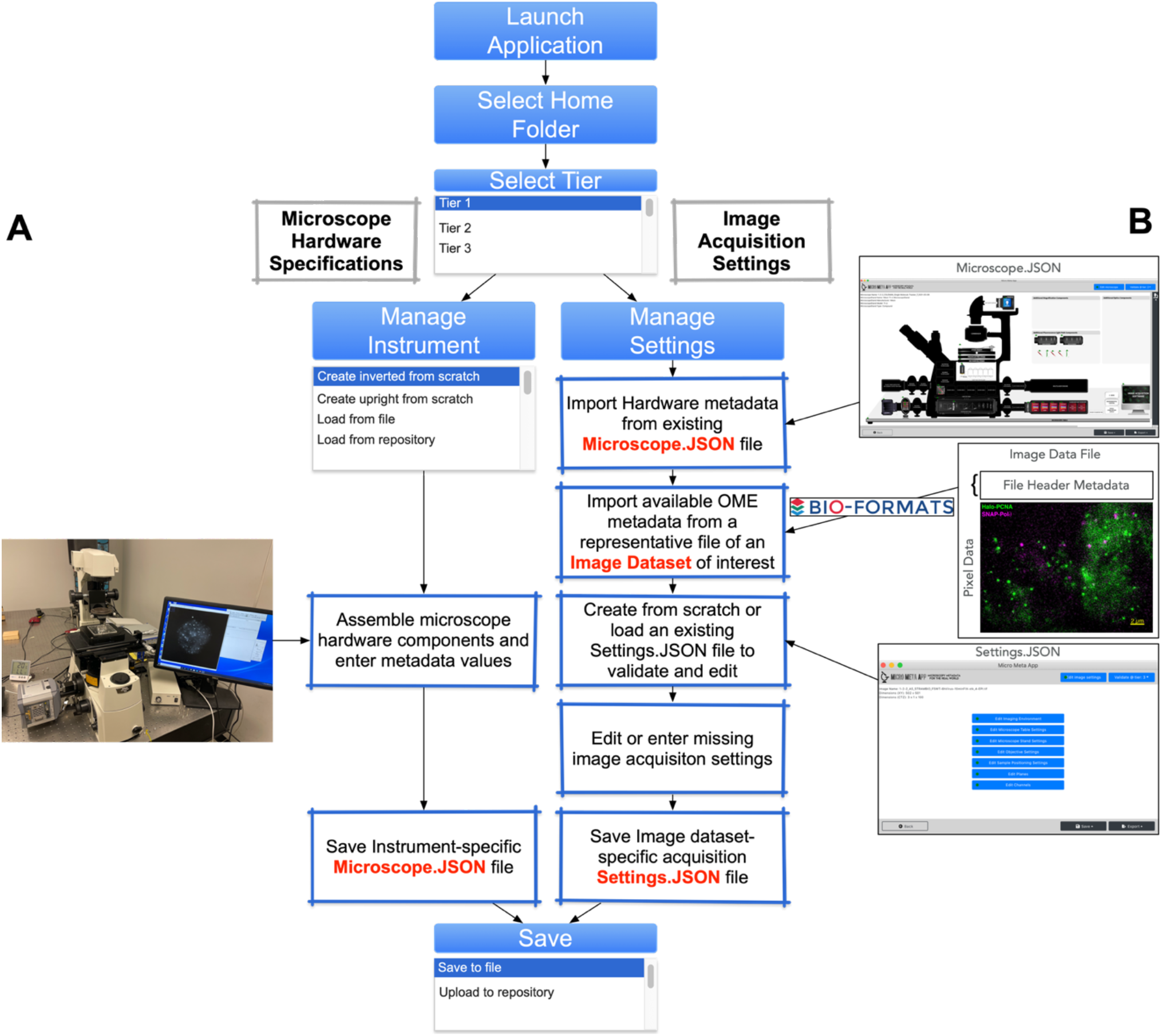
Micro-Meta App data processing workflows. Flow-chart depicting the two data-processing workflows available in Micro-Meta App. **A)** After selecting the documentation tier-level [1] appropriate for a given experiment-design and instrument-complexity as sanctioned by the 4DN-BINA-OME Microscopy Metadata Specifications (38), in the ***Manage Instrument*** section of the App the user is visually guided through the steps (Figures 2 and 3) required for the de novo creation a new Microscope hardware specifications file containing a list of components and associated 4DN-OME-BINA-specified metadata values. Alternatively, an existing Microscope-metadata file can be opened for validation, editing or revision. At the end of a given *Manage Instrument* session, the gathered hardware specifications metadata information is then saved in a *Microscope JSON* file that is used in the *Manage Settings* section of the App as the basis for the documentation of any image dataset acquired using the microscope in question. **B)** When microscope users are ready to document the acquisition-settings associated with a given image dataset, the ***Manage Settings*** section of Micro-Meta App (Figure 1) is used to: 1) choose the appropriate documentation Tier-level (38); 2) select an existing *Microscope.JSON* file describing the hardware configuration of the microscope used for acquisition; 3) open a representative image data file from the dataset of interest, and leverage BioFormats (43) to automatically retrieve available OME-compatible microscopy metadata from the header of the file; and 4) de novo create (or edit) a *Settings.JSON* file containing a detailed 4DN-BINA-OME-sanctioned description of the image acquisition metadata. This file is then saved in association with the image dataset either on the user’s desktop or in an available third-party data portal, and, as an example, can be used to automatically produce Methods text using MethodsJ2 (50). Example Microscope.JSON, Settings.JSON and associated image data file are available at: https://doi.org/10.5281/zenodo.4891883.

## Methods: Implementation and Availability

Micro-Meta App is available in two JavaScript (JS) implementations. The first was designed to facilitate the incorporation of the software in existing third party web portals (i.e., the 4DN Data Portal) (*34, 52*) and was developed using the JavaScript React library, which is widely used to build web-based user interfaces. Starting from this version, a stand-alone version of the App was developed by wrapping the React implementation using the JavaScript Electron library, with the specific purpose of lowering the barrier of adoption of the tool by labs that do not have access or prefer not to use imaging databases. More details about the implementation of Micro-Meta App are available in Supplemental Material.

In order to promote the adoption of Micro-Meta App, incorporation in third party data portals and re-use of the source code by other developers, the executables and source code for both Javascript React and Electron implementations of Micro-Meta App are available on GitHub (*53, 54*). In addition, a website describing Micro-Meta App (*55*) was developed alongside complete documentation and tutorials (*56*).

## Results + Discussion

### Micro-Meta App: an intuitive, highly visual interface to facilitate microscopy metadata collection

While the establishment of data formats, metadata standards and QC procedures is important, it is not *per se* sufficient to make sure reporting and data quality guidelines are adopted by the community. To ensure their routine utilization, it is, therefore, necessary to produce software tools that expedite QC procedures and image data documentation and make it straightforward for investigators to reproduce results and make decisions regarding the utility of specific datasets for addressing their specific questions. However, despite the availability of the OME Data Model and Bioformats (*41, 43*), the lack of standards has resulted in a scarce adoption of minimal information criteria and as a result the metadata provided by instrument and software manufacturers is scarce (Supplemental Table I and II).

Micro-Meta App was developed to address this unmet need. Micro-Meta App consists of a Graphical User Interface (GUI)-based open-source and interoperable software tool to facilitate and (when possible) automate the annotation of fluorescence microscopy datasets. The App provides an interactive approach to navigate through the different steps required for documenting light microscopy experiments based on available OME-compatible community-sanctioned tiered systems of specifications. Thus, Micro-Meta App is not only capable of adapting to varying levels of imaging complexity and user experience but also to evolving data models that might emerge from the community. At the time of writing, the App implements the Core of the OME Data Model and the tiered 4DN-BINA-OME Basic extension (*36–38, 51*). Efforts to implement the current Confocal and Advanced as well as the Calibration and Performance 4DN-BINA-OME extensions are underway (see Future Directions). To achieve this goal, Micro-Meta App is organized around two highly related data processing flows (Figure 1):

1. In the *Manage Instrument* modality, the App guides the users through the steps required to interactively build a diagrammatic representation of a given Microscope (Figures 2A and 3) by dragging-and-dropping individual components onto the workspace and entering the relevant attribute values based on the tier-level that best suites the microscope modality, experimental design, instrument complexity, and image analysis needs (*38*).
2. From this, Micro-Meta App automatically generates structured descriptions of the microscope *Hardware Specifications* and outputs them as interoperable *Microscope.JSON files* (example available on Zenodo as illustrated in Supplemental Material) (*57*) that can be saved locally, used by existing third-party web-portals (*52*), integrated with other software tools (MethodsJ2) and shared with other scientists, thus significantly lowering the manual work required for rigorous record-keeping and enabling rapid uptake and widespread implementation.
3. When user is ready to collect metadata to document the acquisition of specific image data sets, the *Manage Settings* section of the App automatically imports *Hardware Specifications* metadata from previously-prepared Microscope.JSON files and uses the BioFormats library (*43*) to extract available, OME-compatible metadata from an image data file of interest. From this basis, the App interactively guides the user to enter missing metadata specifying the tier-appropriate *Settings* used for a specific *Image Acquisition* session (Figures 2B and 4).
4. As a final step, the App generates interoperable paired Microscope- and Settings-JSON files (example available on Zenodo as illustrated in Supplemental Material) (*57*) that together contain comprehensive documentation of the conditions utilized to produce individual microscopy datasets and can be stored locally or integrated by third-party data portals (i.e., the 4D Nucleome Data Portal) (*58*).

**Figure 2.**
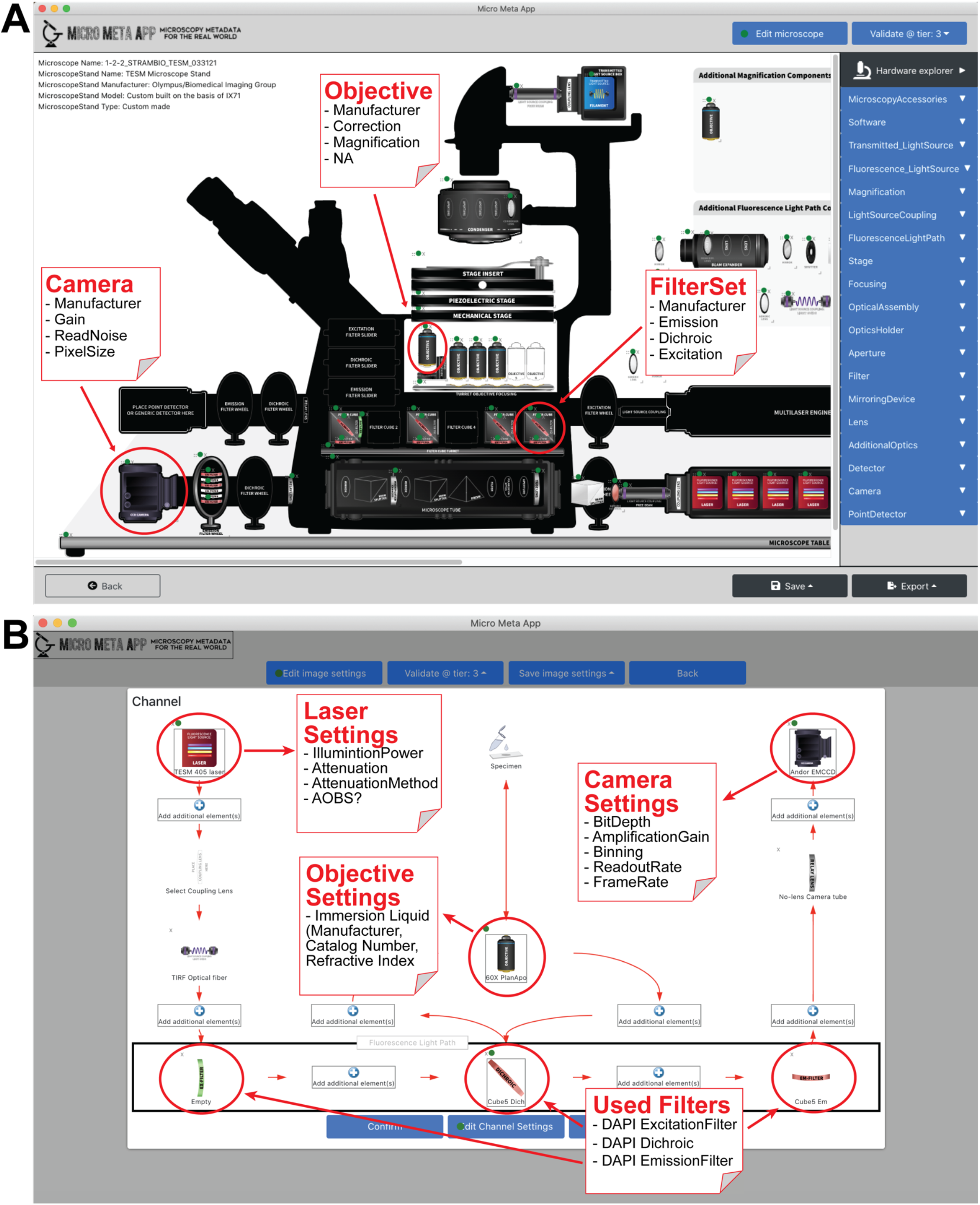
Micro-Meta App allows the intuitive and interactive documentation of imaging experiments. **A)** Using the Manage Instrument (Figure 1) section of Micro-Meta App, microscope user and custodians can visually create accurate representations of the hardware configuration of a given light microscope based on a specific documentation Tier-level, as sanctioned by as sanctioned by the 4DN-BINA-OME Microscopy Metadata model (38). Specifically, the App allows users to drag-and-drop individual hardware components (i.e., Light Sources, Objectives, Filters, Detectors, Lens, etc.,) onto the App workspace, and subsequently enter Tier-specific 4DN-BINA-OME-specified metadata values associated with each of the components to be saved in a reusable *Microscope.JSON* file. In the depicted example, the user clicks on icons representing a given Objective, Filter Set and Camera component and enters 4DN-BINA-OME-sanctioned Hardware Specifications metadata values associated with each of the items (e.g., *Objective Manufacturer, Model, Magnification* and *Numerical Aperture*). **B)** Using the Manage Settings section of Micro-Meta App the user opens a previously created Microscope metadata file and an image whose acquisition has to be documented. From here the App visually guides the user through the steps required to inspect, validate, and (if needed) enter missing 4DN-BINA-OME specified image acquisition metadata that complements information, which is automatically retrieved from the header of image files by using BioFormats. In the example, at each position along the Channel light path the user selects the specific component that was used among those available (e.g., Light Source, Filters, Objectives, and Detectors) and then enters the specific settings that were applied during image acquisition (e.g., *Camera Bit Depth, Amplification Gain, Binning,* etc.) as specified by the 4DN-BINA OME model (38).

Depending on the specific implementation of the Micro-Meta App being used (see Implementation section), the workflow varies slightly. The discussion below refers specifically to the stand-alone version of Micro-Meta App implemented in JavaScript Electron.

#### Manage Instrument Hardware

The purpose of this section of Micro-Meta App is to guide microscope users and custodians in the creation of accurate but at the same time intuitive and easy-to-generate visual depictions of a given microscope. This is done while collecting relevant information for each hardware component that scales with experimental intent, instrument complexity and analytical needs of individual imaging experiments depending on tier-levels sanctioned by the 4DN-BINA-OME Microscopy Metadata specifications (*36–38, 51*). Specifically, the workflow (Figure 3) is composed of the following steps:

1. After launching the application, the user selects an appropriate Tier to be used (Figure 3A) to document a given imaging experiment as determined by following the 4DN-BINA-OME tiered specifications (*36–38, 51*) and launches the *Manage Instrument* modality of Micro-Meta App by clicking the appropriate button (Figure 3B). Because Micro-Meta App was specifically designed to be tier-aware, Micro-Meta App automatically displays only metadata fields that are specified by 4DN-BINA-OME to belong to the tier that was selected upon launching the App (Figure 3A), thus massively reducing the documentation burden. In addition, to increase flexibility, the tier-level utilized for validation can be modified dynamically after opening the main Manage Instrument workspace. This way, the user can, for example, be presented with all Tier 2 appropriate fields while being required to only fill in Tier 1 fields for validation (see also point 3 ii).
2. Once entered in the *Manage Instrument* section, the user is given the option of selecting one of three different methods for managing an instrument (Figure 3C and D):

i. By selecting one of the two *Create from scratch* options (i.e., *Create Inverted from scratch* or *Create Upright from scratch*), the user is presented with a blank microscope canvas of the selected type to work with (Figure 3E).
ii. When selecting *Load from file,* the user is asked to select a pre-existing Microscope.JSON (example available on Zenodo as illustrated in Supplemental Material) (*57*) file from the local file system. Such Microscope.JSON files could, for example, be a template file that was created by microscope custodians at the local microscopy core facility, shared by a colleague, or downloaded from a repository for local use.
iii. Finally, a pre-existing Microscope.JSON file that has already been processed and saved to the local Micro-Meta App’s *Home Folder* can be loaded for further editing using the *Load from repository* modality. Here existing Microscope.JSON files are listed by Manufacturer and by Instrument Name to facilitate the selection of the appropriate file (Figure 3D).
3. Regardless of the chosen *Manage Instrument* modality, in the next step, the user is presented with the main instrument management workspace where they can perform the following actions:

i. In the top bar, the user can select a different tier-level for validation with respect to the one that had been selected upon entering the current Micro-Meta App session (Figure 3E). This feature allows the user to fill additional fields while not being required to provide all mandatory fields for a given tier level.
ii. By clicking the *Edit microscope* button (Figure 3E), the user can then enter attributes that refer to the instrument in general and that allow the description of the Microscope Stand the instrument is built upon. Upon exiting the *Edit Microscope* GUI, the application signals validation by changing the color of the dot on the button from red (i.e., incomplete) to green (i.e., complete and validated).
iii. By clicking the sidebar *Hardware Navigation* selection menus, the user can identify a given hardware component and drag it to the appropriate position on the instrument canvas, or to one of the top right generic “drawers” that can accommodate free-floating components that do not have a pre-existing position on the canvas (i.e., additional Objectives that do not fit in the objective turret). In the example depicted, a blank Objective component is selected from the *Magnification* menu (Figure 3E, [1]) and dragged to one of the objective-slots on the canvas, where it “snaps in place” upon release (Figure 3E, [2]).
iv. Once a component has been placed on the canvas, the icon is highlighted with a red dot to signal that the attributes have not been yet filled in and validated. Thus, the user is alerted about which microscope components need attention and can quickly identify which icon to work on. By clicking on the icon (Figure 3F, [3]), the user gains access to different tabs, each containing simple forms that display the required, tier-appropriate, metadata fields sanctioned by 4DN-BINA-OME (Figure 3F, [4]). To further increase usability, the *Confirm* button found at the bottom of each window can be used to automatically jump to mandatory fields that have to be filled before exiting. Upon completing all required fields, the icon is highlighted with a green dot making it easier for the user to assess documentation progress.
4. Once all appropriate microscope hardware components for a given instrument have been added to the canvas and appropriately attended to, the resulting tier-specific microscope *Hardware Specification* descriptions are then output as structured and interoperable Microscope.JSON files (example available on Zenodo as illustrated in Supplemental Material) (*57*). These files can be Saved locally (Figure 3G) or used by existing third-party databases, such as the 4DN Data Portal (*34, 58*) (Figure 6), for later utilization during the Manage Settings modality of the Micro-Meta App (see next section). In addition, such files can be imported in MethodsJ2 and used to automatically generate the Methods and Acknowledgement sections of scientific publications as described in a parallel manuscript (*50*). Finally, the same files could be associated with a Research Resource ID (RRID) (*59*) to acknowledge the work of microscope custodians, used by reviewers of manuscripts, shared with other users of the same microscope or with colleagues that need to document similar instrumentation, and broadly disseminated through appropriate data portals (*33, 34*).

**Figure 3.**
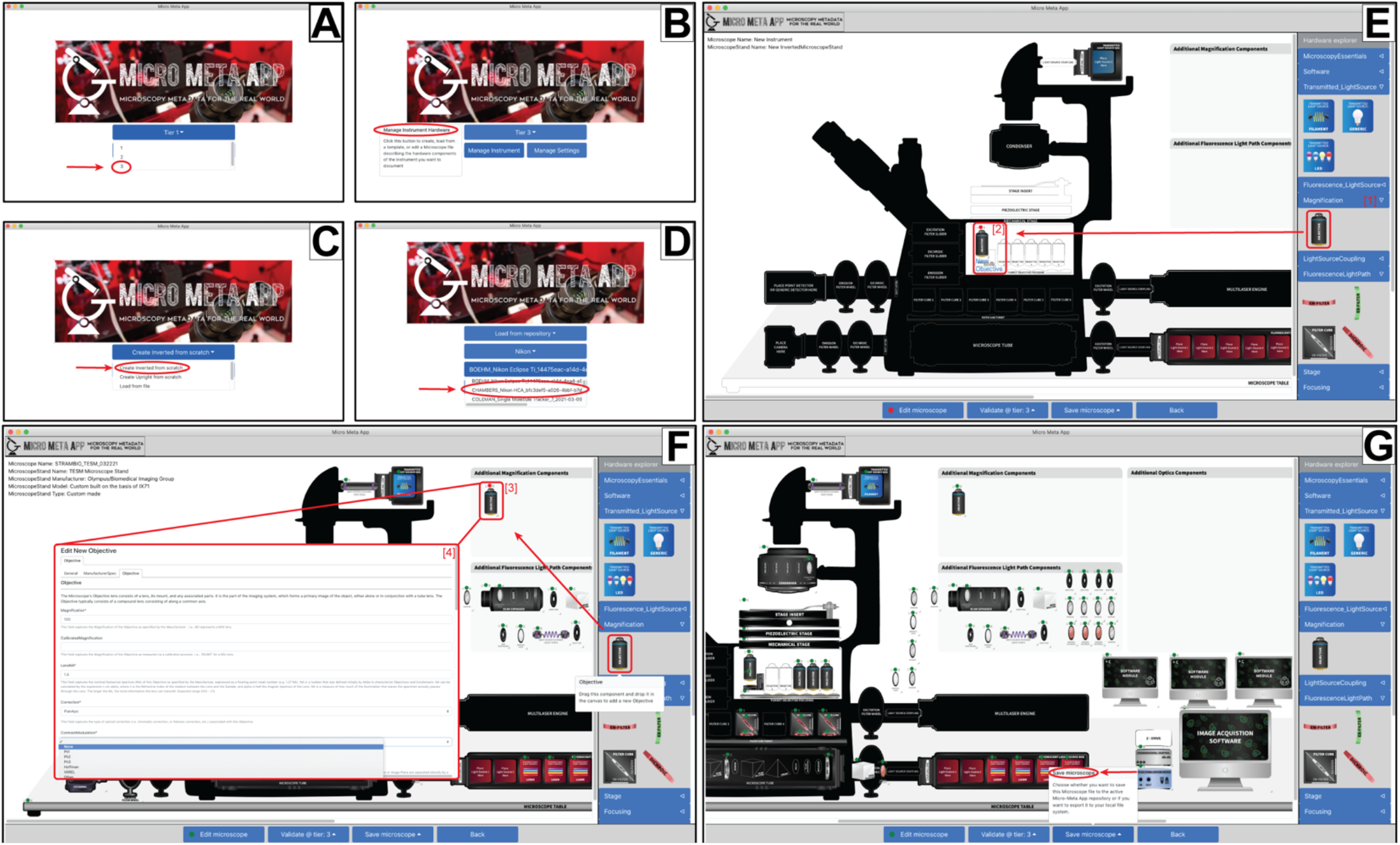
The Micro-Meta App is designed to aid in the collection of 4DN-BINA-OME sanctioned Microscope Hardware Specifications metadata. Micro-Meta App can be used as a stand-alone desktop application. Alternatively, it can be launched from a third-party portal such as the 4D Nucleome Data Portal (Figure 6), where Microscope files can be stored as image dataset attachments. In either situation: **A**) After selecting the desired location for saving *Microscope.JSON* files for later use (not shown), before creating a new microscope file it is first necessary to select the desired Tier for the experiment at hand. **B)** The *Manage Instrument* workflow in Micro-Meta App. Micro-Meta App can be used to create a structured *Microscope.JSON* file containing a description of the hardware components of a given microscopy Instrument based on the 4DN-BINA-OME ontology <*`Instrument`*> class model. **C)** A new microscope can be created totally from scratch by selecting either an Inverted or an Upright Microscope Stand. **D)** Alternatively, a previously saved *Microscope.JSON* file can be selected among those saved on an available repository and opened for further editing, such as in the depicted example. **E)** In this example, the *“Magnification”* drop-down menu is opened [1] and an *“Objective”* is dragged and snapped-in-place in the designated position [2] on the workspace. **F)** In this example, an additional *“Objective”* [3] is dragged onto the workspace and relevant metadata fields are entered using the designated metadata entry form [4]. **G)** Once all changes are entered, the microscope file can be saved to file or to the desired repository.

#### Manage Image Acquisition Settings

This modality is used to produce metadata-rich descriptions of the Image Acquisition Settings utilized to produce a given image dataset (Figure 4). In this modality, Micro-Meta App: 1) imports an existing Microscope.JSON file describing the instrument used to acquire the image data to be documented; 2) utilizes BioFormats (*43*) to read available OME-specified microscopy metadata stored in the header of the desired image file by the manufacturer of the microscope or of the acquisition software (Supplemental Tables I and II); 3) interactively guides the user through the collection of the missing, instrument-specific, tier-appropriate, 4DN-BINA-OME sanctioned Image Acquisition Settings utilized to produce the selected image data; and 4) produces structured Settings.JSON files (example available on Zenodo as illustrated in Supplemental Material) (*57*) that can be stored locally or in appropriate data portals alongside the images and the appropriate Microscope.JSON file. More in detail, the Manage Settings modality of Micro-Meta App articulates around the following steps:

1. After selecting a documentation tier (Figure 3A) and launching the App in the Manage Settings modality (Figure 4A), the user selects an available Microscope.JSON file from the local file system or a suitable repository (Figure 4B), selects an available image dataset to be annotated (Figure 4C) and either creates a new Settings.JSON file or opens an existing file to edit (Figure 4D). The integration of the Bio-Formats API (*43*) as part of Micro-Meta App permits the App to interpret image file headers, extract available metadata and populate Instrument-specific, OME-compatible, tier-appropriate metadata fields to facilitate metadata annotation.
2. As a result of the previous step, a diagrammatic representation of *Image Acquisition Settings* is displayed to the user and components or fields containing missing metadata values are highlighted in red to solicit the user’s attention (Figure 4E). After attending to all missing values, the user can then produce a validated 4DN-BINA-OME-compatible and tier-appropriate Settings.JSON file, which coupled with the corresponding Microscope.JSON file can be associated with the relevant image datasets in the local file system or on an appropriate repository, such as the 4DN Data Portal (*34, 52*). The Manage Settings section of the App consists of four types of user interaction interfaces:

i. Simple buttons with associated tabbed data entry forms, such as those that allow inspection, editing or entry of general information about the image Pixel structure (Figure 4E, [1.1 and [1.2]).
ii. Interface for the selection of one of the available hardware components, addition to the Settings.JSON file and editing of associated settings metadata fields. This type of user interface is used for *Edit Objective Settings* (Figure 4E, [3.1], [3.2] and [3.3]) and is also used for *Edit Imaging Environment, Edit Microscope Table Settings, Edit Microscope Stand Settings* and *Edit Sample Positioning Settings.*
iii. Specialized Plane-management interface (Figure 4E, [2.1], [2.2] and [2.3]). This interface is used either to inspect and, if necessary, edit the automatically imported *Planes* metadata or to record such metadata in case none was available in the header of the image data file to be annotated.
iv. Specialized Channel-management interface. Special attention was dedicated to the development of the GUI utilized to define the configuration and settings of the Light Path (i.e., Light Source → FilterSet → Detector) associated with each individual Channel (Figure 4F and G). To this aim, an intuitive Channel GUI (Figure 4F, [4.1] and [4.2]) is organized graphically around a visual representation of the Fluorescence Light Path where users can select among different light sources, filters, and detectors available in the underlying Microscope.JSON file and provide the appropriate settings to configure a given Channel. For example, the user would first select a given Light Source among those available (Figure 4F, [6.1]), and then enter the appropriate Light Source Settings (Figure 4F, [6.2]). The same Channel-specific interface can also be used to manage advanced Light Path features, such as in cases in which a custom-developed microscope has to be described (Figure 4G).
3. Once all components have been selected and configured, the Image Acquisition Settings are compiled in a structured Settings.JSON file (example available on Zenodo as illustrated in Supplemental Material) (*57*) and saved either locally or remotely as desired.

**Figure 4.**
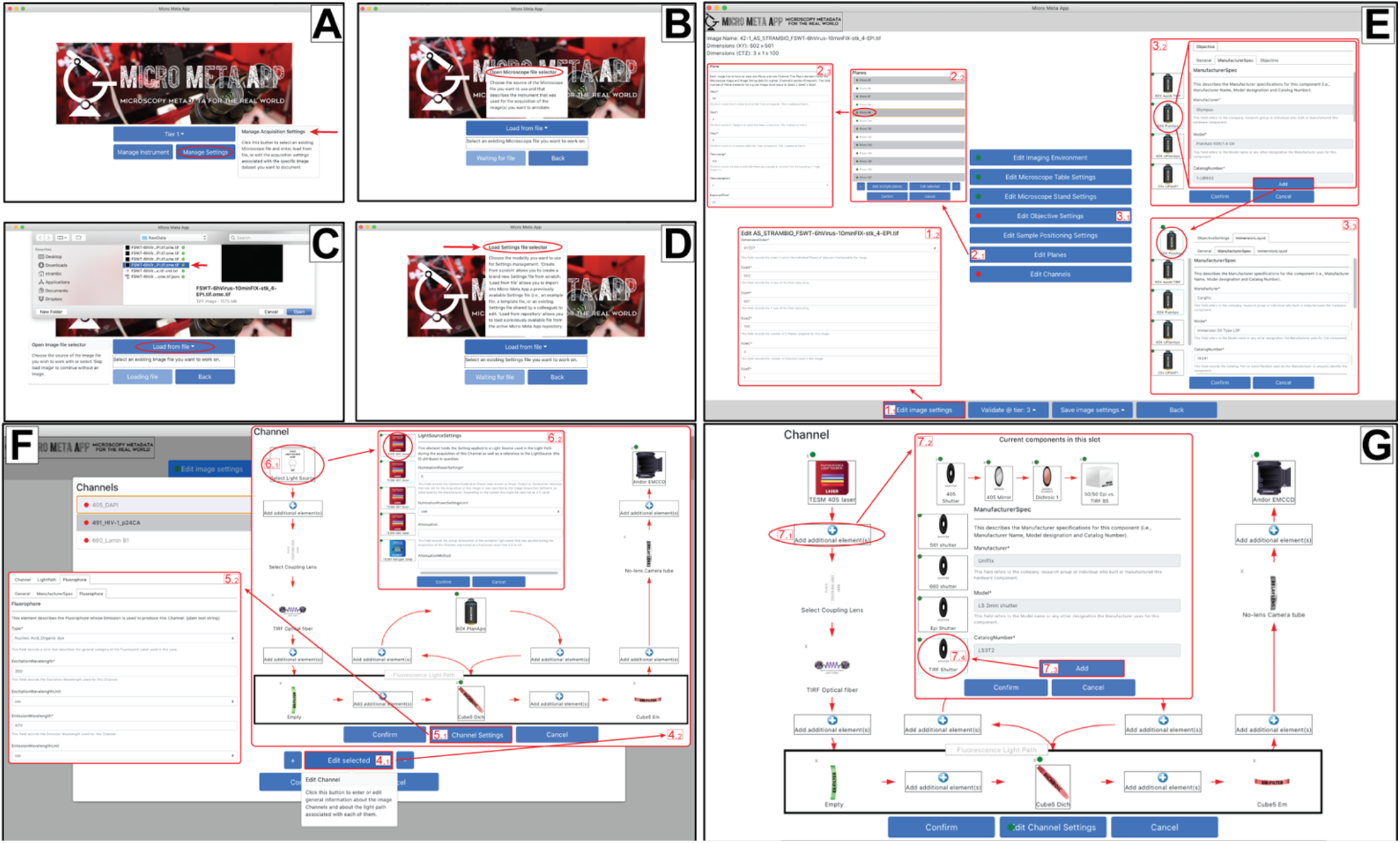
The Micro-Meta App utilizes the information contained in a previously generated Microscope metadata file and interactively aids in the collection of 4DN-BINA-OME sanctioned Image Acquisition settings metadata. Once a Microscope metadata file is created and saved (Figure 4), it can be used as the basis for collecting the Image Acquisition Settings utilized for a specific microscopy experiment. **(A-D)** After clicking on the *Manage Settings* button **(A)**, the user first selects the Microscope metadata file describing the instrument used to acquire the image of interest **(B)**, then the image file **(C)**, and finally, if available, an existing Settings metadata file **(D)**. From these files Micro-Meta App imports all relevant metadata for the user to inspect, edit and when missing enter from scratch. Specifically, Micro-Meta App allows to select the individual Microscope hardware components that were used to acquire a specific image and enter settings associated with each component. **E)** In the main window of *Manage Settings* the user can access different sections of the Image Acquisition Settings metadata by pressing different buttons and launching of the corresponding metadata collection windows. In this example, clicking the Edit image settings button [1.1] opens the metadata entry form for general image structure metadata associated with the image Pixels (e.g., DimensionOrder, SizeX, SizeY, SizeZ) allowing to inspect and if needed edit these values [1.2]. In addition, the Edit Planes button [2.1] opens an interface where the list of available image Planes is displayed [2.2], and individual Planes can be selected, so that associated metadata (e.g., TimeStamp, ExposureTime) can be inspected and edited [2.3]. Finally, the Edit Objective Settings button [3.1] allows to select the Objective that was used to acquire the image of interest, among those available in the Microscope file [3.2] and enter the relevant Objective Settings [3.3]. The same procedure followed for Objective Settings, can be also used to edit Imaging Environment, Microscope Table, Microscope Stand, and Sample Positioning Settings (not shown). **F)** After clicking the *Edit Channels* button in the main *Manage Settings* window (Panel E), the user opens an interface where the list of image Channels that were found in the image file header are displayed (top left) to be individually accessed and edited (button 4.1). The associated Channel interface [4.2] presents a a button called *Edit Channel Settings* [5.1] that launches a specialized window [5.2] to enter general information about the Channel (i.e., IlluminationType, ContrastMethod, and, when relevant, Fluorophore). In addition, the Channel window presents an interactive user interface for managing the different components of the channel’s Light Path (i.e., *LightSource, Fluorescence Light Path, Objective, Detector*). In this example, the user clicks on the *Light Source* button [6.1] to select one of the available Light Sources present in the Microscope file, add it to the Light Path and enter the associated settings that were applied during image acquisition [6.2]. **G)** The Channel interface can be used to manage advanced features of the Light Path of individual channels. This is done by inserting additional optical elements at one of the seven *Add additional element(s*) [7.1] insert buttons found at key locations along the Light Path. In the example displayed, the insertion point located between the Light Source and the illumination port found at the back of the Microscope initially contains a Shutter, a Mirror, a Dichroic and a Beam Splitter [7.2], and the *Add* [7.3] button is used to append an additional Shutter [7.4].

### Beta testing

Micro-Meta App was developed in the context of community efforts organized around the 4DN Consortium need for imaging data dissemination and integration with omics datasets (*33, 34*) and the BINA (*35*) effort to improve rigor quality control and reproducibility in light microscopy. As part of this effort, several core facilities were identified to serve as reference beta-testing sites for Micro-Meta App (Supplemental Table III). To this aim, the stand-alone JavaScript Electron implementation of Micro-Meta App was locally deployed and microscope custodians at individual beta testing sites were trained both on the use of the App and on bug and feature request reporting. Such feedback was collected either directly or by taking advantage of the GitHub issue-reporting feature and incorporated into the main development branch in a close-iterative cycle ahead of the release of the initial production version of the software.

### Case Studies

#### Utilization at core facilities

In response to significant interest we observed in the community and after beta testing, microscope custodians at several light microscopy facilities (Supplemental Table III) volunteered to serve as case studies on the use of Micro-Meta App to document both microscope instrumentation and example published image datasets produced in microscopy platforms (*60–79*). The application was utilized by microscope custodians at 16 sites to document one of their microscopes alongside the settings utilized for published images based on the 4DN-BINA-OME model Core + Basic Extension (*37, 38, 51*) (Figure 5 and Supplemental Figures 2-16).

**Figure 5.**
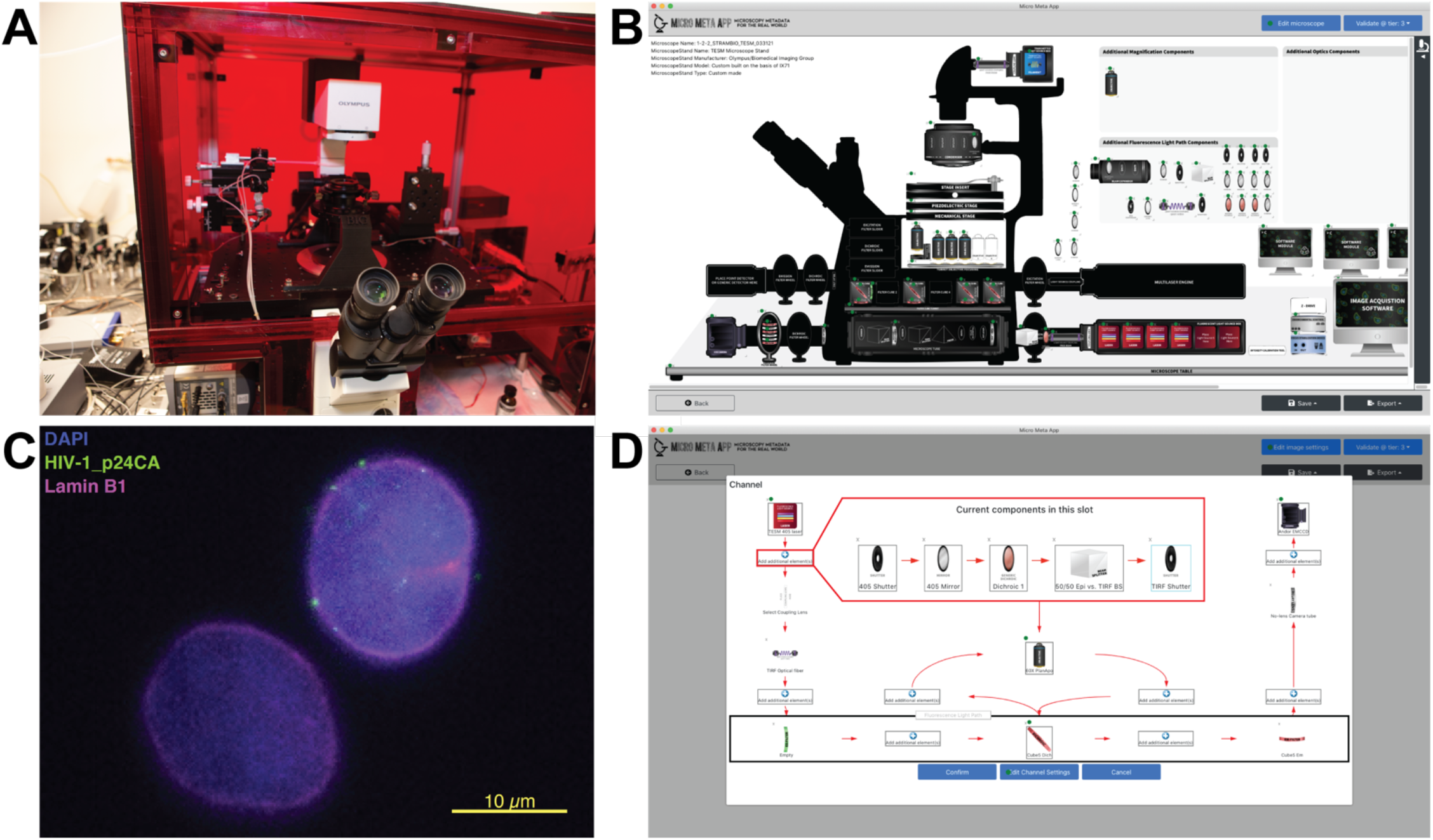
Micro-Meta App was utilized by several light microscopy core facilities to document individual microscope instrumentation and the settings that were applied to the microscope to acquire specific example datasets. Illustrated here is the use of Micro-Meta App for Tier 3 (*Manufacturing/Technical Development/Full Documentation*), 4DN-BINA-OME-specified (38) documentation of: **(A-B)** the custom build TIRF Epifluorescence Structured light Microscope (TESM; based on Olympus IX71) developed, built (75) and owned by the Biomedical Imaging Group at the Program in Molecular Medicine of the University of Massachusetts Medical School (UMMS; Supplemental Table III); and **(C-D)** the settings that were applied to the microscope for the acquisition of example images obtained by Nicholas Vecchietti and Caterina Strambio-De-Castillia. **A)** Picture of the indicated microscope. **B)** Micro-Meta App generated schematic representation of the indicated microscope. **C)** TZM-bl human cells were infected with HIV-1 retroviral three-part vector (FSWT+PAX2+pMD2.G). Six hours post-infection cells were fixed for 10 min with 1% formaldehyde in PBS, and permeabilized. Cells were stained with mouse anti-p24 primary antibody followed by DyLight488-anti-Mouse secondary antibody, to detect HIV-1 viral Capsid. In addition, cells were counterstained using rabbit anti-Lamin B1 primary antibody followed by DyLight649-anti-Rabbit secondary antibody, to visualize the nuclear envelope and with DAPI to visualize the nuclear chromosomal DNA. Displayed is a representative image obtained using the indicated microscope (Panels A and B) by Nicholas Vecchietti and Caterina Strambio-De-Castillia. **D)** Micro-Meta App generated schematic representation of the light path associated with the DAPI channel utilized for the acquisition of the image in Panel C. **E)** The insert shows a detail of the optical path followed by the 405-laser beam, in which the TIRF light path was used for epifluorescence illumination by orienting the beam perpendicular to the specimen plane. Specifically, in this portion of the light path the 405-laser beam passes through an electronically controlled shutter (*405 Shutter*), a mirror (*405 Mirror*), a dichroic filter (*Dichroic 1*), a 50/50 beam splitter (*50/50 Epi* vs. *TIRF BS*) and finally a shutter to control the TIRF light path illumination (*TIRF Shutter*). Example Microscope.JSON, Settings.JSON and associated image data file for this use case are available at: https://doi.org/10.5281/zenodo.4891883.

As indicated, the microscopes whose hardware was documented using Micro-Meta App comprised advanced custom-built microscopes, widefield microscopes and microscope stands associated with confocal systems, produced by all four major manufacturers. As a further testament of the robustness of the approach, several different major categories of imaging experiments were covered in this case study including:

1. Immunofluorescence imaging of the three-dimensional distribution of HIV-1 retroviral particles in the nucleus of infected human cells (Figure 5; example Microscope- and Settings.JSON files for this use case are available on Zenodo as illustrated in Supplemental Material) (*57*).
2. Three-dimensional visualization of superhydrophobic polymer-nanoparticles (Supplemental Figure 2).
3. Immunofluorescence imaging of cryosection of Mouse kidney (Supplemental Figure 4).
4. Three-dimensional immunofluorescence imaging of the phagocytic activity of human rhinovirus 16 (HRV16) infected macrophages (Supplemental Figure 7).
5. Live-cell imaging of *N. benthamiana* leaves cells-derived protoplasts transiently expressing YFP-tagged *P. chromatophora* proteins (Supplemental Figure 9).
6. Single-particle tracking of Halo-tagged PCNA in Lox cells (Supplemental Figure 12).
7. Live-cell imaging of bacterial cells expressing PopZ tagged with super-folder-GFP (Supplemental Figure 15).
8. Transmitted light brightfield visualization of swimming spermatocytes (Supplemental Figure 16).

In the case of commercial microscopes and given the type of experimental question and imaging modality, the most appropriate reporting tier-level was 4DN-BINA-OME Tier 2 (*38*). The exception was represented by one case in which no quantitative analysis was necessary, for which Tier 1 was sufficient (Supplemental Figure 2). On the other hand, two custom-built systems were documented at Tier 3 (Figure 5 and Supplemental Figure 16), again as sanctioned by the 4DN-BINA specifications (*38*).

The most striking result of these use cases was that in comparison with the metadata reporting baseline represented by the metadata fields made available using BioFormats alone, the use of Micro-Meta App significantly increased the uniformity of reported metadata fields. This facilitates the comparison of image data within and across different microscopes and imaging experiments. In addition, because the underlying data model utilized by Micro-Meta App is dynamically defined on the basis of shared guidelines that can evolve depending on needs and technological development, the use of this documentation method maximized reproducibility, quality and value and minimized effort on the part of individual scientists. The example set of Microscope- and Settings.JSON files produced for the use case illustrated in Figure 5, alongside the associated image file are available on Zenodo as illustrated in Supplemental Material (*57*).

#### Integration to 4DN Data Portal

One of the initial impetus for the development of Micro-Meta App was the need to expedite and, when possible, automate the rigorous reporting of imaging experiments and quality control procedures. In this context, Micro-Meta App was developed to be directly integrated into the 4DN Data Portal (*34, 58*). For this use, the App’s data flow was adapted to allow the direct integration of the content of the Microscope- and Settings-JSON files into the portal database. This allows individual fields to be utilized for filtering and searching purposes and to be visualized directly on the portal (Figure 6). In addition to maximizing flexibility, interfaces were developed to allow the import of pre-existing files the user might have produced using the desktop version of the App.

**Figure 6.**
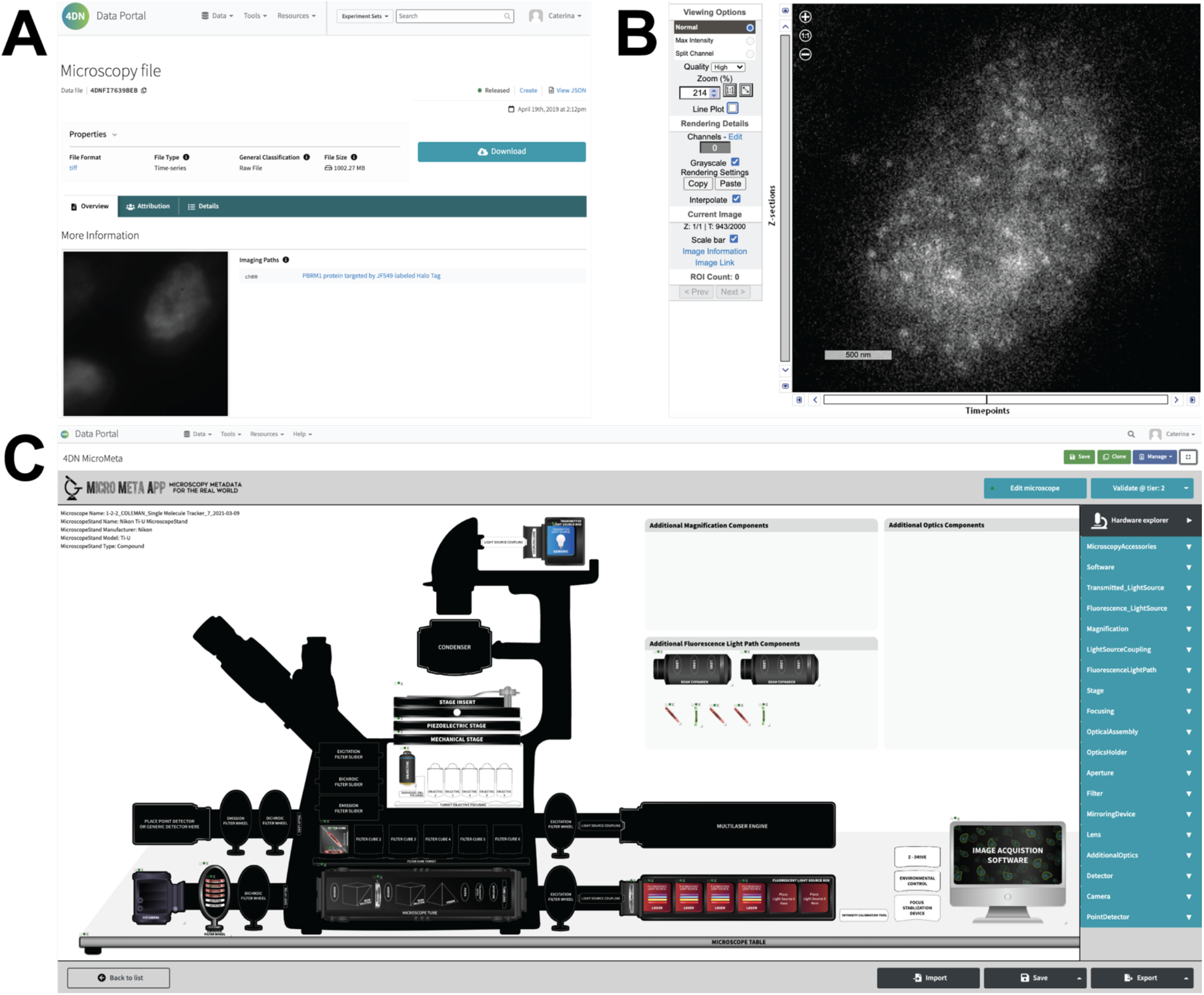
Micro-Meta App is integrated in the 4DN Data Portal for the documentation of imaging experiments. In order to facilitate the full documentation of microscopy experiments **(A)**, guarantee image data **(B)** quality and reproducibility and facilitate their integration with genomic data, Micro-Meta App **(C)** was incorporated in the 4DN Data Portal and made an integral part of the imaging experiment documentation and quality control workflow developed by the 4DN Data Coordination and Integration Center of the NIH-funded 4D Nucleome Consortium (34, 52, 57). For this use the App’s data flow was adapted to allow the direct ingestion of the content of the JSON Microscope- and Settings-files into the portal database, which allows individual fields to be utilized for filtering and searching purposes and to be visualized directly on the portal.

#### Micro-Meta App, microscopy platforms and teaching

Micro-Meta App provides a digital representation of freely configurable microscopes, ideal for microscopy platform staff to provide users with a detailed inventory of all available microscopes and a property that makes it perfectly suited for teaching purposes (Supplemental Video 1). Two major teaching use cases have been explored in the context of the Foundations in Biomedical Science (BBS 614) course (https://www.umassmed.edu/gsbs/academics/courses/) administered to first-year students at the Graduate School of Biomedical Sciences at the University of Massachusetts Medical school: 1) Micro-Meta App was used for students to work on specific problem sets; 2) Micro-Meta App was used for self-driven exploration of microscope components, functions and imaging modalities. In both cases, it is advisable to create specific teaching Microscope.JSON files that students can load and work on. Specifically, the features and complexity of these teaching Microscope.JSON files need to be aligned with course level and content by choosing the most appropriate tier-level among those available and, if necessary, by structuring the file without adhering to any one specific tier (*36, 80*). For example, a problem set might be assigned that focuses on choosing the most appropriate filter set for a given imaging experiment. Specifically, students are instructed to choose an appropriate light source and then specify each filter to be associated with the filter set. In this case, the depth of information associated with Tier 3 might be needed for the filters and a short list of possible light sources might be provided for the students to choose from (e.g., laser combiner with the wrong laser lines for the experiment and broadband source). At the same time the rest of the Microscope.JSON file could be kept at a very basic level to reduce grading complexity. In another example, a course might start on Day 1 with a Microscope.JSON file that only has a few components at Tier 1. On subsequent days, more components might be added, and the tier-level might be raised up to Tier 3 depending on the specific course teaching goals.

In addition to the tested use-cases, Micro-Meta App might be used in teaching modules available online (e.g., core facility user training) or used for flipped-classroom settings, where Micro-Meta App would be used for the practical application of microscopy concepts learned individually. Specifically, Micro-Meta App might be used to create snapshots of microscopes or as a guided sequence for familiarizing the user with the intricacies of specific instrument hardware configurations. As with any teaching material, alignment of the teaching microscope in Micro-Meta App with course content, intended use and grading complexity is critical for success.

### Future Directions

We will work closely within the context of 4DN (*33, 34*), BINA (*35*), Global Bioimaging (*3, 81*) and QUAREP-LiMi’s WG7 on Metadata (https://quarep.org/working-groups/wg-7-metadata/) on the following fronts:

1. <u>Implementation of additional microscopy modalities</u>: The current version of Micro-Meta App implements the Core of the OME data model and the 4DN-BINA-OME Basic extension (*38, 39*). One of the most urgent next steps will be implementing the Confocal and Advance extensions and subsequently, in close collaboration with QUAREP-LiMi (*4, 40*) to implement the Calibration and Performance extension (*38, 39*).
2. <u>Instrument Performance and Calibration implementation</u>: The 4DN-BINA-OME Microscopy Metadata Specifications (*38, 39*) include a Calibration and Performance extension which is not currently implemented in Micro-Meta App. In close collaboration with several of the core WG of QUAREP-LiMi we will work to integrate Quality Control metadata (procedure description and output metrics) into Micro-Meta App. As a starting point, illumination and detector calibration metrics calculated using the open-hardware Meta-Max(*82*) calibration tool will be automatically imported and used to annotate imaging datasets.
3. <u>Further integration with MethodsJ2</u>: Future development will: 1) integrate the *Micro-Meta App* software metadata *Settings.JSON* file so that it can serve as a source of Microscopy Metadata for *MethodsJ2* (*50*). 2) Adapt *Micro-Meta App* to generate methods text directly so researchers can use their platform of choice.
4. <u>Further OMERO and OMERO.mde integration</u>: As part of the initial Micro-Meta App development endeavor, a partial-functionality, prototype OMERO-plugin version of the App was developed and integrated into the OMERO instance, which is available at the University of Massachusetts Medical School (UMMS) (*83*). In an effort to facilitate the wide adoption of Micro-Meta App by the imaging community, integration with OMERO will be completed, including the extracting experimental metadata as specified by extended specifications as developed using *OMERO.mde* (*49*) and saving 4DN-BINA-OME metadata (*38, 39*) as collections of key-value pairs associated with individual image datasets.
5. <u>Creation of Instrument and Hardware components databases</u>: While engaging with microscope hardware manufacturers to ensure the full automation of data provenance and quality control reporting for light microscopy, it will be necessary to further reduce the documentation burden imposed on bench-side scientists, therefore maximizing their adoption of community standards. To this purpose, we will work in the context of Bioimaging North America to develop sharable databases of Microscope.JSON files. This effort could be integrated with the RRID effort (*59*) and will have the added advantage of promoting the recognition of Microscope-configurations as an essential and quantifiable scientific output, therefore providing credit to the work of imaging scientists and, in particular, microscope custodians.
6. <u>Outreach and education effort</u>: To involve both microscope users, custodians and manufacturers and promote their adoption of Micro-Meta App. In the case of Manufacturers, Micro-Meta App provides them with an opportunity to produce pre-filled JSON files describing individual components and make them available to the community from individual sites (similar to Fiji plugin sites) and utilized to automate during the production of Microscope- and Settings.JSON files.
7. <u>FPbase spectra-viewer integration</u>: The customized spectra-viewer feature of FPbase (*84, 85*) will be integrated into Micro-Meta App so the users can produce interactive microscope-specific spectral representations of each optical configuration. Such representations can, for example, be used to calculate the excitation and collection efficiency of a given Channel/Fluorophore combination. Vice versa, the possibility of integrating Micro-Meta App-generated Microscope.JSON files into the FPbase microscope-configurations feature will be explored. The interoperability of these two tools will improve the capacity of core-facilities to provide realistic teaching scenarios thus facilitating user training.

## Conclusions

For this work to have a broad impact tools such as Micro Meta App are required so that biological scientists will be able to quickly adopt metadata recommendations and incorporate them in their everyday work without requiring an extensive time commitment and regardless of their imaging expertise. This means that tools will need to be developed, accessible and straightforward to use in order to be successful and implement the proposed guidelines and metadata standards broadly. Ultimately, this will lead to automating all aspects of the process utilized by members of the community to annotate and upload metadata-rich imaging datasets to both local repositories such as OMERO (*86*) and public repositories such as Image Data Resource (IDR) (*20*) or other public image archives (*22*). In addition to the development of community standards for microscopy, reaching this goal will entail the automated interpretation of metadata stored in image file headers, the development of a communitywide repository for the storage of metadata specifications for commercial and custom-made microscopy hardware commonly utilized by imaging laboratories, and the automated annotation of imaging datasets to be uploaded in imaging data portals or disseminated through other means.

## Abbreviation list

BINA: BioImaging North America;
4DN: 4D Nucleome;
FAIR: Findable Accessible Interoperable and Reproducible;
OME: Open Microscopy Environment;
QUAREP-LiMi: QUAlity Assessment and REProducibility for Instrument and Images in Light Microscopy

## Availability, requirements, resources, and documentation

- *Project name:* Micro-Meta App
- *Project home page:* https://github.com/WU-BIMAC/MicroMetaApp.github.io *Documentation:* https://micrometaapp-docs.readthedocs.io/en/latest/index.html
- *Executable available at:*

- Desktop application (Javascript Electron): https://github.com/WU-BIMAC/MicroMetaApp-Electron/releases/latest
- *Source code available at:*

- Desktop application (Javascript Electron): https://zenodo.org/record/4750765
- Data-portal integratable application (Javascript React): https://zenodo.org/record/4751438
- *Example Microscope.JSON, Settings.JSON and associated image data file (Figure 5 use case): https://doi.org/10.5281/zenodo.4891883*
- *Operating system(s):* MacOS and PC
- *Programming language:* Javascript and Java
- *Other requirements:* Java v1.8.0. In addition, see dependencies listed in Supplemental Material.
- *License:* GNU GPL v3 (https://www.gnu.org/licenses/gpl-3.0.html)

## Authors contributions

Author contributions categories utilized here were devised by the CRediT initiative (*87, 88*).

**Alessandro Rigano:** Conceptualization, Methodology, Software, Validation, Investigation; **Shannon Ehmsen:** Resources, Visualization; **Serkan Utku Ozturk:** Software; **Joel Ryan:** Validation, Resources, Data Curation; **Alexander Balashov:** Conceptualization, Methodology, Software; **Mathias Hammer:** Conceptualization, Validation; **Koray Kirli:** Validation; **Kevin Bellve’**: Resources; **Ulrike Boehm:** Validation, Resources, Data Curation, Writing – Review & Editing; **Claire M. Brown:** Validation, Resources, Data Curation, Writing – Review & Editing; **James Chambers:** Validation, Resources, Data Curation; **Andrea Cosolo:** Validation, Writing – Review & Editing; **Robert Coleman:** Validation, Resources, Data Curation; **Kevin Fogarty:** Resources; **Orestis Faklaris:** Validation, Resources, Data Curation, Writing – Review & Editing; **Thomas Guilbert:** Validation, Resources, Data Curation, Writing – Review & Editing; **Anna B. Hamacher:** Resources; **Michelle S. Itano:** Validation, Resources, Data Curation, Writing – Review & Editing; **Daniel P. Keeley:** Validation, Data Curation; **Susanne Kunis:** Resources; **Judith Lacoste:** Validation, Resources, Data Curation, Writing – Review & Editing; **Alex Laude:** Validation, Resources, Data Curation, Writing – Review & Editing; **Willa Ma:** Validation, Data Curation; **Marco Marcello:** Validation, Resources, Data Curation, Writing – Review & Editing; **Paula Montero-Llopis:** Validation, Resources, Data Curation, Writing – Review & Editing; **Glyn Nelson:** Validation, Resources, Data Curation, Writing – Review & Editing; **Roland Nitschke:** Validation, Resources, Data Curation, Writing – Review & Editing; **Jaime A. Pimentel**: Validation, Resources, Data Curation, Writing – Review & Editing; **Stefanie Weidtkamp-Peters:** Validation, Resources, Data Curation; **Peter Park:** Supervision, Project administration, Funding acquisition; **Burak Alver:** Conceptualization, Validation, Resources, Writing – Review & Editing, Supervision, Project administration, Funding acquisition; **David Grunwald:** Conceptualization, Methodology, Investigation, Resources, Writing – Review & Editing, Supervision, Project administration, Funding acquisition; **Caterina Strambio-De-Castillia:** Conceptualization, Methodology, Software, Validation, Investigation, Resources, Data Curation, Writing – Original Draft, Writing – Review & Editing, Visualization, Supervision, Project administration, Funding acquisition.

## Acknowledgments

We would like to thank Lawrence Lifshitz at the Biomedical Imaging Group of the Program in Molecular Medicine at the University of Massachusetts Medical School for invaluable intellectual input and countless fruitful discussions and for their friendship, advice, and steadfast support throughout the development of this project. We acknowledge Matteo Luban for critically reading and editing the manuscript.

This project could never have been carried out without the leadership, insightful discussions, support and friendship of all OME consortium members, with particular reference to Jason Swedlow, Josh Moore, Chris Allan, Jean Marie Burel, and Will Moore. We are massively indebted to the RIKEN community for their fantastic work to bring open science into biology. We would like to particularly acknowledge Norio Kobayashi and Shuichi Onami for their friendship and support.

We thank all members of Bioimaging North America, German Bioimaging, Euro-Bioimaging (in particular Antje Keppler and Federica Paina) and QUAREP-LiMi (in particular all members of the Working Group 7 – Metadata; quarep.edu) for invaluable intellectual input, fruitful discussions and advice. We are also indebted to the following individuals for their continued and steadfast support: Jeremy Luban, Roger Davis, and Thoru Pederson at the University of Massachusetts Medical School; Burak Alver, Joan Ritland, Rob Singer, and Warren Zipfel at the 4D Nucleome Project; Ian Fingerman, John Satterlee, Judy Mietz, Richard Conroy, and Olivier Blondel at the NIH.

This work was supported by NIH grant #1U01EB021238 and NSF grant 1917206 to D.G., NIH grant # U01CA200059 to C.S.D.C and D.G., NIH grant # 5R01GM126045-04 to R.C. and by grant #2019-198155 (5022) awarded to C.S.D.C. by the Chan Zuckerberg Initiative DAF, an advised fund of Silicon Valley Community Foundation, as part of their Imaging Scientist Program. D.S. was funded in part by NIH/NCI grants U54CA209988 and U2CCA23380. C.M.B. was funded in part by grant #2020-225398 from the Chan Zuckerberg Initiative DAF, an advised fund of Silicon Valley Community Foundation. R.N. was funded by the Deutsche Forschungsgemeinschaft (DFG, German Research Foundation) grant number Ni 451/9-1 MIAP-Freiburg. C.S.S. was supported by the Netherlands Organisation for Scientific Research (NWO), under NWO START-UP project no. 740.018.015 and NWO Veni project no. 16761. T.G. is a member of RTmfm network and IMAG’IC core facility is supported by the National Infrastructure France BioImaging (grant ANR-10-INBS-04) and IBISA consortium. S.W.-P. was funded by the Deutsche Forschungsgemeinschaft (DFG, German Research Foundation) project-ID 267205415 – SFB 1208, project INF. The UNC Neuroscience Microscopy Core (RRID:SCR_019060) is supported, in part, by funding from the NIH-NINDS Neuroscience Center Support Grant P30 NS045892 and the NIH-NICHD Intellectual and Developmental Disabilities Research Center Support Grant P50 H103573. M.S.I. was supported by grant number 2019-198107, from the Chan Zuckerberg Initiative DAF, an advised fund of Silicon Valley Community Foundation.

## References

1. Nature Editorial Staff, Better research through metrology. Nat. Methods. 15, 395 (2018).

2. J. Pines, Image integrity and standards. Open Biol. 10(2020), p. 200165.

3. J. R. Swedlow, P. Kankaanpää, U. Sarkans, W. Goscinski, G. Galloway, R. P. Sullivan, C. M. Brown, C. Wood, A. Keppler, B. Loos, S. Zullino, D. L. Longo, S. Aime, S. Onami, A Global View of Standards for Open Image Data Formats and Repositories. arXiv [q-bio.OT] (2020), (available at http://arxiv.org/abs/2010.10107).

4. G. Nelson, U. Boehm, S. Bagley, P. Bajcsy, J. Bischof, C. M. Brown, A. Dauphin, I. M. Dobbie, J. E. Eriksson, O. Faklaris, J. Fernandez-Rodriguez, A. Ferrand, L. Gelman, A. Gheisari, H. Hartmann, C. Kukat, A. Laude, M. Mitkovski, S. Munck, A. J. North, T. M. Rasse, U. Resch-Genger, L. C. Schuetz, A. Seitz, C. Strambio-De-Castillia, J. R. Swedlow, I. Alexopoulos, K. Aumayr, S. Avilov, G.-J. Bakker, R. R. Bammann, A. Bassi, H. Beckert, S. Beer, Y. Belyaev, J. Bierwagen, K. A. Birngruber, M. Bosch, J. Breitlow, L. A. Cameron, J. Chalfoun, J. J. Chambers, C.-L. Chen, E. Conde-Sousa, A. D. Corbett, F. P. Cordelieres, E. Del Nery, R. Dietzel, F. Eismann, E. Fazeli, A. Felscher, H. Fried, N. Gaudreault, W. I. Goh, T. Guilbert, R. Hadleigh, P. Hemmerich, G. A. Holst, M. S. Itano, C. B. Jaffe, H. K. Jambor, S. C. Jarvis, A. Keppler, D. Kirchenbuechler, M. Kirchner, N. Kobayashi, G. Krens, S. Kunis, J. Lacoste, M. Marcello, G. G. Martins, D. J. Metcalf, C. A. Mitchell, J. Moore, T. Mueller, M. S. Nelson, S. Ogg, S. Onami, A. L. Palmer, P. Paul-Gilloteaux, J. A. Pimentel, L. Plantard, S. Podder, E. Rexhepaj, M. Royeck, A. Royon, M. A. Saari, D. Schapman, V. Schoonderwoert, B. Schroth-Diez, S. Schwartz, M. Shaw, M. Spitaler, M. T. Stoeckl, D. Sudar, J. Teillon, S. Terjung, R. Thuenauer, C. D. Wilms, G. D. Wright, R. Nitschke, QUAREP-LiMi: A community-driven initiative to establish guidelines for quality assessment and reproducibility for instruments and images in light microscopy. arXiv [q-bio.OT] (2021), (available at http://arxiv.org/abs/2101.09153).

5. G. Marqués, T. Pengo, M. A. Sanders, Imaging methods are vastly underreported in biomedical research. Elife. 9(2020), doi:10.7554/eLife.55133.

6. P. Montero Llopis, R. A. Senft, T. J. Ross-Elliott, R. Stephansky, D. P. Keeley, P. Kosha, G. Marqués, Y. Gao, B. R. Carlson, T. Pengo, M. A. Sanders, L. A. Cameron, M. S. Itano, Best practices and tools for reporting reproducible fluorescence microscopy methods. Nat. Methods. 35(2021), doi:10.1038/s41592-021-01156-w.

7. S. Ram, J. Liu, A Semiotics Framework for Analyzing Data Provenance Research. Journal of Computing Science and Engineering. 2, 221–248 (2008).

8. S. Ram, J. Liu, A Semantic Foundation for Provenance Management. J. Data Semant. 1, 11–17 (2012).

9. J. M. Heddleston, J. S. Aaron, S. Khuon, T.-L. Chew, A guide to accurate reporting in digital image processing: can anyone reproduce your quantitative analysis? J. Cell Sci. 134: jcs254151(2021), doi:10.1242/jcs.254151.

10. J. S. Aaron, T.-L. Chew, A guide to accurate reporting in digital image acquisition: can anyone replicate your microscopy data? J. Cell Sci. 134: jcs254144(2021), doi:10.1242/jcs.254144.

11. M. R. Sheen, J. L. Fields, B. Northan, J. Lacoste, L.-H. Ang, S. Fiering, Reproducibility Project: Cancer Biology, Replication Study: Biomechanical remodeling of the microenvironment by stromal caveolin-1 favors tumor invasion and metastasis. Elife. 8(2019), doi:10.7554/eLife.45120.

12. M. P. Viana, J. Chen, T. A. Knijnenburg, R. Vasan, C. Yan, J. E. Arakaki, M. Bailey, B. Berry, A. Borensztejn, J. M. Brown, S. Carlson, J. A. Cass, B. Chaudhuri, K. R. Cordes Metzler, M. E. Coston, Z. J. Crabtree, S. Davidson, C. M. DeLizo, S. Dhaka, S. Q. Dinh, T. P. Do, J. Domingus, R. M. Donovan-Maiye, T. J. Foster, C. L. Frick, G. Fujioka, M. A. Fuqua, J. L. Gehring, K. A. Gerbin, T. Grancharova, B. W. Gregor, L. J. Harrylock, A. Haupt, M. C. Hendershott, C. Hookway, A. R. Horwitz, C. Hughes, E. J. Isaac, G. R. Johnson, B. Kim, A. N. Leonard, W. W. Leung, J. J. Lucas, S. A. Ludmann, B. M. Lyons, H. Malik, R. McGregor, G. E. Medrash, S. L. Meharry, K. Mitcham, I. A. Mueller, T. L. Murphy-Stevens, A. Nath, A. M. Nelson, L. Paleologu, T. Alexander Popiel, M. M. Riel-Mehan, B. Roberts, L. M. Schaefbauer, M. Schwarzl, J. Sherman, S. Slaton, M. Filip Sluzewski, J. E. Smith, Y. Sul, M. J. Swain-Bowden, W. Joyce Tang, D. J. Thirstrup, D. M. Toloudis, A. P. Tucker, V. Valencia, W. Wiegraebe, T. Wijeratna, R. Yang, R. J. Zaunbrecher, Allen Institute for Cell Science, G. T. Johnson, R. N. Gunawardane, N. Gaudreault, J. A. Theriot, S. M. Rafelski, Robust integrated intracellular organization of the human iPS cell: where, how much, and how variable. BioRxiv.org (2021), p. 2020.12.08.415562.

13. R. Botvinik-Nezer, F. Holzmeister, C. F. Camerer, A. Dreber, J. Huber, M. Johannesson, M. Kirchler, R. Iwanir, J. A. Mumford, R. A. Adcock, P. Avesani, B. M. Baczkowski, A. Bajracharya, L. Bakst, S. Ball, M. Barilari, N. Bault, D. Beaton, J. Beitner, R. G. Benoit, R. M. W. J. Berkers, J. P. Bhanji, B. B. Biswal, S. Bobadilla-Suarez, T. Bortolini, K. L. Bottenhorn, A. Bowring, S. Braem, H. R. Brooks, E. G. Brudner, C. B. Calderon, J. A. Camilleri, J. J. Castrellon, L. Cecchetti, E. C. Cieslik, Z. J. Cole, O. Collignon, R. W. Cox, W. A. Cunningham, S. Czoschke, K. Dadi, C. P. Davis, A. D. Luca, M. R. Delgado, L. Demetriou, J. B. Dennison, X. Di, E. W. Dickie, E. Dobryakova, C. L. Donnat, J. Dukart, N. W. Duncan, J. Durnez, A. Eed, S. B. Eickhoff, A. Erhart, L. Fontanesi, G. M. Fricke, S. Fu, A. Galván, R. Gau, S. Genon, T. Glatard, E. Glerean, J. J. Goeman, S. A. E. Golowin, C. González-García, K. J. Gorgolewski, C. L. Grady, M. A. Green, J. F. Guassi Moreira, O. Guest, S. Hakimi, J. P. Hamilton, R. Hancock, G. Handjaras, B. B. Harry, C. Hawco, P. Herholz, G. Herman, S. Heunis, F. Hoffstaedter, J. Hogeveen, S. Holmes, C.-P. Hu, S. A. Huettel, M. E. Hughes, V. Iacovella, A. D. Iordan, P. M. Isager, A. I. Isik, A. Jahn, M. R. Johnson, T. Johnstone, M. J. E. Joseph, A. C. Juliano, J. W. Kable, M. Kassinopoulos, C. Koba, X.-Z. Kong, T. R. Koscik, N. E. Kucukboyaci, B. A. Kuhl, S. Kupek, A. R. Laird, C. Lamm, R. Langner, N. Lauharatanahirun, H. Lee, S. Lee, A. Leemans, A. Leo, E. Lesage, F. Li, M. Y. C. Li, P. C. Lim, E. N. Lintz, S. W. Liphardt, A. B. Losecaat Vermeer, B. C. Love, M. L. Mack, N. Malpica, T. Marins, C. Maumet, K. McDonald, J. T. McGuire, H. Melero, A. S. Méndez Leal, B. Meyer, K. N. Meyer, G. Mihai, G. D. Mitsis, J. Moll, D. M. Nielson, G. Nilsonne, M. P. Notter, E. Olivetti, A. I. Onicas, P. Papale, K. R. Patil, J. E. Peelle, A. Pérez, D. Pischedda, J.-B. Poline, Y. Prystauka, S. Ray, P. A. Reuter-Lorenz, R. C. Reynolds, E. Ricciardi, J. R. Rieck, A. M. Rodriguez-Thompson, A. Romyn, T. Salo, G. R. Samanez-Larkin, E. Sanz-Morales, M. L. Schlichting, D. H. Schultz, Q. Shen, M. A. Sheridan, J. A. Silvers, K. Skagerlund, A. Smith, D. V. Smith, P. Sokol-Hessner, S. R. Steinkamp, S. M. Tashjian, B. Thirion, J. N. Thorp, G. Tinghög, L. Tisdall, S. H. Tompson, C. Toro-Serey, J. J. Torre Tresols, L. Tozzi, V. Truong, L. Turella, A. E. van ‘t Veer, T. Verguts, J. M. Vettel, S. Vijayarajah, K. Vo, M. B. Wall, W. D. Weeda, S. Weis, D. J. White, D. Wisniewski, A. Xifra-Porxas, E. A. Yearling, S. Yoon, R. Yuan, K. S. L. Yuen, L. Zhang, X. Zhang, J. E. Zosky, T. E. Nichols, R. A. Poldrack, T. Schonberg, Variability in the analysis of a single neuroimaging dataset by many teams. Nature. 582, 84–88 (2020).

14. U. Sarkans, W. Chiu, L. Collinson, M. C. Darrow, J. Ellenberg, D. Grunwald, J.-K. Hériché, A. Iuding, G. G. Martins, T. Meehan, K. Narayan, A. Patwardhan, M. Robert, G. Russell, H. R. Saibil, C. Strambio-De-Castillia, J. R. Swedlow, C. Tischer, V. Uhlmann, P. Verkade, M. Barlow, O. Bayraktar, E. Birney, C. Catavitello, C. Cawthorne, S. Wagner-Conrad, E. Duke, P. Paul-Gilloteaux, E. Gustin, M. Harkiolaki, P. Kankaanpää, T. Lemberger, J. McEntyre, J. Moore, A. W. Nicholls, S. Onami, H. Parkinson, M. Parsons, M. Romanchikova, N. Sofroniew, J. Swoger, N. Utz, L. Voortman, F. Wong, P. Zhang, G. J. Kleywegt, A. Brazma, REMBI: Recommended Metadata for Biological Images – realizing the full potential of the bioimaging revolution by enabling data reuse. Nat. Methods.ePub(2021, doi:10.1038/s41592-021-01166-8.

15. J. Rung, A. Brazma, Reuse of public genome-wide gene expression data. Nat. Rev. Genet. 14, 89–99 (2013).

16. A. Brazma, P. Hingamp, J. Quackenbush, G. Sherlock, P. Spellman, C. Stoeckert, J. Aach, W. Ansorge, C. A. Ball, H. C. Causton, T. Gaasterland, P. Glenisson, F. C. Holstege, I. F. Kim, V. Markowitz, J. C. Matese, H. Parkinson, A. Robinson, U. Sarkans, S. Schulze-Kremer, J. Stewart, R. Taylor, J. Vilo, M. Vingron, Minimum information about a microarray experiment (MIAME)-toward standards for microarray data. Nat. Genet. 29, 365–371 (2001).

17. J. P. A. Ioannidis, D. B. Allison, C. A. Ball, I. Coulibaly, X. Cui, A. C. Culhane, M. Falchi, C. Furlanello, L. Game, G. Jurman, J. Mangion, T. Mehta, M. Nitzberg, G. P. Page, E. Petretto, V. van Noort, Repeatability of published microarray gene expression analyses. Nat. Genet. 41, 149–155 (2009).

18. A. Brazma, Minimum Information About a Microarray Experiment (MIAME)--successes, failures, challenges. ScientificWorldJournal. 9, 420–423 (2009).

19. S.-A. Sansone, P. Rocca-Serra, M. Brandizi, A. Brazma, D. Field, J. Fostel, A. G. Garrow, J. Gilbert, F. Goodsaid, N. Hardy, P. Jones, A. Lister, M. Miller, N. Morrison, T. Rayner, N. Sklyar, C. Taylor, W. Tong, G. Warner, S. Wiemann, The First RSBI (ISA-TAB) Workshop: “Can a Simple Format Work for Complex Studies?”OMICS. 12, 143–149 (2008).

20. E. Williams, J. Moore, S. W. Li, G. Rustici, A. Tarkowska, A. Chessel, S. Leo, B. Antal, R. K. Ferguson, U. Sarkans, A. Brazma, R. E. C. Salas, J. R. Swedlow, The Image Data Resource: A Bioimage Data Integration and Publication Platform. Nat. Methods. 14, 775–781 (2017).

21. A. Iudin, P. K. Korir, J. Salavert-Torres, G. J. Kleywegt, A. Patwardhan, EMPIAR: a public archive for raw electron microscopy image data. Nat. Methods. 13, 387–388 (2016).

22. J. Ellenberg, J. R. Swedlow, M. Barlow, C. E. Cook, U. Sarkans, A. Patwardhan, A. Brazma, E. Birney, A call for public archives for biological image data. Nat. Methods. 15, 849–854 (2018).

23. Movincell Consortium, Multi-dimensional marine organism dataview. Movincell (2015), (available at http://movincell.org/).

24. Y. Tohsato, K. H. L. Ho, K. Kyoda, S. Onami, SSBD: a database of quantitative data of spatiotemporal dynamics of biological phenomena. Bioinformatics. 32, 3471–3479 (2016).

25. D. N. Orloff, J. H. Iwasa, M. E. Martone, M. H. Ellisman, C. M. Kane, The cell: an image library-CCDB: a curated repository of microscopy data. Nucleic Acids Res. 41, D1241–50 (2013).

26. M. Ellisman, S. Peltier, D. Orloff, W. Willy Wong, S. Penticoff, Center for Research in Biological Systems, Cell Image Library. www.cellimagelibrary.org (2019), (available at http://www.cellimagelibrary.org/home).

27. DORY Working Group, Defining Our Research Methodology. https://doryworkspace.org/ (2019), (available at https://doryworkspace.org/).

28. Allen Institute for Cell Science, Allen Institute for Cell Science. Allen Cell Explorer. alleninstitute.org (2017), (available at https://www.allencell.org/).

29. O. Rozenblatt-Rosen, J. W. Shin, J. E. Rood, A. Hupalowska, Human Cell Atlas Standards and Technology Working Group, A. Regev, H. Heyn, Building a high-quality Human Cell Atlas. Nat. Biotechnol. 39, 149–153 (2021).

30. R. G. H. Lindeboom, A. Regev, S. A. Teichmann, Towards a Human Cell Atlas: Taking Notes from the Past. Trends Genet. (2021), doi:10.1016/j.tig.2021.03.007.

31. O. Rozenblatt-Rosen, M. J. T. Stubbington, A. Regev, S. A. Teichmann, The Human Cell Atlas: from vision to reality. Nature. 550, 451–453 (2017).

32. A. Regev, S. A. Teichmann, E. S. Lander, I. Amit, C. Benoist, E. Birney, B. Bodenmiller, P. Campbell, P. Carninci, M. Clatworthy, H. Clevers, B. Deplancke, I. Dunham, J. Eberwine, R. Eils, W. Enard, A. Farmer, L. Fugger, B. Göttgens, N. Hacohen, M. Haniffa, M. Hemberg, S. Kim, P. Klenerman, A. Kriegstein, E. Lein, S. Linnarsson, E. Lundberg, J. Lundeberg, P. Majumder, J. C. Marioni, M. Merad, M. Mhlanga, M. Nawijn, M. Netea, G. Nolan, D. Pe’er, A. Phillipakis, C. P. Ponting, S. Quake, W. Reik, O. Rozenblatt-Rosen, J. Sanes, R. Satija, T. N. Schumacher, A. Shalek, E. Shapiro, P. Sharma, J. W. Shin, O. Stegle, M. Stratton, M. J. T. Stubbington, F. J. Theis, M. Uhlen, A. van Oudenaarden, A. Wagner, F. Watt, J. Weissman, B. Wold, R. Xavier, N. Yosef, Human Cell Atlas Meeting Participants, The Human Cell Atlas. Elife. 6(2017), doi:10.7554/eLife.27041.

33. 4D Nucleome Consortium, The 4D Nucleome Web Portal. 4dnucleome.org (2017), (available at https://www.4dnucleome.org/).

34. J. Dekker, A. S. Belmont, M. Guttman, V. O. Leshyk, J. T. Lis, S. Lomvardas, L. A. Mirny, C. C. O’Shea, P. J. Park, B. Ren, J. C. R. Politz, J. Shendure, S. Zhong, 4D Nucleome Network, The 4D nucleome project. Nature. 549, 219–226 (2017).

35. C. Strambio-De-Castillia, P. Bajcsy, U. Boehm, J. Chambers, A. D. Corbett, O. Faklaris, N. Gaudreault, J. Lacoste, A. Laude, G. Nelson, R. Nitschke, J. A. Pimentel, D. Sudar, C. M. Brown, A. J. North, Quality Control and Data Management | Bioimaging North America (BINA). Bioimaging North America (2019), (available at https://www.bioimagingna.org/qc-dm-wg).

36. M. Huisman, M. Hammer, A. Rigano, F. Farzam, R. Gopinathan, C. Smith, D. Grunwald, C. Strambio-De-Castillia, Minimum Information guidelines for fluorescence microscopy: increasing the value, quality, and fidelity of image data. arXiv [q-bio.QM] (2019), (available at http://arxiv.org/abs/1910.11370).

37. M. Huisman, M. Hammer, A. Rigano, U. Boehm, J. J. Chambers, N. Gaudreault, J. A. Pimentel, D. Sudar, P. Bajcsy, C. M. Brown, A. D. Corbett, O. Faklaris, J. Lacoste, A. Laude, G. Nelson, R. Nitschke, A. J. North, D. Grunwald, C. Strambio-De-Castillia, A perspective on Microscopy Metadata: data provenance and quality control. arXiv [q-bio.QM] (2021), (available at https://arxiv.org/abs/1910.11370).

38. M. Hammer, M. Huisman, A. Rigano, U. Boehm, J. J. Chambers, N. Gaudreault, J. A. Pimentel, D. Sudar, P. Bajcsy, C. M. Brown, A. D. Corbett, O. Faklaris, J. Lacoste, A. Laude, G. Nelson, R. Nitschke, A. J. North, R. Gopinathan, F. Farzam, C. Smith, D. Grunwald, C. Strambio-De-Castillia, Towards community-driven metadata standards for light microscopy: tiered specifications extending the OME model. BioRxiv (2021), doi:10.1101/2021.04.25.441198v1.

39. A. Rigano, U. Boehm, J. J. Chambers, N. Gaudreault, A. J. North, J. A. Pimentel, D. Sudar, P. Bajcsy, C. M. Brown, A. D. Corbett, O. Faklaris, J. Lacoste, A. Laude, G. Nelson, R. Nitschke, D. Grunwald, C. Strambio-De-Castillia, 4DN-BINA-OME (NBO) Tiered Microscopy Metadata Specifications – v2.01 (https://github.com/WU-BIMAC, 2021; https://zenodo.org/record/4710731).

40. U. Boehm, G. Nelson, C. M. Brown, S. Bagley, P. Bajcsy, J. Bischof, A. Dauphin, I. M. Dobbie, J. E. Eriksson, O. Faklaris, J. Fernandez-Rodriguez, A. Ferrand, L. Gelman, A. Gheisari, H. Hartmann, C. Kukat, A. Laude, M. Mitkovski, S. Munck, A. J. North, T. M. Rasse, U. Resch-Genger, L. C. Schuetz, A. Seitz, C. Strambio-De-Castillia, J. R. Swedlow, R. Nitschke, QUAREP-LiMi: A community endeavour to advance Quality Assessment and Reproducibility in Light Microscopy. Nat. Methods.ePub(2021), doi: 10.1038/s41592-021-01162-y.

41. I. G. Goldberg, C. Allan, J.-M. Burel, D. Creager, A. Falconi, H. Hochheiser, J. Johnston, J. Mellen, P. K. Sorger, J. R. Swedlow, The Open Microscopy Environment (OME) Data Model and XML file: open tools for informatics and quantitative analysis in biological imaging. Genome Biol. 6, R47 (2005).

42. J. R. Swedlow, I. G. Goldberg, E. Brauner, P. K. Sorger, Informatics and Quantitative Analysis in Biological Imaging. Science. 300, 100–102 (2003).

43. M. Linkert, C. T. Rueden, C. Allan, J.-M. Burel, W. Moore, A. Patterson, B. Loranger, J. Moore, C. Neves, D. MacDonald, A. Tarkowska, C. Sticco, E. Hill, M. Rossner, K. W. Eliceiri, J. R. Swedlow, Metadata matters: access to image data in the real world. J. Cell Biol. 189, 777–782 (2010).

44. J. Moore, C. Allan, S. Besson, J.-M. Burel, E. Diel, D. Gault, K. Kozlowski, D. Lindner, M. Linkert, T. Manz, W. Moore, C. Tischer, J. R. Swedlow, OME-NGFF: scalable format strategies for interoperable bioimaging data. bioRxiv (2021), p. 2021.03.31.437929.

45. D. P. W. Russell, P. K. Sorger, Maintaining the provenance of microscopy metadata using OMERO.forms software. BiorXiv, 109199 (2017).

46. J. Hay, E. Troup, I. Clark, J. Pietsch, T. Zieliński, A. Millar, PyOmeroUpload: A Python toolkit for uploading images and metadata to OMERO. Wellcome Open Res. 5, 96 (2020).

47. S. Kunis, OMERO.mde (Github, 2018; https://github.com/sukunis/omero-insight).

48. J. Moore, N. Kobayashi, S. Kunis, S. Onami, J. R. Swedlow, the OME Consortium, in Proceedings of the 12th SWAT4(HC)LS (Semantic Web Applications and Tools for Healthcare and Life Sciences) Conference, A. Burger, R. Cornet, A. Waagmeester, Eds. (http://ceur-ws.org, 2019), p. 17.

49. S. Kunis, S. Hänsch, C. Schmidt, F. Wong, S. Weidtkamp-Peters, OMERO.mde in a use case for microscopy metadata harmonization: Facilitating FAIR principles in practical application with metadata annotation tools. arXiv [q-bio.QM] (2021), (available at http://arxiv.org/abs/2103.02942).

50. J. Ryan, T. Pengo, A. Rigano, P. Montero Llopis, M. S. Itano, L. C. Cameron, G. Marqués, C. Strambio-De-Castillia, M. A. Sanders, C. M. Brown, MethodsJ2: A Tool to Help Improve Microscopy Methods Reporting. Nat. Methods(Manuscript under joint submission; personal communication).

51. M. Hammer, M. Huisman, A. Rigano, U. Boehm, J. J. Chambers, N. Gaudreault, J. A. Pimentel, D. Sudar, P. Bajcsy, C. M. Brown, A. D. Corbett, O. Faklaris, J. Lacoste, A. Laude, G. Nelson, R. Nitschke, A. J. North, R. Gopinathan, F. Farzam, C. Smith, W. Gipfel, J. Ritland, D. Grunwald, C. Strambio-De-Castillia, 4DN-BINA-OME (NBO)-Microscopy Metadata Specifications – Tiers System__v2.01 (GitHub – https://github.com/WU-BIMAC/NBOMicroscopyMetadataSpecs, 2021; https://github.com/WU-BIMAC/NBOMicroscopyMetadataSpecs/tree/master/Tier%20System/stable%20version/v02-01).

52. B. Alver, 4DN-DCIC, P. Park, 4DN Data Portal. https://data.4dnucleome.org/ (2018), (available at https://data.4dnucleome.org/).

53. A. Rigano, A. Balashov, S. Ehmsen, S. U. Ozturk, B. Alver, C. Strambio-De-Castillia, Micro-Meta App – Electron(Github – https://github.com/WU-BIMAC, 2021; https://doi.org/10.5281/zenodo.4750765).

54. A. Rigano, A. Balashov, S. Ehmsen, S. U. Ozturk, B. Alver, C. Strambio-De-Castillia, Micro-Meta App – React(Github – https://github.com/WU-BIMAC, 2021; https://doi.org/10.5281/zenodo.4751438).

55. C. Strambio-De-Castillia, A. Rigano, Micro-Meta App: Microscopy Metadata for the real world!*Micro-Meta App Website* (2020), (available at https://wu-bimac.github.io/MicroMetaApp.github.io/).

56. C. Strambio-De-Castillia, Micro-Meta App ReadTheDocs documentation. micrometaapp-docs.readthedocs.io(2020), (available at https://micrometaapp-docs.readthedocs.io/en/latest/index.html).

57. K. Bellve, A. Rigano, K. Fogarty, C. Strambio-De-Castillia, Example Microscopy Metadata JSON files produced using Micro-Meta App to document the acquisition of example images using the custom-built TIRF Epifluorescence Structured Illumination Microscope (2021), doi:10.5281/zenodo.4891883.

58. A. Rigano, S. U. Öztürk, S. Ehmsen, A. Cosolo, A. Balashov, B. Alver, C. Strambio-De-Castillia, Micro-Meta App – 4DN Data Portal (2021; https://data.4dnucleome.org/tools/micro-meta-app).

59. RRID Initiative, Research Resource Identifier (RRID). https://www.rrids.org/ (2015), (available at https://www.rrids.org/).

60. A. S. Abdelfattah, T. Kawashima, A. Singh, O. Novak, H. Liu, Y. Shuai, Y.-C. Huang, L. Campagnola, S. C. Seeman, J. Yu, J. Zheng, J. B. Grimm, R. Patel, J. Friedrich, B. D. Mensh, L. Paninski, J. J. Macklin, G. J. Murphy, K. Podgorski, B.-J. Lin, T.-W. Chen, G. C. Turner, Z. Liu, M. Koyama, K. Svoboda, M. B. Ahrens, L. D. Lavis, E. R. Schreiter, Bright and photostable chemigenetic indicators for extended in vivo voltage imaging. Science. 365, 699–704 (2019).

61. Y. Qian, K. D. Piatkevich, B. Mc Larney, A. S. Abdelfattah, S. Mehta, M. H. Murdock, S. Gottschalk, R. S. Molina, W. Zhang, Y. Chen, J. Wu, M. Drobizhev, T. E. Hughes, J. Zhang, E. R. Schreiter, S. Shoham, D. Razansky, E. S. Boyden, R. E. Campbell, A genetically encoded near-infrared fluorescent calcium ion indicator. Nat. Methods. 16, 171–174 (2019).

62. J. B. Grimm, A. N. Tkachuk, L. Xie, H. Choi, B. Mohar, N. Falco, K. Schaefer, R. Patel, Q. Zheng, Z. Liu, J. Lippincott-Schwartz, T. A. Brown, L. D. Lavis, A general method to optimize and functionalize red-shifted rhodamine dyes. Nat. Methods. 17, 815–821 (2020).

63. A. Kiepas, E. Voorand, F. Mubaid, P. M. Siegel, C. M. Brown, Optimizing live-cell fluorescence imaging conditions to minimize phototoxicity. J. Cell Sci. 133(2020), doi: 10.1242/jcs.242834.

64. A. Fernandez, C. A. Zentner, M. Shivrayan, E. Samson, S. Savagatrup, J. Zhuang, T. M. Swager, S. Thayumanavan, Programmable Emulsions via Nucleophile-Induced Covalent Surfactant Modifications. Chem. Mater. 32, 4663–4671 (2020).

65. W. C. Drosopoulos, D. A. Vierra, C. A. Kenworthy, R. A. Coleman, C. L. Schildkraut, Dynamic Assembly and Disassembly of the Human DNA Polymerase δ Holoenzyme on the Genome In Vivo. Cell Rep. 30, 1329–1341.e5 (2020).

66. N. V. Ayala-Nunez, G. Follain, F. Delalande, A. Hirschler, E. Partiot, G. L. Hale, B. C. Bollweg, J. Roels, M. Chazal, F. Bakoa, M. Carocci, S. Bourdoulous, O. Faklaris, S. R. Zaki, A. Eckly, B. Uring-Lambert, F. Doussau, S. Cianferani, C. Carapito, F. M. J. Jacobs, N. Jouvenet, J. G. Goetz, R. Gaudin, Zika virus enhances monocyte adhesion and transmigration favoring viral dissemination to neural cells. Nat. Commun. 10, 4430 (2019).

67. A. Dumas, G. Lê-Bury, F. Marie-Anaïs, F. Herit, J. Mazzolini, T. Guilbert, P. Bourdoncle, D. G. Russell, S. Benichou, A. Zahraoui, F. Niedergang, The HIV-1 protein Vpr impairs phagosome maturation by controlling microtubule-dependent trafficking. J. Cell Biol. 211, 359–372 (2015).

68. A. Aghajanian, H. Zhang, B. K. Buckley, E. S. Wittchen, W. Y. Ma, J. E. Faber, Decreased inspired oxygen stimulates de novo formation of coronary collaterals in adult heart. J. Mol. Cell. Cardiol. 150, 1–11 (2021).

69. N. A. Watson, T. N. Cartwright, C. Lawless, M. Cámara-Donoso, O. Sen, K. Sako, T. Hirota, H. Kimura, J. M. G. Higgins, Kinase inhibition profiles as a tool to identify kinases for specific phosphorylation sites. Nat. Commun. 11, 1684 (2020).

70. R. L. Upton, Z. Davies-Manifold, M. Marcello, K. Arnold, C. R. Crick, A general formulation approach for the fabrication of water repellent materials: how composition can impact resilience and functionality. Mol. Syst. Des. Eng. 5, 477–483 (2020).

71. H. C. Lim, T. G. Bernhardt, A PopZ-linked apical recruitment assay for studying protein-protein interactions in the bacterial cell envelope. Mol. Microbiol. 112, 1757–1768 (2019).

72. H. C. Lim, J. W. Sher, F. P. Rodriguez-Rivera, C. Fumeaux, C. R. Bertozzi, T. G. Bernhardt, Identification of new components of the RipC-FtsEX cell separation pathway of Corynebacterineae. PLoS Genet. 15, e1008284 (2019).

73. P. F. L. da Silva, M. Ogrodnik, O. Kucheryavenko, J. Glibert, S. Miwa, K. Cameron, A. Ishaq, G. Saretzki, S. Nagaraja-Grellscheid, G. Nelson, T. von Zglinicki, The bystander effect contributes to the accumulation of senescent cells in vivo. Aging Cell. 18, e12848 (2019).

74. P. Dalle Pezze, G. Nelson, E. G. Otten, V. I. Korolchuk, T. B. L. Kirkwood, T. von Zglinicki, D. P. Shanley, Dynamic Modelling of Pathways to Cellular Senescence Reveals Strategies for Targeted Interventions. PLoS Comput. Biol. 10, 1–20 (2014).

75. S. Proksch, G. Bittermann, K. Vach, R. Nitschke, P. Tomakidi, E. Hellwig, hMSC-Derived VEGF Release Triggers the Chemoattraction of Alveolar Osteoblasts. Stem Cells. 33, 3114–3124 (2015).

76. D. M. Navaroli, K. D. Bellve, C. Standley, L. M. Lifshitz, J. Cardia, D. Lambright, D. Leonard, K. E. Fogarty, S. Corvera, Rabenosyn-5 defines the fate of the transferrin receptor following clathrin-mediated endocytosis. Proceedings of the National Academy of Sciences. 109, E471–80 (2012).

77. L. Hannibal, J. Theimer, V. Wingert, K. Klotz, I. Bierschenk, R. Nitschke, U. Spiekerkoetter, S. C. Grünert, Metabolic Profiling in Human Fibroblasts Enables Subtype Clustering in Glycogen Storage Disease. Front. Endocrinol. 11, 579981 (2020).

78. J. A. Pimentel, J. Carneiro, A. Darszon, G. Corkidi, A segmentation algorithm for automated tracking of fast swimming unlabelled cells in three dimensions. J. Microsc. 245, 72–81 (2012).

79. F. Silva-Villalobos, J. A. Pimentel, A. Darszon, G. Corkidi, Imaging of the 3D dynamics of flagellar beating in human sperm. Conf. Proc. IEEE Eng. Med. Biol. Soc. 2014, 190–193 (2014).

80. A. Rigano, U. Boehm, J. J. Chambers, N. Gaudreault, J. A. Pimentel, D. Sudar, P. Bajcsy, C. M. Brown, A. D. Corbett, O. Faklaris, J. Lacoste, A. Laude, G. Nelson, R. Nitschke, A. J. North, D. Grunwald, C. Strambio-De-Castillia, 4DN-BINA-OME (NBO) Tiered Microscopy Metadata Specifications – v2.00 (GitHub – https://github.com/WU-BIMAC/NBOMicroscopyMetadataSpecs, 2021; https://zenodo.org/record/4682257).

81. GlobalBioimaging, Global Bioimaging website. https://globalbioimaging.org/ (2018), (available at https://globalbioimaging.org/).

82. M. Huisman, thesis, University of Massachusetts Medical School (2019).

83. A. Rigano, W. Moore, S. Ehmsen, B. Alver, C. Strambio-De-Castillia, Micro-Meta App – OMERO plug-in (Github – https://github.com/WU-BIMAC, 2021; https://doi.org/10.5281/zenodo.4750762).

84. T. J. Lambert, FPbase: a community-editable fluorescent protein database. Nat. Methods. 16, 277–278 (2019).

85. T. Lambert, FPbase (Github-https://github.com/tlambert03/FPbase, 2019; https://github.com/tlambert03/FPbase).

86. C. Allan, J.-M. Burel, J. Moore, C. Blackburn, M. Linkert, S. Loynton, D. MacDonald, W. J. Moore, C. Neves, A. Patterson, M. Porter, A. Tarkowska, B. Loranger, J. Avondo, I. Lagerstedt, L. Lianas, S. Leo, K. Hands, R. T. Hay, A. Patwardhan, C. Best, G. J. Kleywegt, G. Zanetti, J. R. Swedlow, OMERO: flexible, model-driven data management for experimental biology. Nat. Methods. 9, 245–253 (2012).

87. L. Allen, A. O’Connell, V. Kiermer, How can we ensure visibility and diversity in research contributions? How the Contributor Role Taxonomy (CRediT) is helping the shift from authorship to contributorship. Learn. Publ. 32, 71–74 (2019).

88. Elsevier editors, CRediT author statement – editorial. Elsevier (2020) (available at https://www.elsevier.com/authors/policies-and-guidelines/credit-author-statement).

