## Supplemental Material for "Micro-Meta App: an interactive software tool to facilitate the collection of microscopy metadata based on community-driven specifications"

#### SUPPLEMENTAL ONLINE RESOURCES

##### Example JSON Files - Figure 5 Use Case

Example **Microscopy Metadata JSON files** produced using Micro-Meta App to document the acquisition of the **FSWT-6hVirus-10minFIX-stk_4-EPI.tif.ome.tif** example image-file using the custom-built TIRF Epifluorescence Structured Illumination Microscope (TESM) custom-built by the Biomedical Imaging Group at the University of Massachusetts Medical School (Navaroli et al., 2010) as depicted in Figure 5 are available on Zenodo at: https://doi.org/10.5281/zenodo.4891883

Also available at the same location is the example **FSWT-6hVirus-10minFIX-stk_4-EPI.tif.ome.tif** image utilized in this case.

#### SUPPLEMENTAL VIDEOS

##### Supplemental Video 1: Micro-Meta App-an introduction

A video that was used as an introduction to Micro-Meta App for use by Graduate Students at the University of Massachusetts Medical School is available at: https://vimeo.com/557097919

#### SUPPLEMENTAL METHODS AND SOFTWARE IMPLEMENTATION

##### Software implementation

Micro-Meta App is available in two JavaScript (JS) implementations. The first implementation was designed to facilitate the incorporation of the software in existing third-party web servers (i.e., the 4DN data portal) (Rigano et al., 2021d; a) and was developed using the JavaScript React library, which is widely used to build web-based user interfaces. Starting from this version and to lower the barrier of adoption of the tool by microscope-users, laboratories and core-facilities that do not have access to imaging databases, a stand-alone version of the App was developed by wrapping the React implementation using the JavaScript Electron library.

**Dependencies**

For the Micro-Meta App to work, the following elements are generated in advance as described in the following sections and made available via GitHub:

1. JSON schema: a JSON file or a series of files that define the underlying schema, which is used to construct the GUI of the application on the basis of the 4DN-BINA-OME Microscopy Metadata specifications (Hammer et al., 2021).

2. Dimensions and coordinates: a JSON file that codifies the dimensions of the canvases utilized by the Manage Instrument Graphical User Interface (GUI) of the Micro-Meta App as well as the position on the canvas occupied by icons representing each microscope hardware component.
3. Icons: A series of SVG files, each containing an icon representing the individual hardware components.

In addition, the stand-alone version of Micro-Meta App depends on the 4DN Microscopy Metadata Reader (Rigano and Strambio-De-Castillia, 2021a), which implements the BioFormats library (Linkert et al., 2010) to allow the user to import known OME-metadata directly from the file-header of an image data file of interest. To maximize flexibility, these elements can be customized by individual users.

**XSD to JSON Schema converter**

The main function of this Java-encoded component (Rigano and Strambio-De-Castillia, 2021b) is to transform the XML Schema Definition (XSD) implementation of the 4DN-BINA-OME Data Model (Hammer et al., 2021; Huisman et al., 2021; Rigano et al., 2021c) into a JSON-based schema, which is subsequently ingested by Micro-Meta App to automatically generate the software GUI and the associated data insertion forms. The XSD to JSON Schema converter middleware utilizes the Xerces2 Java XML Parser (The Apache Software Foundation, 2018) and the W3C Java XML bindings libraries (W3C, 2003) to navigate the XSD schema and produces two kinds of version-aware JSON files:

1. A comprehensive JSON file containing an array of the schemas for all necessary individual components that constitutes the 4DN-BINA-OME Data Model (e.g., Objective, Filter, or Detector). This comprehensive JSON file is made available on GitHub and is specifically designed to facilitate the remote loading of the schema by web-portal embedded React implementations of the Micro-Meta App. This comprehensive JSON schema is available as an individual file on GitHub [1].
2. A series of JSON files, each containing the schema of individual components, which were designed to be employed by the Electron implementation of the App. These individual schema files are available within a subdirectory of the main repository on GitHub[2].

The middleware was specifically designed to maximize flexibility and extensibility. As such, the software allows the introduction of implementation-specific modifications of the resulting JSON schema so that it can be adapted for special purposes. For example, the introduction of a "Version" field allows the validation of whether the data being saved is compatible with the specific version of the schema being employed. As a further example, the introduction of the "Category" field allows the organization of different components in specific sub-menus across the sidebar. To facilitate the evolution of the model while ensuring back-compatibility, the GitHub repository supports versioning by storing all revisions of the output JSON schema.

**TXT to JSON Dimensions converter**

This Java-encoded component (Rigano and Strambio-De-Castillia, 2021b) is used to process an input text file containing the dimensions of the Manage Instrument canvas of Micro-Meta App alongside the desired XY positions where each icon has to be placed. As a result, the software produces a Dimensions and Coordinate JSON file that is made available for remote loading from GitHub [3] and is ready to be used to implement the "snap-in-place" functionality of the software.

---

[1] https://github.com/WU-BIMAC/4DNMetadataSchemaXSD2JSONConverter/blob/master/latest/fullSchema.json
[2] https://github.com/WU-BIMAC/4DNMetadataSchemaXSD2JSONConverter/tree/master/latest/schemas
[3] https://github.com/WU-BIMAC/4DNMetadataSchemaXSD2JSONConverter/tree/master/latest/dimensions

**ICON**

SVG files containing icons representing the different microscope hardware components were generated specifically for this project and are made available in a version-aware manner for remote loading from GitHub (Rigano and Strambio-De-Castillia, 2021b)[4]. Custom-made icons can be similarly generated by developer users and manufacturers for representation of their hardware components within the application.

**Microscopy Metadata Reader**

This software is written in JAVA to fully take advantage of the Bio-Formats library and the OME-XML metadata structure (Rigano and Strambio-De-Castillia, 2021a). Using these two dependencies, this software accesses all the OME-compatible metadata present in user-selected images and maps it to the 4DN-BINA-OME Microscopy Metadata specifications (Hammer et al., 2021; Rigano et al., 2021c) that extend the OME Data Model (Goldberg et al., 2005; Linkert et al., 2010) to produce a temporary, Micro-Meta App-compatible JSON object that can be read by Micro-Meta App. The object is then passed on to the Micro-Meta App and read by the Manage Settings section of the App to pre-populate the corresponding metadata fields.

#### JavaScript React implementation of Micro-Meta App

This is the core implementation of the Micro-Meta App, and it is the starting point for embedding it into a third-party web-portal, and for wrapping it into Electron for local execution (Rigano et al., 2021a). This component ingests JSON schema files, JSON Dimension and Coordinate files and Icon images produced as described above and uses a series of custom React classes to produce a set of individual windows composing the GUI. Specifically, these windows can be categorized into two main sections: the Instrument Manager and the Settings Manager (Figure 2). In the Instrument Manager section, the application employs a Canvas setup that provides the flexibility to incorporate as many icon elements as necessary to describe the hardware components of any given microscope. For this purpose, the toolbar is generated dynamically as dictated by the underlying JSON schema produced as described above and by the selected tier level. On this basis, elements present in each of the graphical menus present on the sidebar can be dragged to the canvas and dropped either to a custom position specified by the user, or they can be snapped-in-place as defined by the position and dimension file produced by the Dimension Converter component.

In the Settings Manager, the application builds a series of nested windows that are launched by individual buttons allowing the user to select individual hardware components and enter specific settings for each of them. Of particular interest is the Channel interface in which the user can define the LightPath configuration associated with each channel of a given image by following a predefined graphical flowchart (Figure 5D).

#### JavaScript Electron implementation of Micro-Meta App

This implementation of the Micro-Meta App is encoded in JavaScript (Rigano et al., 2021b). It utilizes the Electron library to wrap the React implementation of Micro-Meta App with all necessary schemas, icon images, and position/dimension to ensure its correct functionality. This discrete executable file can interact directly with the underlying operating system and launch the software from the local file system.

[4] https://github.com/WU-BIMAC/4DNMetadataSchemaXSD2JSONConverter/tree/master/latest/images

#### SUPPLEMENTAL FIGURES

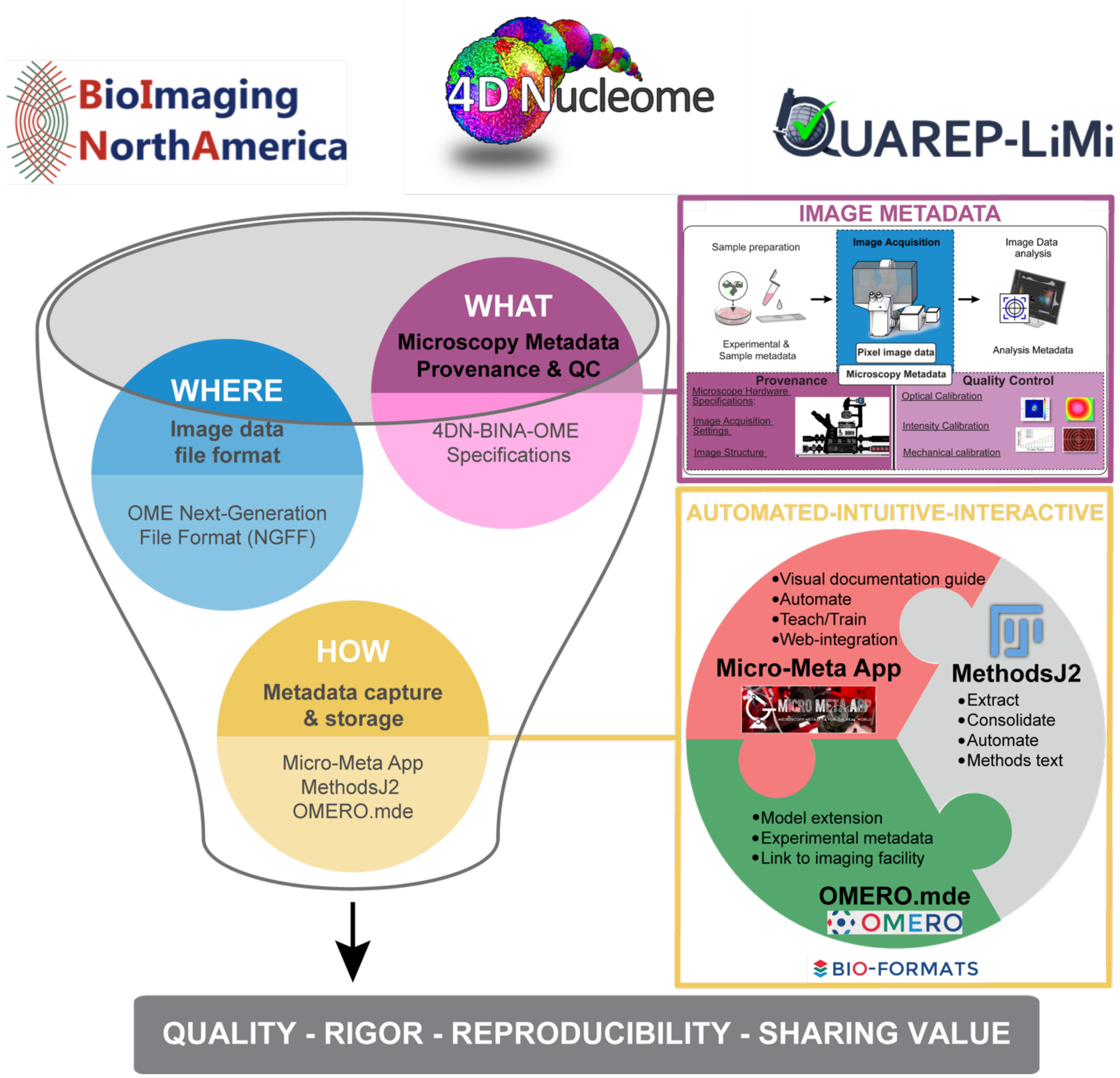

**Supplemental Figure 1: Quality, rigor, reproducibility and sharing value for imaging experiments require the definition of community-driven Microscopy Metadata specifications and the adoption of easy-to-use metadata collection tools to facilitate the documentation and quality control tasks for experimental scientists.** The establishment of FAIR (Wilkinson et al., 2016), community-driven Microscopy Image Data Standards implies parallel development on three interrelated fronts: (*WHERE*) Next-Generation File Formats (NGFF) where the ever-increasing scale and complexity of image data and metadata would be contained for exchange (Moore et al., 2021); *blue bubble*). (*WHAT*) Community-driven specifications for what 'data provenance' information (microscope hardware specifications, image acquisition settings and image structure metadata) and quality control metrics are essential for rigor, reproducibility, and reuse and should therefore be captured in Microscopy Metadata (*magenta bubble*). (*HOW*) Shared rules for how the (ideally) automated capture, representation and storage of Microscopy Metadata should be implemented in practice (*yellow bubble*). Micro-Meta App, MethodsJ2 and OMERO.mde are three highly interoperable tools and complementary that function to: 1) train users on the importance of documentation and quality control; 2) facilitate metadata extraction, collection, and storage; 3) automatically write Methods sections; and 4) facilitate the development of experimental metadata specifications in connection with local core facilities. To facilitate adoption by users with different use-styles and preferences, the three tools each work in a specialized environment: Micro-Meta App is used as stand-alone app or can be integrated in third-party image data portals. MethodsJ2 works as an ImageJ plugin. OMERO.mde works in the context of the OMERO image data repository. The different tools are based on different software platforms in order to appeal to the broadest community including microscope builders, imaging scientists working in core facilities and experimental scientists. The concept is to bring the tools to software platforms people are already using and lower the bar to enable broad uptake.

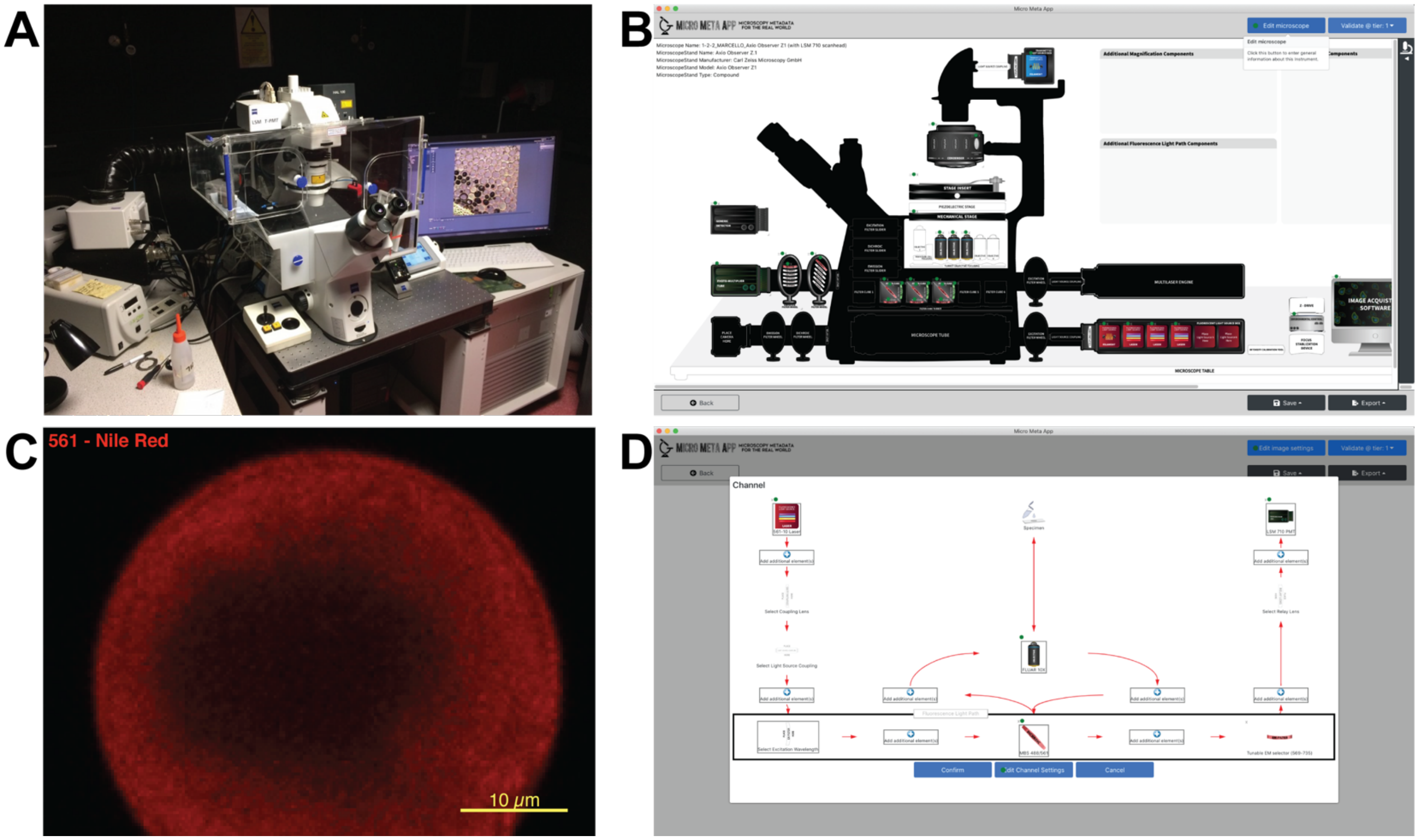

**Supplemental Figure 2 | Micro-Meta App was utilized by several light microscopy core facilities to document individual microscope instrumentation and the settings that were applied to the microscope to acquire specific example datasets**. Illustrated here is the use of Micro-Meta App for Tier 1 *(Minimum Information/Materials & Methods)*, 4DN-BINA-OME-specified (Hammer et al., 2021) documentation of: **(A-B)** the ***Zeiss*** Axio Observer Z1 inverted microscope equipped with a LSM 710 confocal scan head, owned by the Centre for Cell Imaging (CCI) of the University of Liverpool (Supplemental Table III); and **(C-D)** the settings that were applied to the microscope for the acquisition of example published image data sets (Upton et al., 2020). **A)** Picture of the indicated microscope. **B)** Micro-Meta App generated schematic representation of the indicated microscope. **C)** Particles composed of superhydrophobic polymer–nanoparticle composite (SPNC) material were analyzed by confocal microscopy to shed light on the differences in composite architecture between different coating polymers. Preparations of SiO2-polydimethylsiloxane (PDMS) coated SPNC (40–75 μm) particles were labelled by staining PDMS with Nile Red. Confocal 3D Z-stacks were acquired using the microscope indicated in Panels A and B. Depicted here is a slice from the middle (24 μm) of a Z-stack depicting a representative particle, which is presented with permission from Upton et al., 2020. **D)** Micro-Meta App generated schematic representation of the light path associated with the ***561 Channel*** utilized for the acquisition of the image in Panel C.

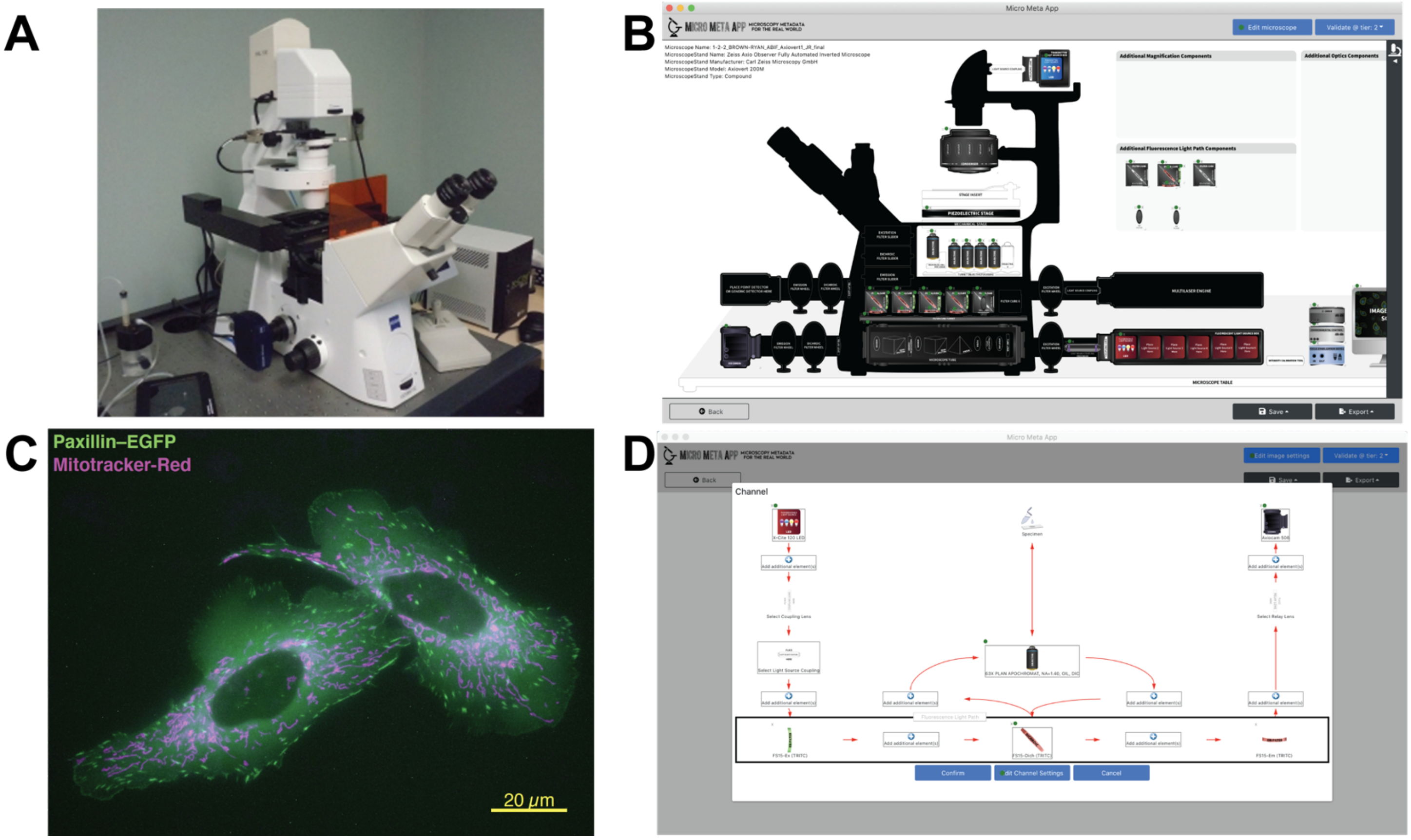

**Supplemental Figure 3 | Micro-Meta App was utilized by several light microscopy core facilities to document individual microscope instrumentation and the settings that were applied to the microscope to acquire specific example datasets**. Illustrated here is the use of Micro-Meta App for Tier 2 *(Advanced Quantification and/or Live cell Imaging)*, 4DN-BINA-OME-specified (Hammer et al., 2021) documentation of: **(A-B)** the ***Zeiss Axio Observer (Axiovert 200M)*** inverted, epifluorescence microscope owned by the Advanced BioImaging Facility (ABIF) of McGill University (Supplemental Table III); and **(C-D)** the settings that were applied to the microscope for the acquisition of example published image data sets (Kiepas et al., 2020). **A)** Picture of the indicated microscope. **B)** Micro-Meta App generated schematic representation of the indicated microscope. **C)** Chinese Hamster Ovary K1 (CHO-K1) cells stably expressing paxillin–EGFP to visualize the cytoplasm and adhesions were seeded onto fibronectin-coated glass coverslips, allowed to adhere, and grow overnight, and stained with MitoTracker Red to visualize mitochondrial morphology. Images for Paxillin–EGFP (GFP channel, visualized in green) and MitoTracker™ Red (TRITC channel, visualized in magenta) were acquired every minute and 6 min, respectively, using the indicated microscope (Panels A and B) with Diffuse Light Delivery (DLD; 0.0093 mW×60,000 ms) illumination. Displayed is a representative image from time zero adapted with permission from Kiepas et al., 2020. **D)** Micro-Meta App generated schematic representation of the light path associated with the ***TRITC Channel*** utilized for the acquisition of the image in Panel C.

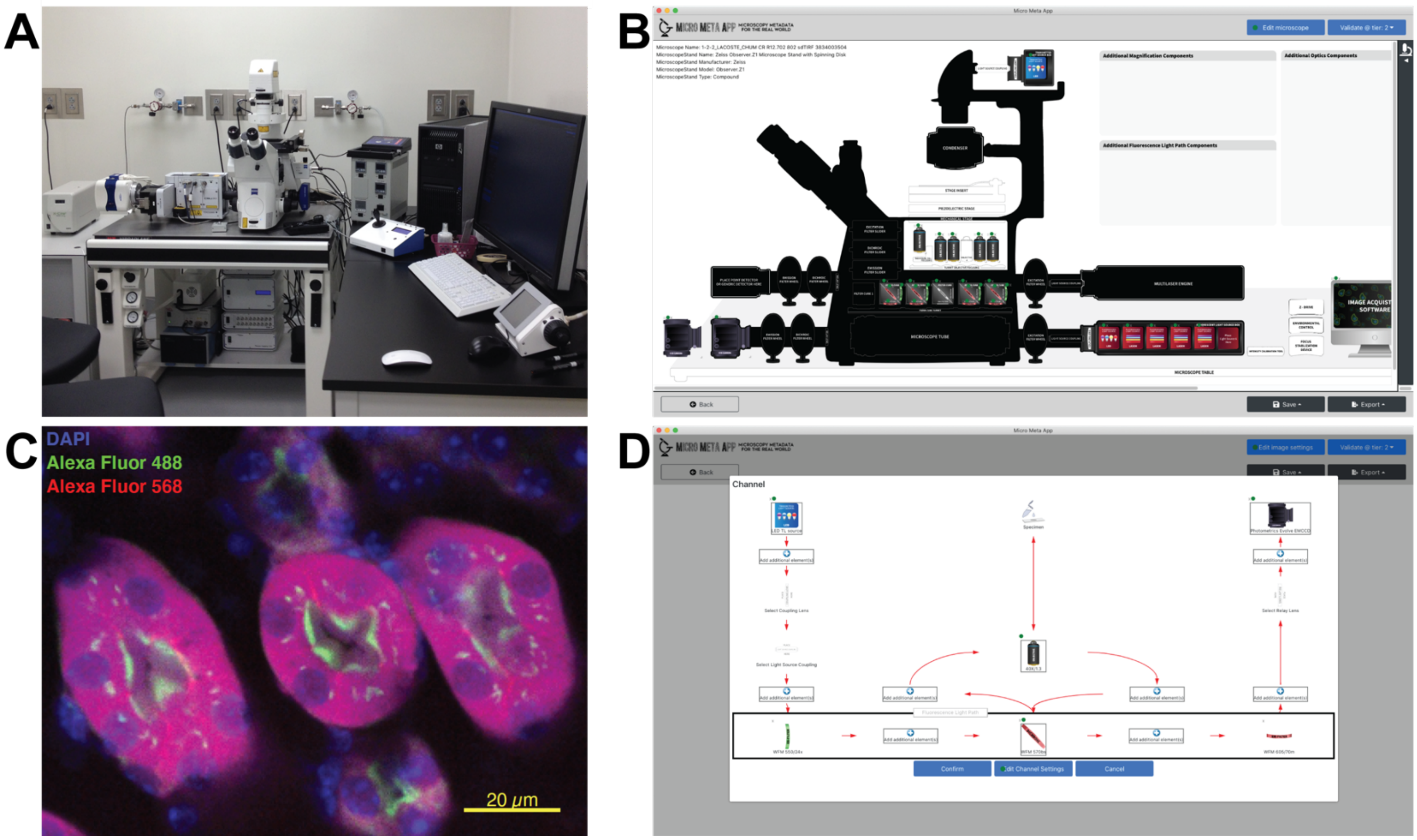

**Supplemental Figure 4 | Micro-Meta App was utilized by several light microscopy core facilities to document individual microscope instrumentation and the settings that were applied to the microscope to acquire specific example datasets**. Illustrated here is the use of Micro-Meta App for Tier 2 *(Advanced Quantification and/or Live cell Imaging)*, 4DN-BINA-OME-specified (Hammer et al., 2021) documentation of: **(A-B)** the ***Zeiss Observer.Z1*** inverted, epifluorescence microscope with Spinning Disk owned by the "Imagerie Cellulaire" core facility of the CR CHUM (Centre de recherche du Centre Hospitalier de l'Université de Montréal), and whose calibration and performance quality control is managed by MIA Cellavie Inc. (Supplemental Table III); and **(C-D)** the settings that were applied to the microscope for the acquisition of the indicated example image data sets**. A)** Picture of the indicated microscope. **B)** Micro-Meta App generated schematic representation of the indicated microscope. **C)** Displayed is a representative single-plane image of a 16 µm cryosection of Mouse kidney stained with DAPI (Blue channel, displayed in blue), WGA-AlexaFluor488 (Green channel, displayed in green), and Phalloidin-Alexa Fluor 568 (Red channel, displayed in red), and mounted in Gelvatol mounting medium. The displayed image was obtained using the indicated microscope in Panels A and B. **D)** Micro-Meta App generated schematic representation of the light path associated with the ***Red Channel*** utilized for the acquisition of the image in Panel C.

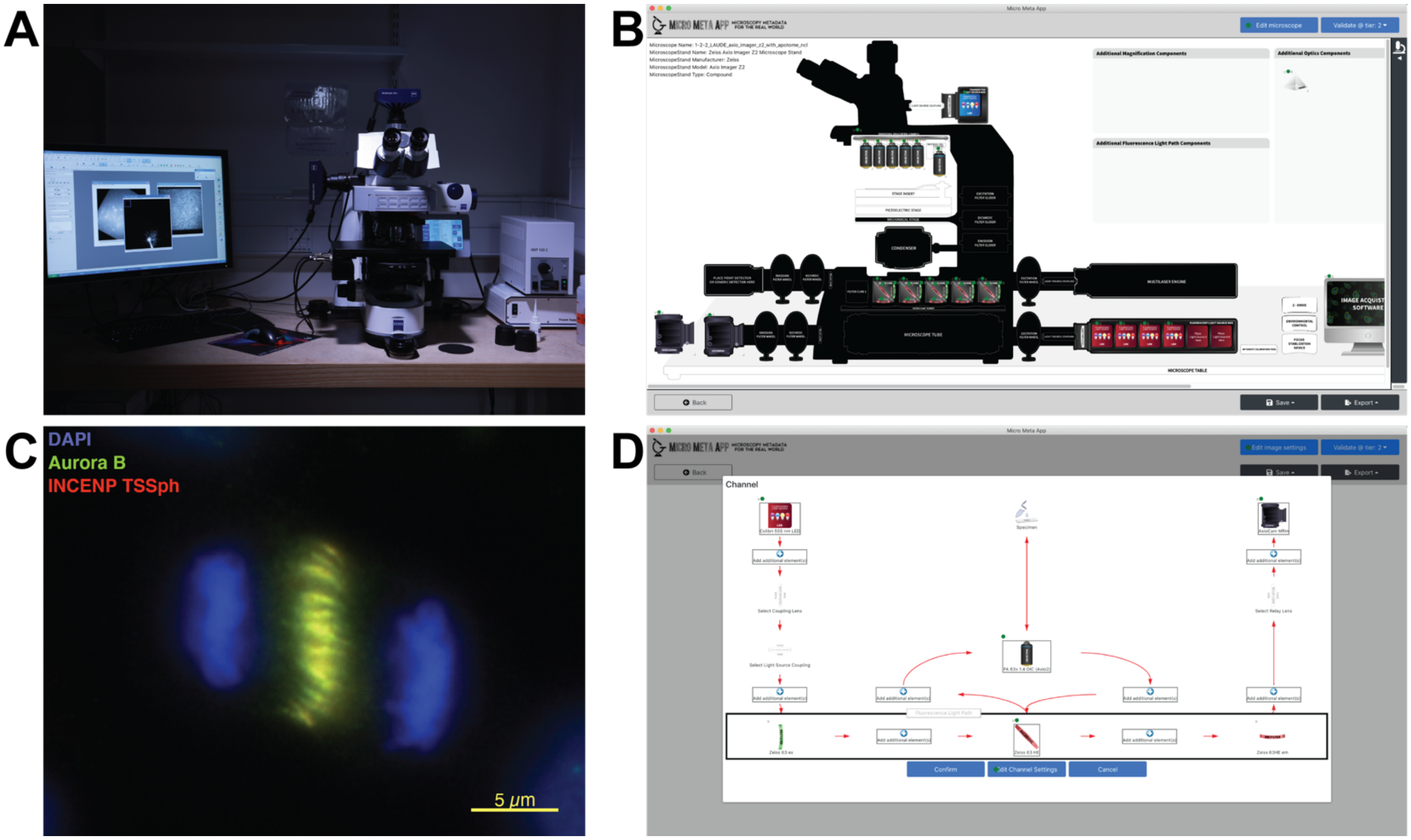

**Supplemental Figure 5: Micro-Meta App was utilized by several light microscopy core facilities to document individual microscope instrumentation and the settings that were applied to the microscope to acquire specific example datasets.** Illustrated here is the use of Micro-Meta App for Tier 2 *(Advanced Quantification and/or Live cell Imaging)*, 4DN-BINA-OME-specified (Hammer et al., 2021) documentation of: **(A-B)** the ***Zeiss Axio Imager Z2*** upright epifluorescence microscope with Apotome owned by the Bioimaging Unit of the University of Newcastle (Supplemental Table III); and **(C-D)** the settings that were applied to the microscope for the acquisition of example published image data sets (Watson et al., 2020). **A)** Picture of the indicated microscope. **B)** Micro-Meta App generated schematic representation of the indicated microscope. **C)** Mitotic HeLa cells were labeled for: DNA (DAPI DNA stain, DAPI channel, Zeiss 49 Filter set, visualized in blue), Aurora B (primary sheep anti-Aurora B polyclonal antibody, followed by donkey anti-sheep Alexa Fluor 488 secondary; GFP channel, Zeiss 38HE Filter set, displayed in green), and INCENP TSSph (primary rabbit anti-INCENP-TSSph polyclonal antibody, followed by donkey anti-rabbit Alexa Fluor 488 secondary; Red channel, Zeiss 63HE Filter set, displayed in red). Images were captured by widefield fluorescence microscopy using the microscope displayed in A & B equipped with a Colibri2 LED excitation light source, a Zeiss Plan Apo 63x/ 1.4NA objective, and a AxioCam MRmV3 camera. The displayed image was obtained using the indicated microscope (Panels A and B) and was adapted with permission from Watson et al., 2020. **D)** Micro-Meta App generated schematic representation of the light path associated with the ***INCENP-TSSph (Alexa Fluor 594) Channel*** utilized for the acquisition of the image in Panel C.

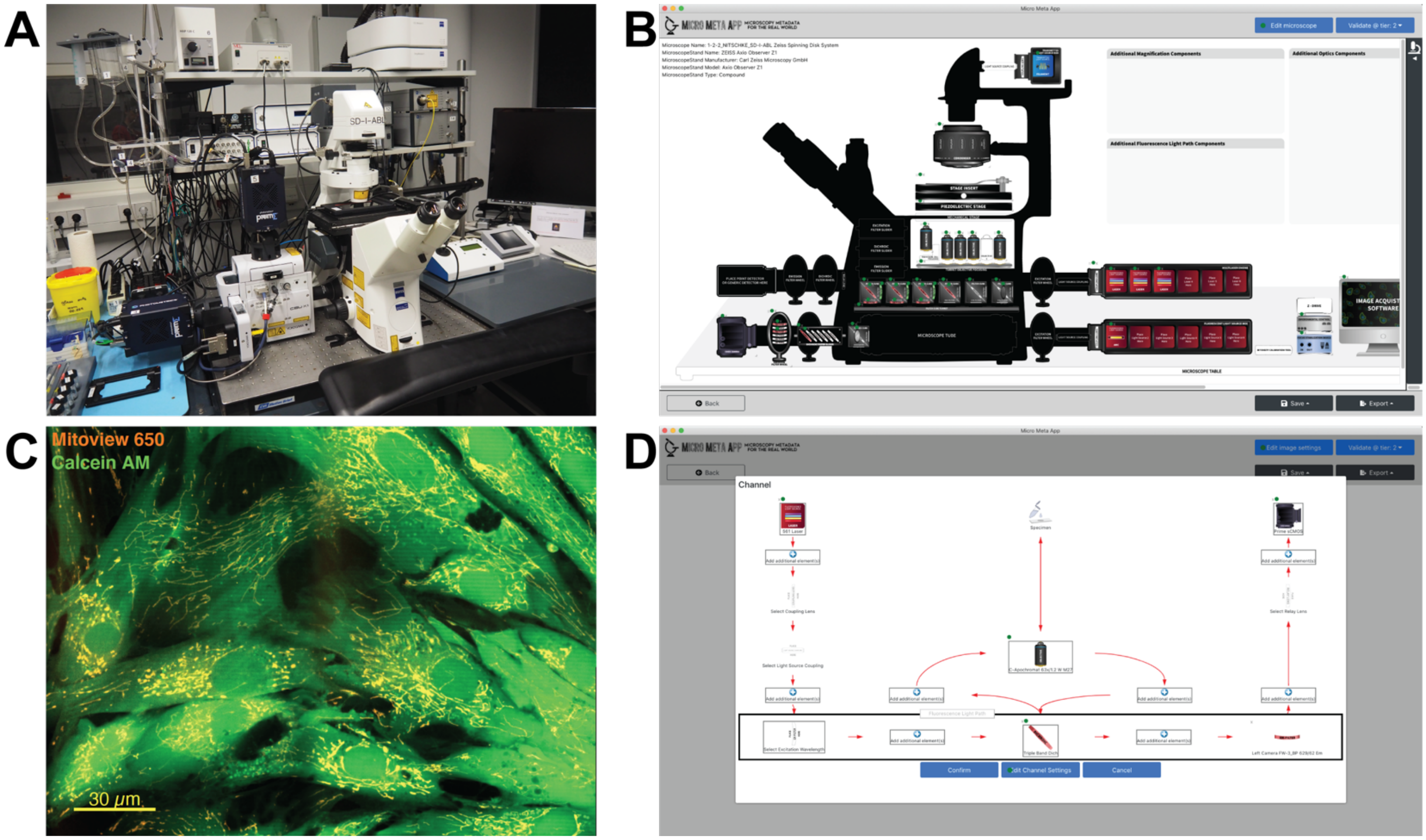

**Supplemental Figure 6 | Micro-Meta App was utilized by several light microscopy core facilities to document individual microscope instrumentation and the settings that were applied to the microscope to acquire specific example datasets**. Illustrated here is the use of Micro-Meta App for Tier 2 *(Advanced Quantification and/or Live cell Imaging)*, 4DN-BINA-OME-specified (Hammer et al., 2021) documentation of: **(A-B)** the ***Zeiss Axio Observer Z1*** inverted, epifluorescence microscope owned by the Life Imaging Center (LIC) Centre for Integrative Signaling Analysis (CISA) of the University of Freiburg (Supplemental Table III); and **(C-D)** the settings that were applied to the microscope for the acquisition of example published image data sets (Hannibal et al., 2020). **A)** Picture of the indicated microscope. **B)** Micro-Meta App generated schematic representation of the indicated microscope. **C)** Microscopy of live human skin fibroblasts from Glycogen Storage Disease (GSD) patients. After growth under standard conditions (humidified 5% CO2 incubator at 37°C) for 3–5 days, cells were trypsinized and 1000 cells were seeded in glass-bottom cell culture dishes. Cells were grown overnight under standard conditions and live cell staining was applied using Calcein™ AM for cell body (Ch 1 - GFP channel, displayed in green), and MitoView™ 650 for mitochondria (Ch 2 - AF555 channel, displayed in red). A representative overlay image is presented with permission from Hannibal et al., 2020. **D)** Micro-Meta App generated schematic representation of the light path associated with the ***Ch2 – AF555 Channel*** utilized for the acquisition of the image in Panel C.

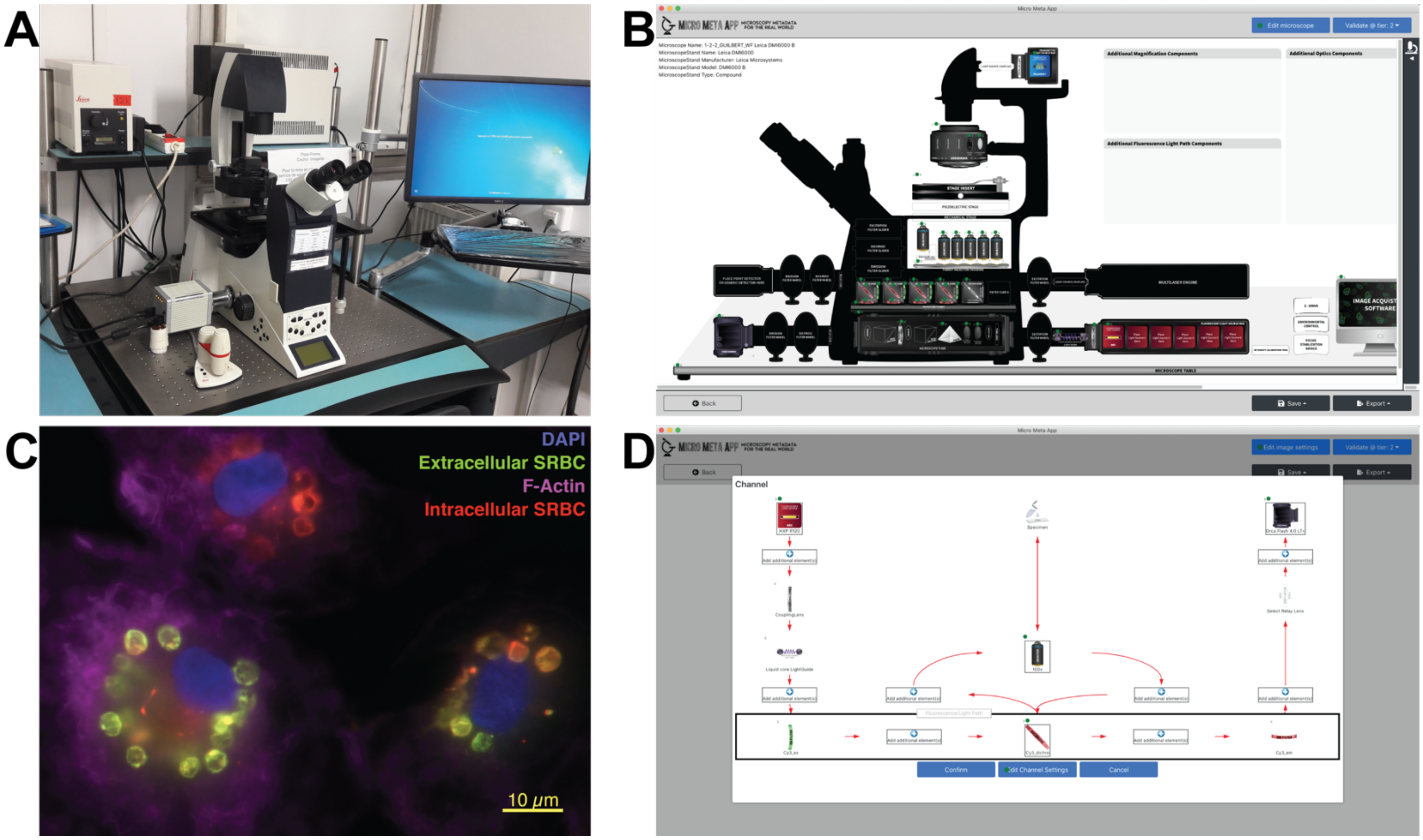

**Supplemental Figure 7 | Micro-Meta App was utilized by several light microscopy core facilities to document individual microscope instrumentation and the settings that were applied to the microscope to acquire specific example datasets**. Illustrated here is the use of Micro-Meta App for Tier 2 *(Advanced Quantification and/or Live cell Imaging)*, 4DN-BINA-OME-specified (Hammer et al., 2021) documentation of: **(A-B)** the ***Leica Microsystems DMI6000*** inverted epifluorescence microscope owned by the IMAG'IC Confocal Microscopy Facility of the Institut Cochin, at the Université de Paris (Supplemental Table III); and **(C-D)** the settings that were applied to the microscope for the acquisition of example published image data sets (Jubrail et al., 2020). **A)** Picture of the indicated microscope. **B)** Micro-Meta App generated schematic representation of the indicated microscope. **C)** Primary human macrophages were treated with human rhinovirus 16 (HRV16) for one hour, washed, and allowed to rest overnight at 37°C. The cells were then incubated at 37°C for 60 min with rabbit IgG-opsonized Sheep Red Blood Cells (SRBCs) to induce phagocytosis. At each time point, macrophages were fixed and labeled with Alexa Fluor 488-labeled F (ab') anti-rabbit IgG to detect the external SRBCs (GFP channel, visualized in green), then permeabilized and labeled with Cy5-labelled F (ab') anti-rabbit IgG to detect the internal SRBCs (CY5 channel, visualized in red), with Alexa Fluor 546-conjugated phalloidin to detect F-actin (CY3 Channel, visualized in magenta), and DAPI to stain the nuclei (DAPI channel, visualized blue). The displayed image was obtained using the indicated microscope (Panels A and B) and was adapted with permission from Jubrail et al., 2020. **D)** Micro-Meta App generated schematic representation of the light path associated with the ***F-Actin (Phalloidin-AF546) Channel*** utilized for the acquisition of the image in Panel C.

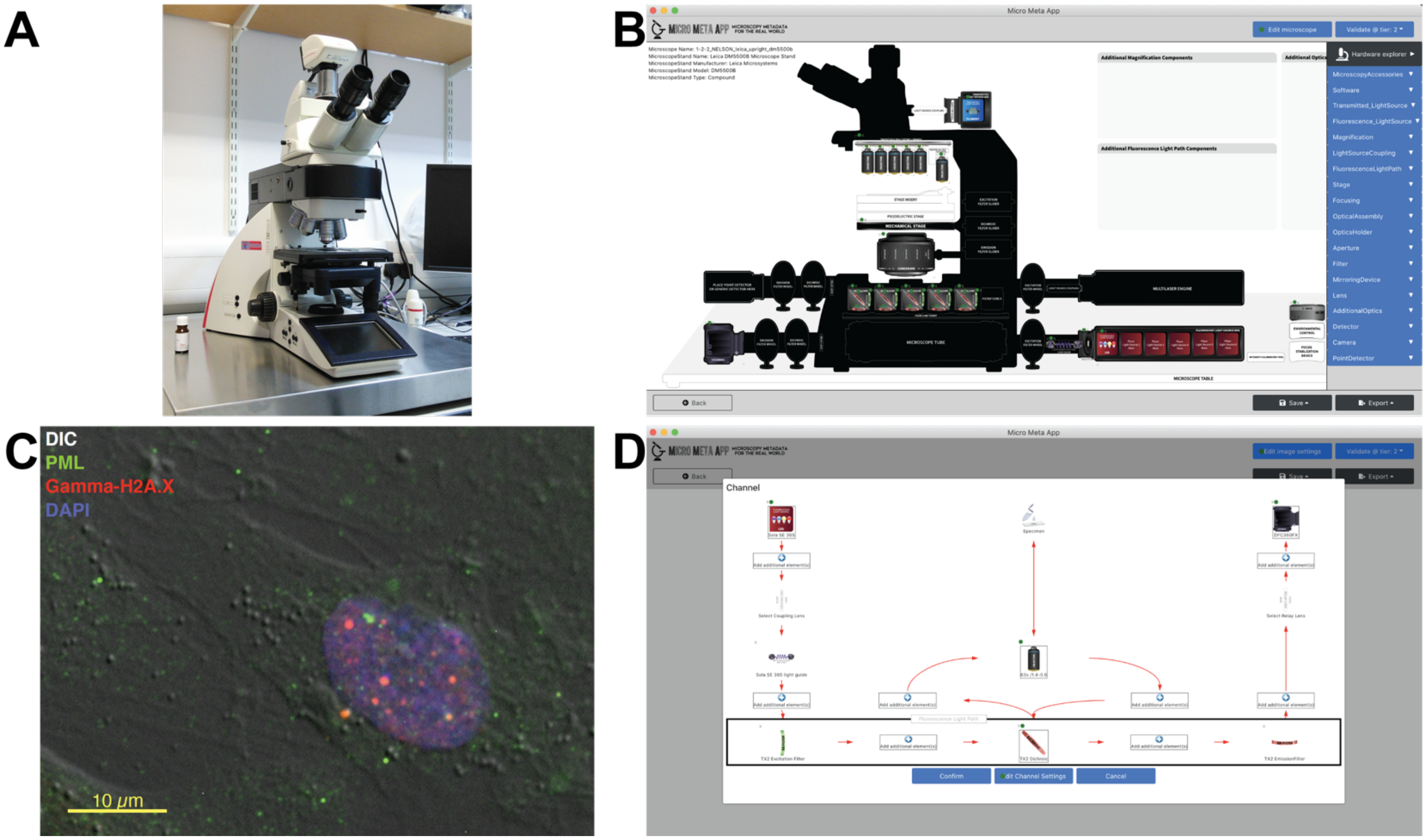

**Supplemental Figure 8 | Micro-Meta App was utilized by several light microscopy core facilities to document individual microscope instrumentation and the settings that were applied to the microscope to acquire specific example datasets**. Illustrated here is the use of Micro-Meta App for <u>Tier 2</u> *(Advanced Quantification and/or Live cell Imaging)*, 4DN-BINA-OME-specified (Hammer et al., 2021) documentation of: **(A-B)** the ***Leica Microsystems DM5500B*** upright, epifluorescence microscope owned by the Bioimaging Unit and located in the Edwardson Building on the Campus for Ageing and Vitality of the University of Newcastle (Supplemental Table III); and **(C-D)** the settings that were applied to the microscope for the acquisition of example published image data sets (Nelson et al., 2012). **A)** Picture of the indicated microscope. **B)** Micro-Meta App generated schematic representation of the indicated microscope. **C)** Immunofluorescence of young human diploid fibroblasts measuring the frequency of colocalization of PML bodies (as measured by staining cells with anti-PML primary antibody, followed by Alexa Fluor 488-conjugated secondary antibody; green) with DNA double-strand breaks (as measured by staining cells with anti-gamma-H2A.X primary antibody, followed by Alexa Fluor 555-conjugated secondary antibody; red), overlaid with a DIC transmission image (grey) and DAPI nuclear counterstain (blue) when grown in the presence or absence of senescent cells for 10 or 20 days. The displayed image was obtained using the indicated microscope (Panels A and B) and was adapted with permission from Nelson et al., 2012. **D)** Micro-Meta App generated schematic representation of the light path associated with the ***CY3 (gamma-H2A.X; red) Channel*** utilized for the acquisition of the image in Panel C.

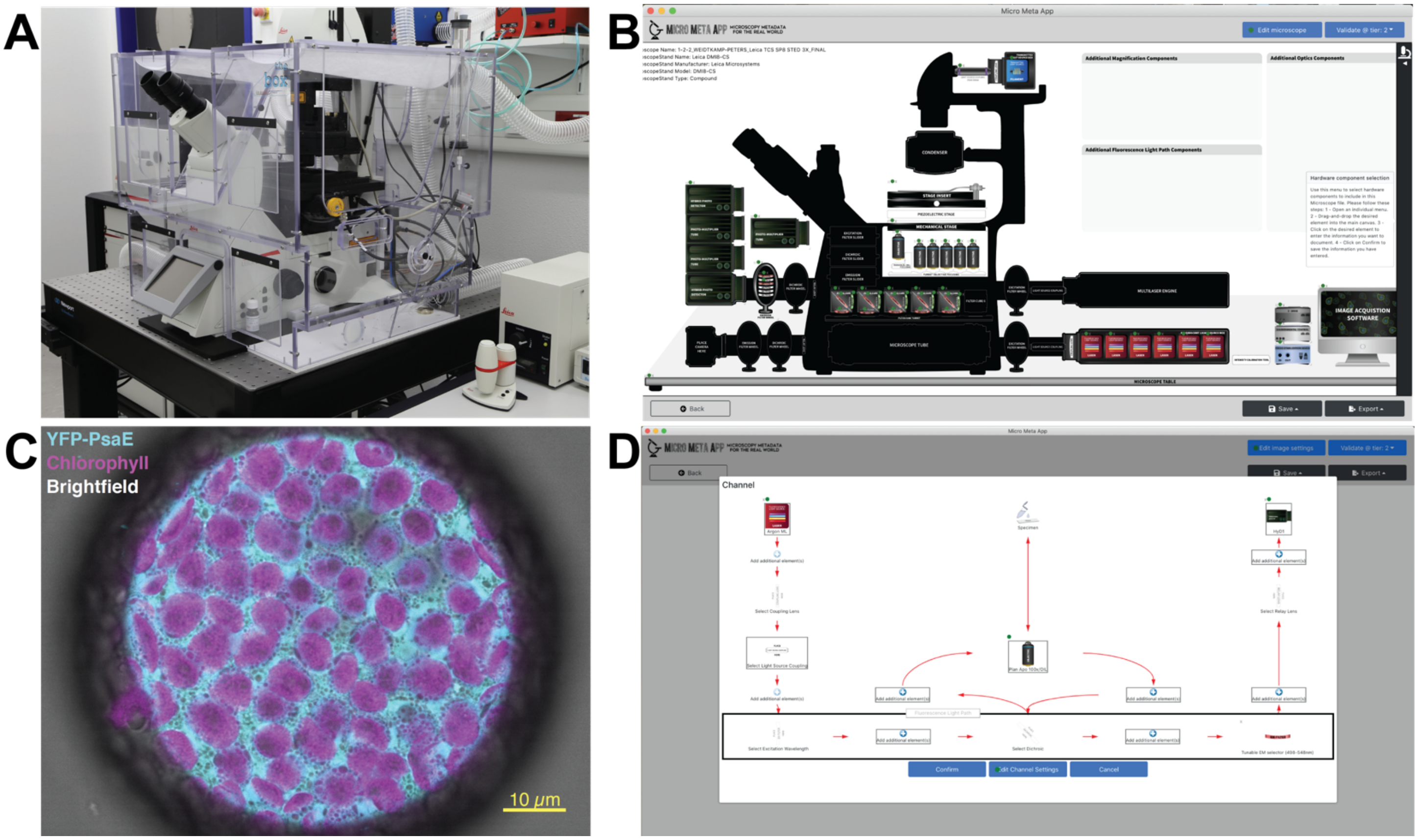

**Supplemental Figure 9: Micro-Meta App was utilized by several light microscopy core facilities to document individual microscope instrumentation and the settings that were applied to the microscope to acquire specific example datasets**. Illustrated here is the use of Micro-Meta App for Tier 2 *(Advanced Quantification and/or Live cell Imaging)*, 4DN-BINA-OME-specified (Hammer et al., 2021) documentation of: **(A-B)** the ***Leica Microsystems TCS SP8 STED 3X laser scanning confocal microscope system (with DMI8-CS inverted compound microscope stand)***, owned by the Center for Advanced imaging (CAi) at the School of Mathematics and Natural Sciences of the Heinrich-Heine-Universität Düsseldorf (Supplemental Table III); and **(C-D)** the settings that were applied to the microscope for the acquisition of example published image data sets (Singer et al., 2017). **A)** Picture of the indicated microscope. **B)** Micro-Meta App generated schematic representation of the indicated microscope. **C)** *N. benthamiana* cells were transformed to transiently express an N-Terminally YPF-tagged variant of the *P. chromatophora* PsaE protein, and prepared to produce protoplasts. Washed protoplasts were subsequently mounted on glass slides and prepared for multi-color fluorescence and transmitted light microscopy observation using the microscope illustrated in Panels A and B, equipped with a Leica HC PL APO 100x/ 1.40 OIL CS2 objective. Shown with permission (Singer et al., 2017) is a representative image of a protoplast in which the YFP fluorescence channel (*YFP-PsaE*, 488 laser, 498-548nm emission selection; displayed in Cyan) is overlaid with the Chlorophyll autofluorescence channel (*Chlorophyll*, 488 laser, 649-687nm emission selection; displayed in Magenta) and with the transmitted light channel (*Brightfield*; displayed in Grey). **D)** Micro-Meta App generated schematic representation of the light path associated with ***Channel 0 (YFP-PsaE)*** utilized for the acquisition of the image in Panel C.

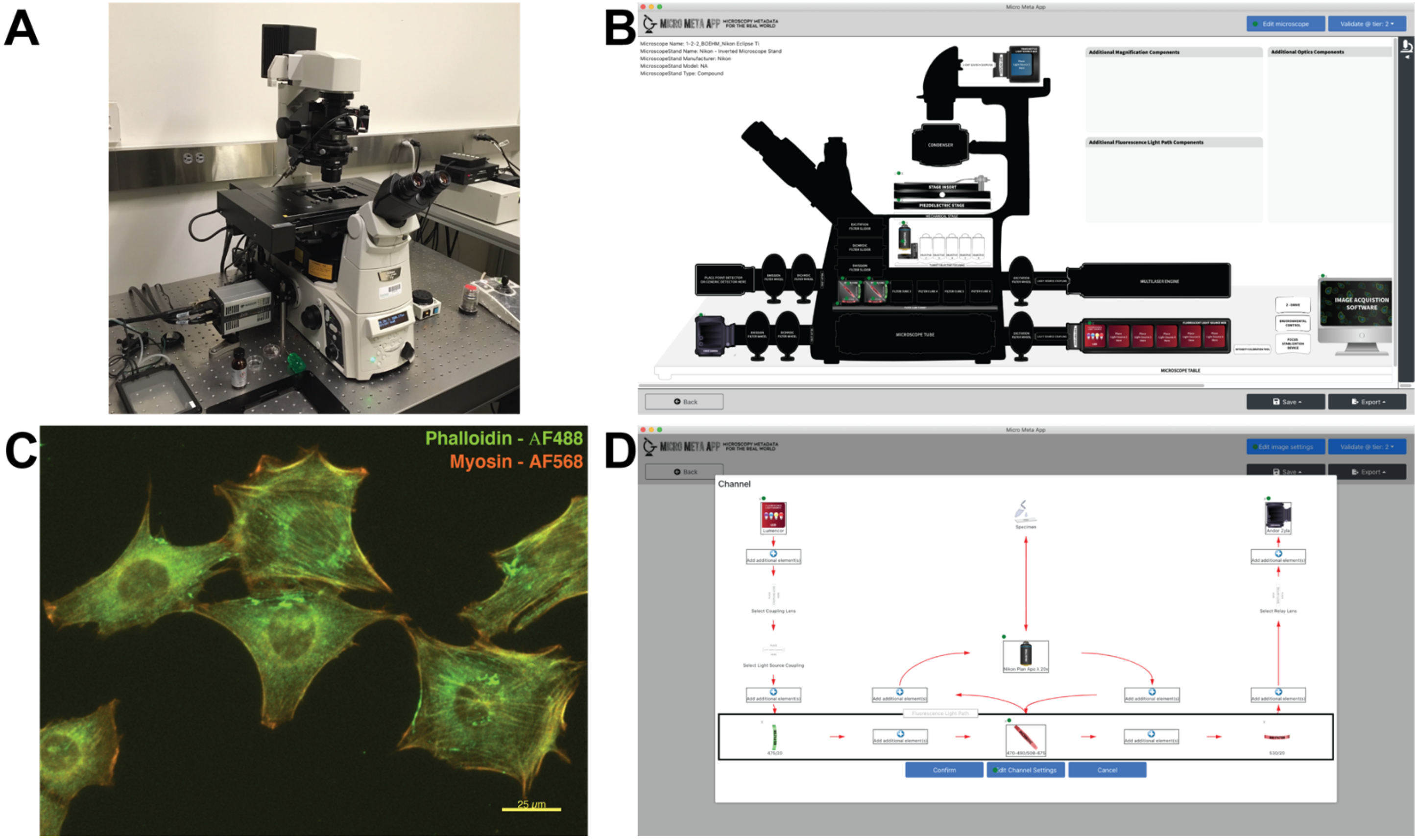

**Supplemental Figure 10 | Micro-Meta App was utilized by several light microscopy core facilities to document individual microscope instrumentation and the settings that were applied to the microscope to acquire specific example datasets**. Illustrated here is the use of Micro-Meta App for Tier 2 *(Advanced Quantification and/or Live cell Imaging)*, 4DN-BINA-OME-specified (Hammer et al., 2021) documentation of: **(A-B)** the ***Nikon Eclipse Ti*** inverted, epifluorescence microscope owned by the Advanced Imaging Center (AIC) at the Janelia Research Campus of the Howard Hughes Medical Institute (Supplemental Table III); and **(C-D)** the settings that were applied to the microscope for the acquisition of example acquired by Ulrike Boehm. **A)** Picture of the indicated microscope. **B)** Micro-Meta App generated schematic representation of the indicated microscope. **C)** COS cells were fixed, permeabilized, blocked, and stained with Alexa Fluor 488-conjugated Phalloidin (488 channel, visualized in green), and anti-Myosin antibodies (visualized with Alexa Fluor 568 conjugated secondary antibody; 568 channel, visualized in magenta). Displayed is a representative single plane image obtained using the indicated microscope (Panels A and B). **D)** Micro-Meta App generated schematic representation of the light path associated with the ***488 Channel*** utilized for the acquisition of the image in Panel C.

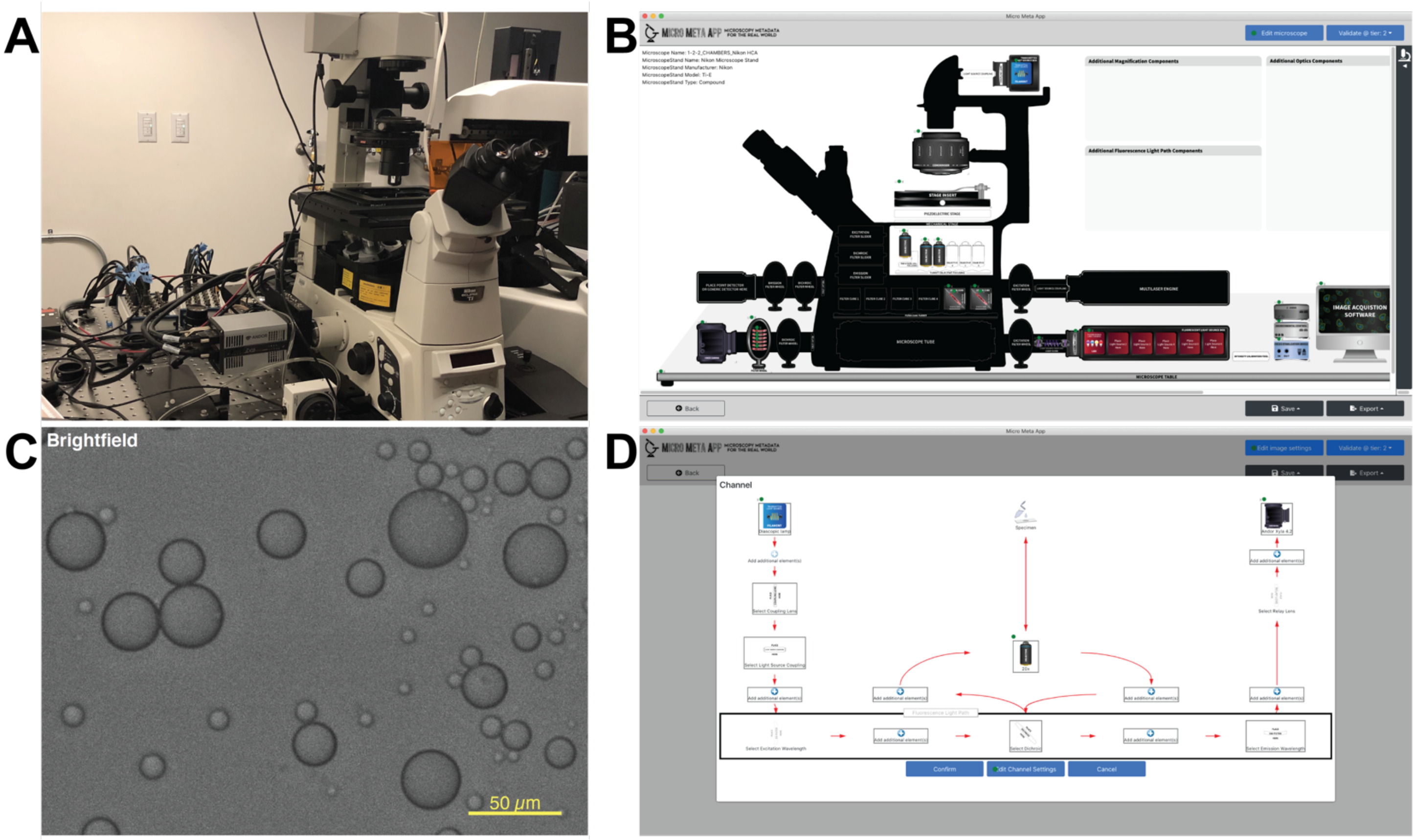

**Supplemental Figure 11 | Micro-Meta App was utilized by several light microscopy core facilities to document individual microscope instrumentation and the settings that were applied to the microscope to acquire specific example datasets**. Illustrated here is the use of Micro-Meta App for Tier 2 *(Advanced Quantification and/or Live cell Imaging)*, 4DN-BINA-OME-specified (Hammer et al., 2021) documentation of: **(A-B)** the ***Nikon Eclipse Ti-E*** inverted, epifluorescence microscope owned by the Light Microscopy Facility at the (IALS-LIF) Institute for Applied Life Sciences of the University of Massachusetts at Amherst (Supplemental Table III); and **(C-D)** the settings that were applied to the microscope for the acquisition of example published image data sets (Fernandez et al., 2020). **A)** Picture of the indicated microscope. **B)** Micro-Meta App generated schematic representation of the indicated microscope. **C)** Diascopic image of amine-functionalized vesicular assemblies before exposure to a component that results in bursting behavior in the study of new drug delivery macromolecular assemblies. This image is part of a time series (10 us exposure in 12-bit mode) using the 20x objective and is representative of the type of data collected to measure bursting behavior and dynamics. The displayed image was obtained using the indicated microscope (Panels A and B) and was adapted with permission from Fernandez et al., 2020. **D)** Micro-Meta App generated schematic representation of the light path associated with the ***Brightfield Channel*** utilized for the acquisition of the image in Panel C.

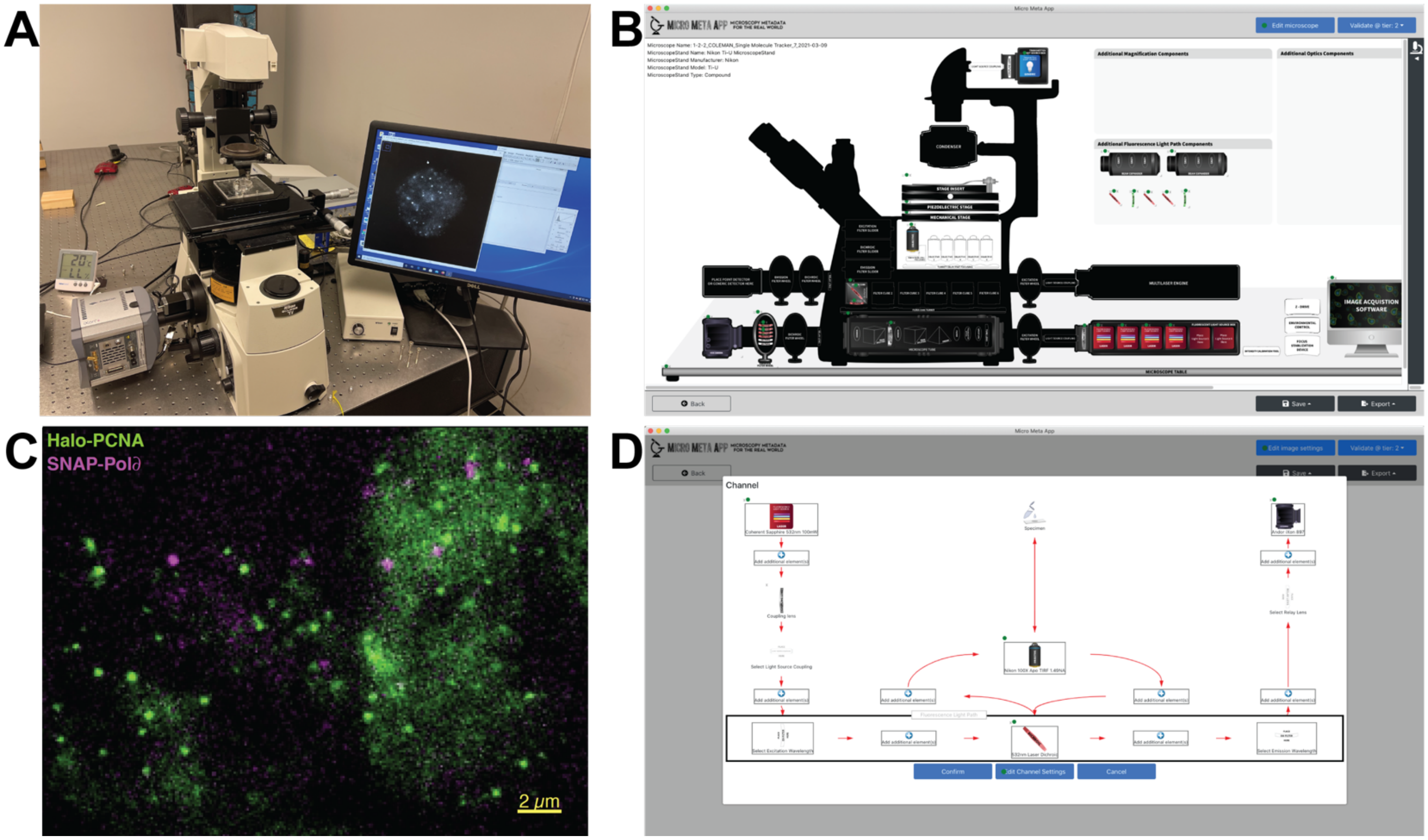

**Supplemental Figure 12 | Micro-Meta App was utilized by several light microscopy core facilities to document individual microscope instrumentation and the settings that were applied to the microscope to acquire specific example datasets**. Illustrated here is the use of Micro-Meta App for Tier 2 *(Advanced Quantification and/or Live cell Imaging)*, 4DN-BINA-OME-specified (Hammer et al., 2021) documentation of: **(A-B)** the custom build ***TIRF HILO Epifluorescence light Microscope (THEM; based on Nikon Eclipse Ti)*** developed, built and owned by the Coleman laboratory at the Anatomy and Structural Biology Department of The Albert Einstein College of Medicine (Supplemental Table III); and **(C-D)** the settings that were applied to the microscope for the acquisition of example published image data sets (Drosopoulos et al., 2020). **A)** Picture of the indicated microscope. **B)** Micro-Meta App generated schematic representation of the indicated microscope. **C)** LOX cells stably expressing Halo-PCNA and SNAP-Pol∂ were plated in selective media, incubated with 1mg/ml Doxycycline and incubated at 37C. Immediately prior to imaging, cells were incubated at 37C with JF549-HTL (Janelia Labs) and SNAP-Cell647-SiR (New England Biolabs), washed to remove unincorporated dye and placed in L-15 imaging media (Life technologies) + 10% FBS for imaging. Imaging sessions were carried out at room temperature and only cells displaying a punctate nuclear distribution pattern of PCNA were imaged. Cells were continuously illuminated using 532nm (13 W/cm2, Coherent) and 640nm (9.5 W/cm2, Coherent) lasers for JF549-HTL and SNAP-Cell 647-SiR imaging respectively. Two dimensional time-lapse images of single molecules were acquired using the indicated microscope (Panels A and B). Sequential dual color images of live cell nuclei were acquired for 22 minutes using 500 ms exposures on an EMCCD camera (iXon, Andor) with continuously alternating between the HALO-PCNA (labeled with JF549-HTL, displayed in red) and the SNAP-Pol∂ (labeled with SNAP-Cell 647-SiR, displayed in magenta) channels. The displayed image was obtained using the indicated microscope (Panels A and B) and was adapted with permission from Drosopoulos et al., 2020. **D)** Micro-Meta App generated schematic representation of the light path associated with the ***HALO-PCNA (JF549-HTL) Channel*** utilized for the acquisition of the image in Panel C.

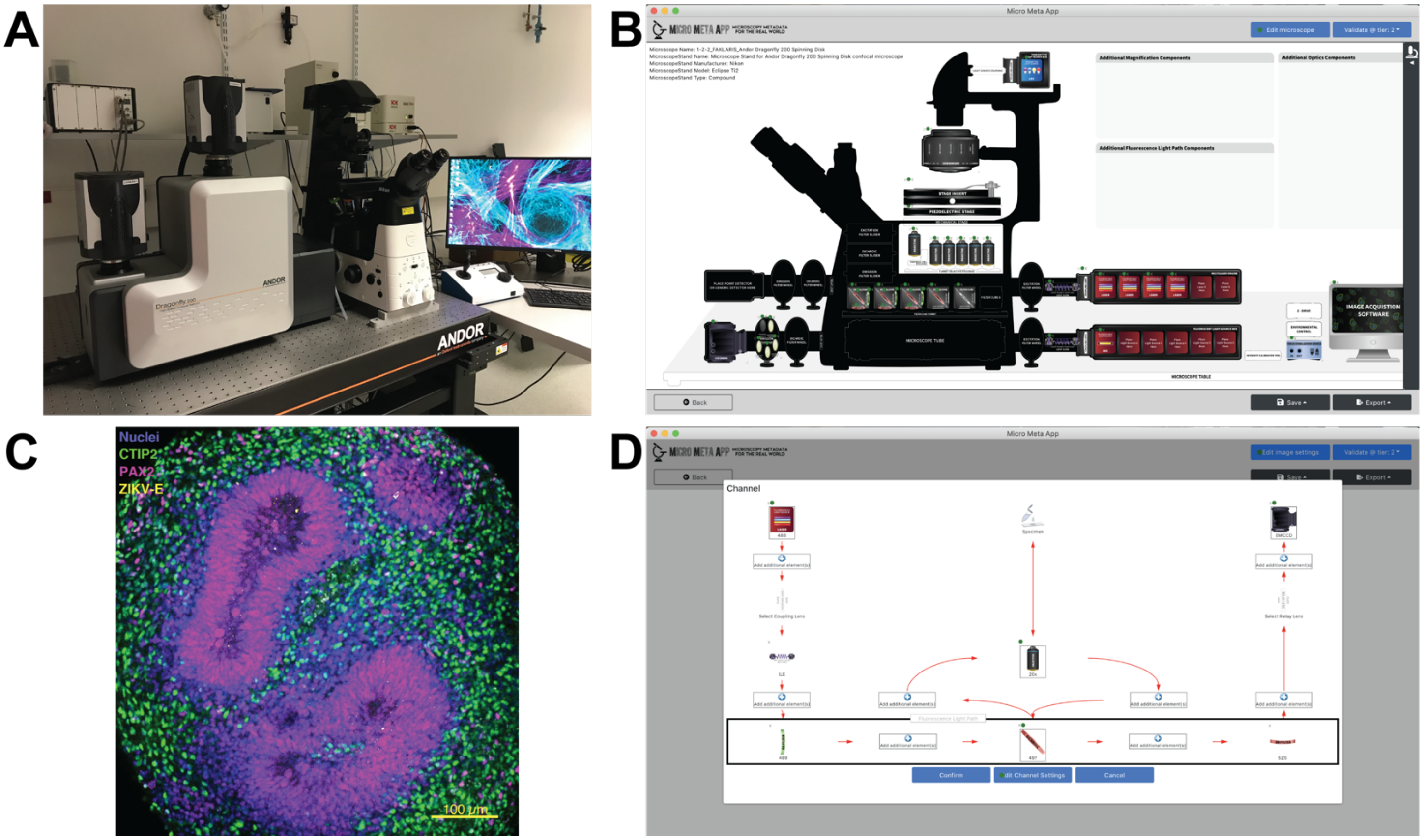

**Supplemental Figure 13 | Micro-Meta App was utilized by several light microscopy core facilities to document individual microscope instrumentation and the settings that were applied to the microscope to acquire specific example datasets**. Illustrated here is the use of Micro-Meta App for Tier 2 *(Advanced Quantification and/or Live cell Imaging)*, 4DN-BINA-OME-specified (Hammer et al., 2021) documentation of: **(A-B)** the ***Andor Dragonfly Spinning Disk confocal microscope system (with Nikon Eclipse Ti inverted compound microscope stand)*** owned by the Montpellier Resources Imagerie at the Centre de Recherche de Biologie cellulaire de Montpellier (MRI-CRBM) of the University of Montpellier (Supplemental Table III); and **(C-D)** the settings that were applied to the microscope for the acquisition of example published image data sets (Ayala-Nunez et al., 2019). **A)** Picture of the indicated microscope. **B)** Micro-Meta App generated schematic representation of the indicated microscope. **C)** Cerebral organoids were cocultured with ZIKV-infected monocytes, fixed, permeabilized, stained with DAPI to detect all cellular nuclei (405 channel, visualized in blue) anti-CTIP2 (488 channel, visualized in green), anti-PAX6 (568 channel, visualized in magenta) and anti-ZIKV2-E (Flavivirus group antigen Antibody (D1-4G2-4-15 (4G2); 640 channel, visualized in yellow), and clarified. 3D Z-stacks of representative organoids were acquired using the indicated microscope (Panels A and B) equipped with a 20x, NA 0.8, air Nikon objective. The displayed image corresponds to a representative Z-plane and was adapted with permission from Ayala-Nunez et al., 2019. **D)** Micro-Meta App generated schematic representation of the light path associated with the ***488 Channel*** utilized for the acquisition of the image in Panel C.

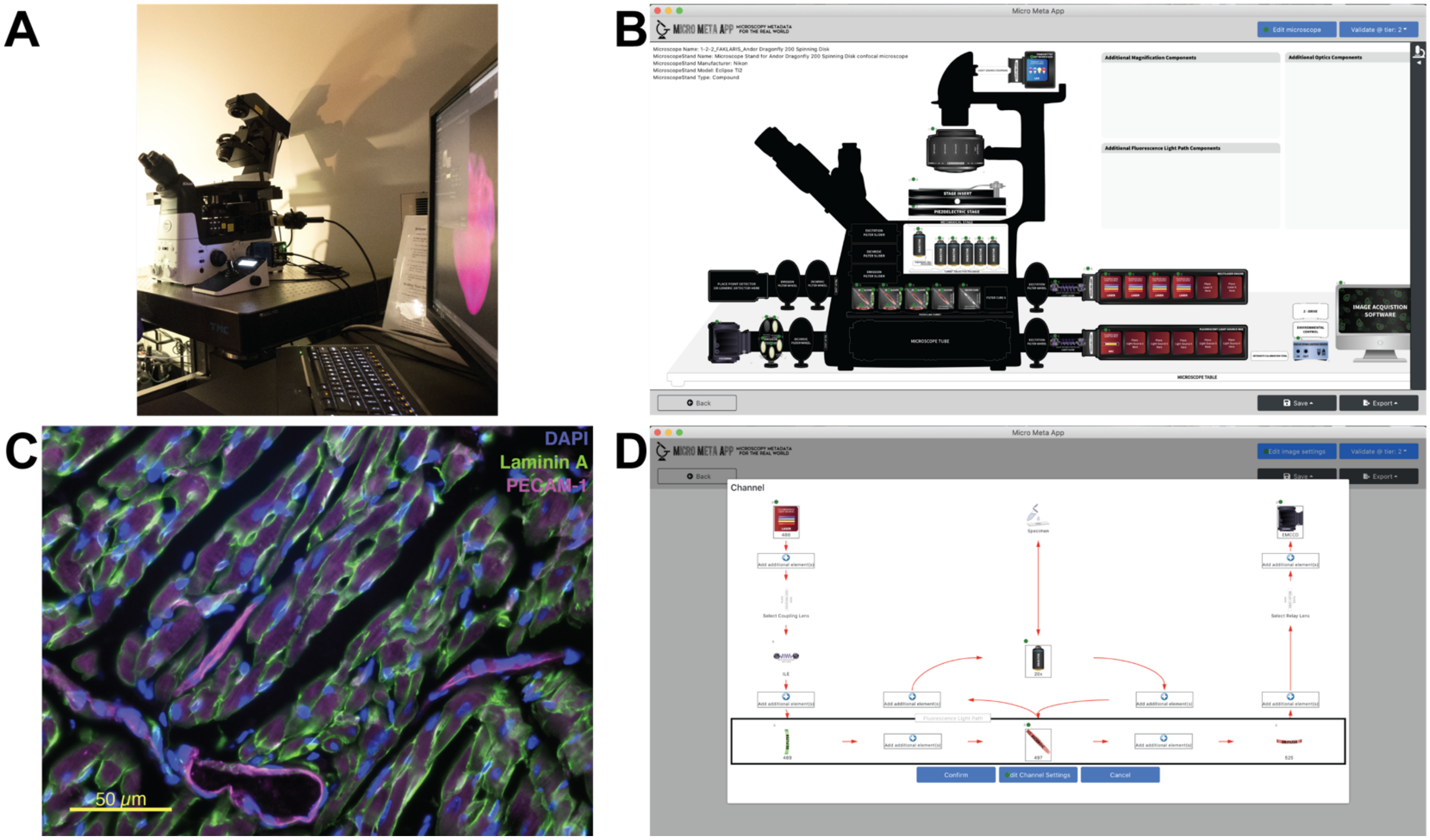

**Supplemental Figure 14 | Micro-Meta App was utilized by several light microscopy core facilities to document individual microscope instrumentation and the settings that were applied to the microscope to acquire specific example datasets.** Illustrated here is the use of Micro-Meta App for Tier 2 *(Advanced Quantification and/or Live cell Imaging)*, 4DN-BINA-OME-specified (Hammer et al., 2021) documentation of: **(A-B)** the ***Nikon Eclipse Ti2*** inverted, epifluorescence microscope owned by the Microscopy Core, at the Neuroscience Center of the University of North Carolina (Supplemental Table III); and **(C-D)** the settings that were applied to the microscope for the acquisition of example published image data sets (Aghajanian et al., 2021). **A)** Picture of the indicated microscope. **B)** Micro-Meta App generated schematic representation of the indicated microscope. **C)** Mouse hearth myocardium tissues from the left ventricles were fixed, cryo-protected with 30% sucrose overnight at 4C, embedded in OCT compound, and sectioned on a transverse plane to produce 8 μm sections. Sections were washed in PBS, permeabilized, blocked, and stained with rabbit antibodies against Laminin A (GFP Channel, visualized with Alexa Fluor 488 conjugated secondary antibody, displayed in green), rat antibodies against PECAM-1 (CD31; Texas Red Channel, visualized with Alexa Fluor 568 conjugated secondary antibody, displayed in magenta), and DAPI (displayed in blue). Displayed is a representative single plane image extracted from a 3D stack obtained from randomly selected areas of the tissue obtained using the indicated microscope (Panels A and B). The displayed image was obtained using the indicated microscope (Panels A and B) and was adapted with permission from Aghajanian et al., 2021. **D)** Micro-Meta App generated schematic representation of the light path associated with the ***Laminin A (Alexa Fluor 488) Channel*** utilized for the acquisition of the image in Panel C.

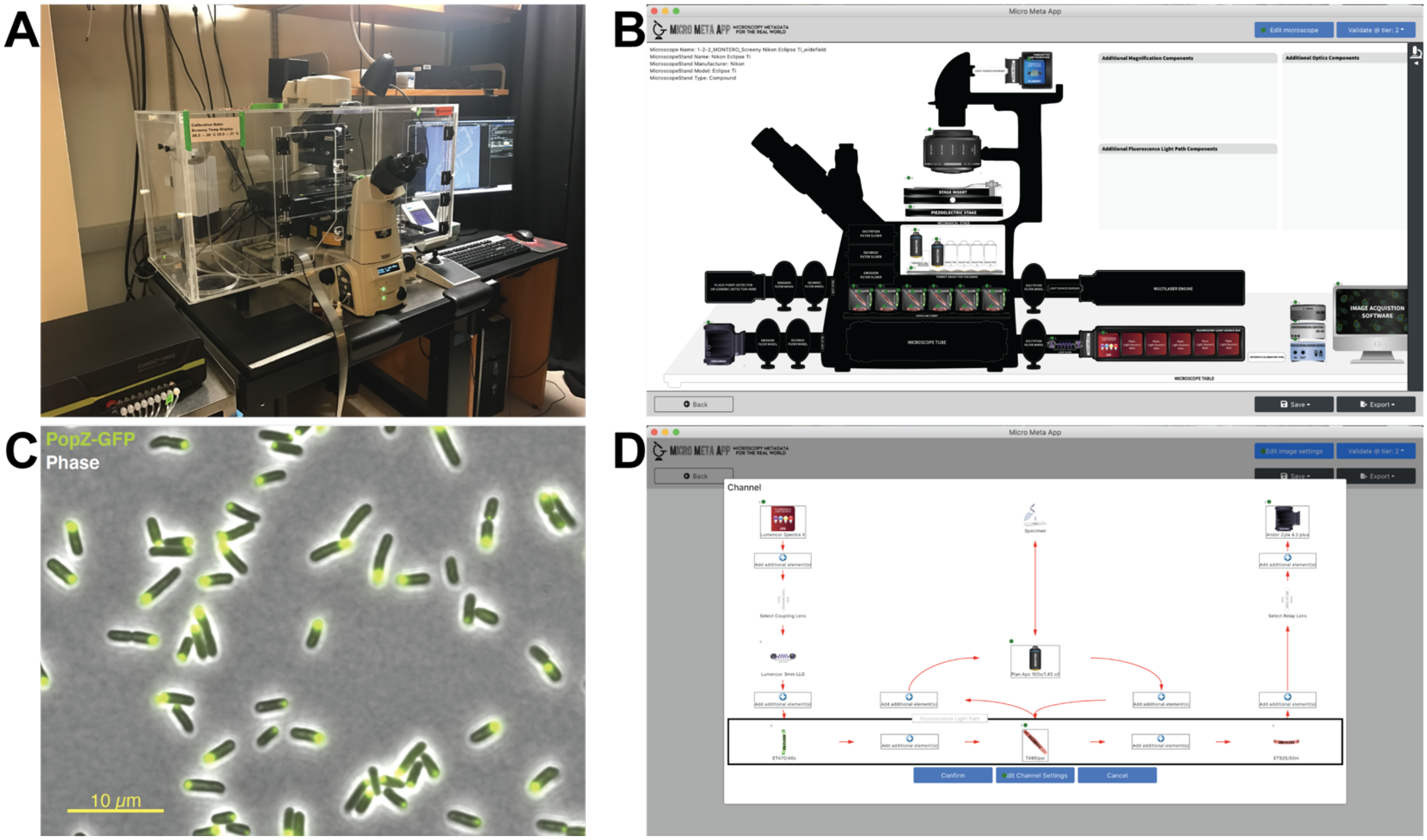

**Supplemental Figure 15 | Micro-Meta App was utilized by several light microscopy core facilities to document individual microscope instrumentation and the settings that were applied to the microscope to acquire specific example datasets**. Illustrated here is the use of Micro-Meta App for Tier 2 *(Advanced Quantification and/or Live cell Imaging)*, 4DN-BINA-OME-specified (Hammer et al., 2021) documentation of: **(A-B)** the ***Nikon_Eclipse Ti2*** inverted LED-based widefield microscope owned by the MicRoN Facility in the Department of Microbiology at Harvard Medical School and **(C-D)** the settings that were applied to the microscope for the acquisition of example published image data sets (Lim and Bernhardt 2019). **A)** Picture of the indicated microscope. **B)** Micro-Meta App generated schematic representation of the indicated microscope. **C)** *Escherichia coli* strain TB28 cells expressing PopZ-superfolderGFP were grown at 37 °C in minimal M9 medium supplemented with 0.2% casamino acids and 0.2% maltose (M9 Mal medium). Prior to imaging, 500 µL of the cell culture ($OD_{600}$ 0.2-0.4) was pelleted by gentle centrifugation (2 minutes at 4000 x g). Most of the supernatant (450 µL) was discarded. The cell pellet was subsequently resuspended in the remaining 50 $\mu$L of the original growth medium. One microliter of the resulting culture was spotted on 2% agarose pads made using M9 Mal medium and covered with #1.5 coverslips. Images were acquired using Nikon Elements 4.30 acquisition software. Overlay images show the GFP channel depicted in green, and the Phase contrast channel depicted in in grey. The displayed image was obtained using the indicated microscope (Panels A and B) and was adapted with permission from Lim and Bernhardt 2019. **D)** Micro-Meta App generated schematic representation of the light path associated with the ***PopZ-GFP Channel*** utilized for the acquisition of the image in Panel C.

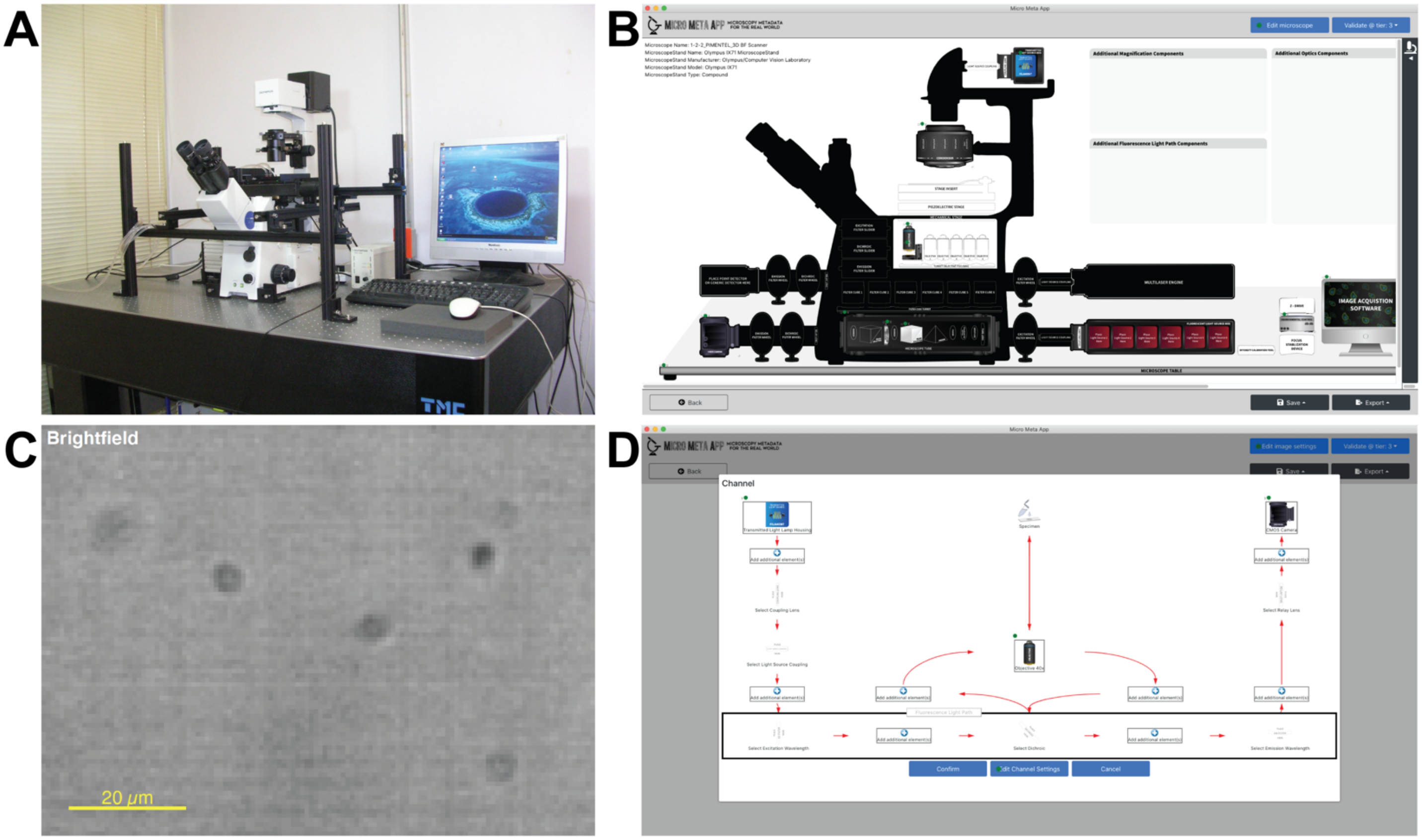

**Supplemental Figure 16 | Micro-Meta App was utilized by several light microscopy core facilities to document individual microscope instrumentation and the settings that were applied to the microscope to acquire specific example datasets**. Illustrated here is the use of Micro-Meta App for Tier 3 *(Manufacturing/Technical Development/Full Documentation)*, 4DN-BINA-OME-specified (Hammer et al., 2021) documentation of: **(A-B)** the custom build ***3D BrightField Scanner (based on Olympus IX71)*** inverted epifluorescence microscope developed and built by the Computer Vision Laboratory of the Institute of Biotechnology and owned in conjunction with the Laboratorio Nacional de Microscopia Avanzada (LNMA) at the Universidad Nacional Autonoma de Mexico (UNAM; Supplemental Table III); and **(C-D)** the settings that were applied to the microscope for the acquisition of example published image data sets (Pimentel et al., 2012; Silva-Villalobos et al., 2014). **A)** Picture of the indicated microscope. **B)** Micro-Meta App generated schematic representation of the indicated microscope. **C)** Sperm obtained from the sea urchin *Strongylocentrotus purpuratus* were diluted in Artificial Salt Water and imaged in Petri dishes used as imaging chambers. In order to capture sperm displacement information, images were continually acquired at 2000 fps using a Optronics CR5000x2 camera during continual oscillatory 3D objective focal scanning for a 250 μm depth trough the sperm sample at a frequency of 30 Hz. This acquisition rate combined with the piezo electric movement guarantees that one frame is acquired each 8μm in the z-direction. Displayed is a single focal plane from a representative 3D+t stack. Dark spots corresponds to unfocused swimming sea urchin sperm. Image obtained using the indicated microscope (Panels A and B) and adapted with permission from Pimentel et al., 2012; Silva-Villalobos et al., 2014. D) Micro-Meta App generated schematic representation of the light path associated with the ***Brightfield Channel*** utilized for the acquisition of the image in Panel C.

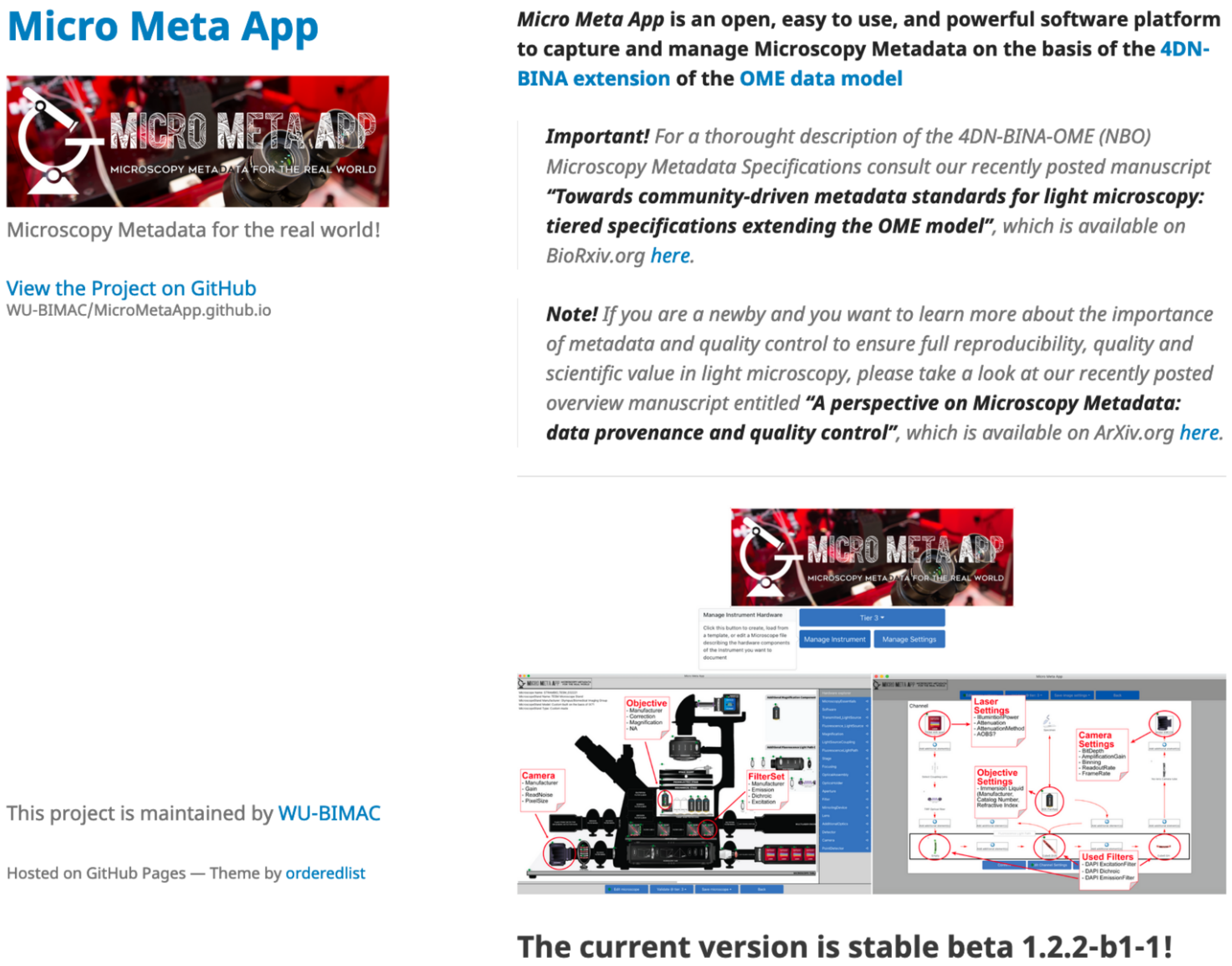

**Supplemental Figure 17 | Micro-Meta App website.** the Micro-Meta App website is available at https://wu-bimac.github.io/MicroMetaApp.github.io/ and was developed to promote outreach and adoption.

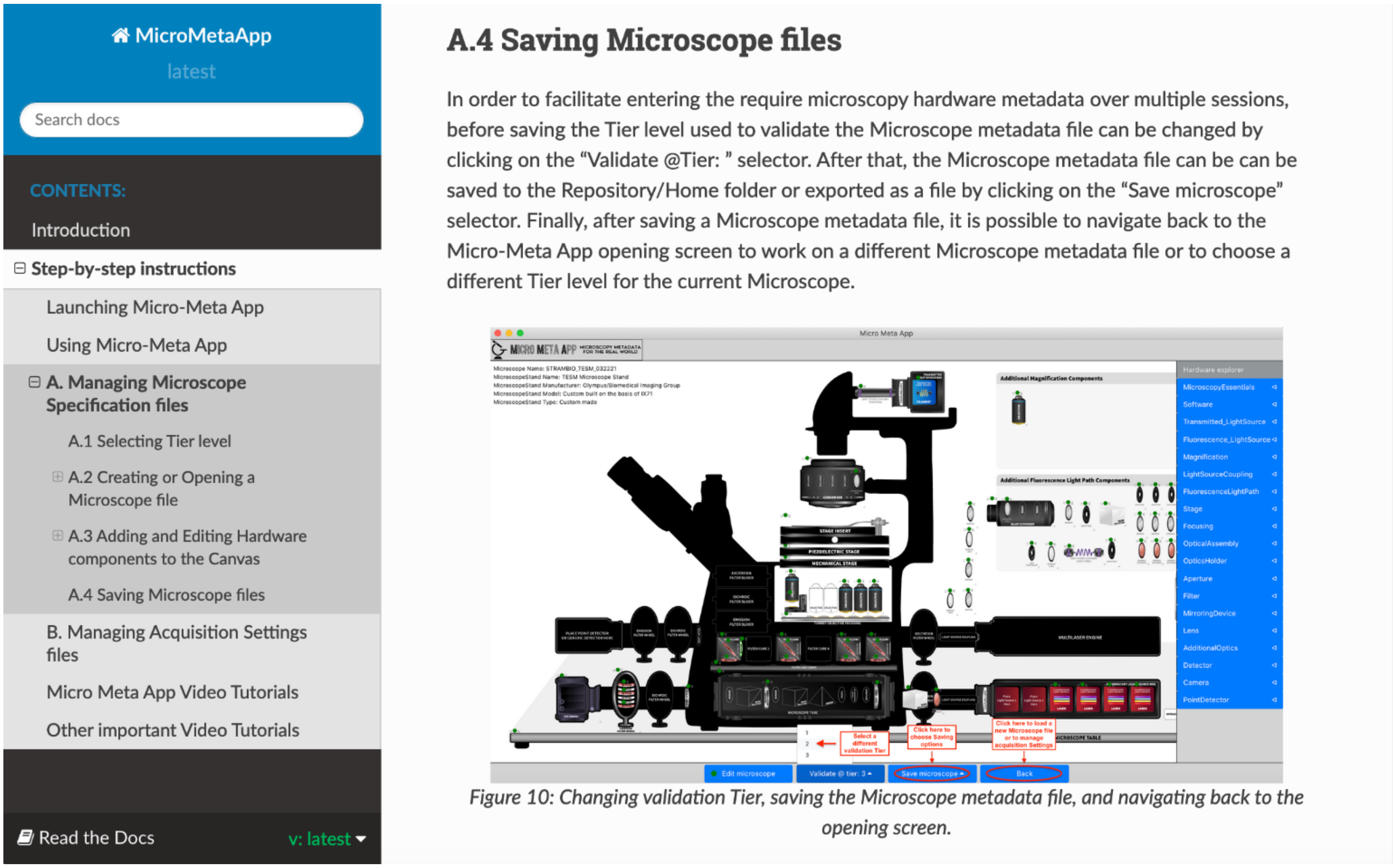

MicroMetaApp
latest

Search docs

CONTENTS:

Introduction

⊟ Step-by-step instructions

Launching Micro-Meta App

Using Micro-Meta App

⊟ A. Managing Microscope Specification files

A.1 Selecting Tier level

⊞ A.2 Creating or Opening a Microscope file

⊞ A.3 Adding and Editing Hardware components to the Canvas

A.4 Saving Microscope files

B. Managing Acquisition Settings files

Micro Meta App Video Tutorials

Other important Video Tutorials

Read the Docs v: latest

#### A.4 Saving Microscope files

In order to facilitate entering the require microscopy hardware metadata over multiple sessions, before saving the Tier level used to validate the Microscope metadata file can be changed by clicking on the "Validate @Tier: " selector. After that, the Microscope metadata file can be can be saved to the Repository/Home folder or exported as a file by clicking on the "Save microscope" selector. Finally, after saving a Microscope metadata file, it is possible to navigate back to the Micro-Meta App opening screen to work on a different Microscope metadata file or to choose a different Tier level for the current Microscope.

*Figure 10: Changing validation Tier, saving the Microscope metadata file, and navigating back to the opening screen.*

**Supplemental Figure 18 | Micro-Meta App Read the Docs documentation and tutorial site.** The Micro-Meta App Read the Docs documentation is available at https://micrometaapp-docs.readthedocs.io/en/latest/index.html. The site contains step-by-step instructions and video tutorials that explain how to use the App.

### SUPPLEMENTAL TABLES

#### Supplemental Table I

**Table I: Percentage of OME Data Model Core <Instrument> element attributes that can be retreived from image file headers and interpreted by BioFormats**

| OME Core Data Model | | Microscope Manufacturer | Nikon | Nikon | Zeiss | Leica | Leica | Customized Olympus | Customized Olympus |
|---|---|---|---|---|---|---|---|---|---|
| | | Acquisition/ Processing Software | Nikon NIS-Elements | Bitplane Imaris | Zeiss CZI | Leica LAS AF | MetaMorph Stack | Team Viewer | µManager |
| | | File format | .nd2 | .ims | .czi | .lif | .nd | .tif | ome.tif |
| | | Fraction of OME Core prescribed metadata fields that can be retreived from the file header | | | | | | | |
| **<Instrument> Sub-Element** | **Nr. of fields per element** | | | | | | | | |
| **Microscope** | **5** | | - | - | 20% | 40% | - | - | - |
| **Light Source / Laser** | **15** | | - | - | - | - | 20% | - | - |
| **Objective** | **12** | | 25% | - | 50 - 58% | 58% | 33% | - | - |
| **Filter Set** | **5** | | - | - | 40 -80% | - | - | - | - |
| **Filter** | **12** | | - | - | 0 - 25% | 8% | - | - | - |
| **Dichroic** | **5** | | - | - | 0 - 20% | - | - | - | - |
| **Detector** | **11** | | 27% | - | 18 - 45% | 27% | 18% | - | - |
| **ALL** | **65** | | **9%** | **-** | **18 - 26%** | **20%** | **14%** | **-** | **-** |

### Supplemental Table II

**Table II: Fraction of OME Data Model Core <Image> element attributes that can be retreived from image file headers and interpreted by BioFormats**

| OME Core Data Model | | | | | | | | | |
|---|---|---|---|---|---|---|---|---|---|
| | | Microscope Manufacturer | Nikon | Nikon | Zeiss | Leica | Leica | Customized Olympus | Customized Olympus |
| | | Acquisition/ Processing Software | Nikon NIS-Elements | Bitplane Imaris | Zeiss CZI | Leica LAS AF | MetaMorph Stack | Team Viewer | µManager |
| | | File format | .nd2 | .ims | .czi | .lif | .nd | .tif | ome.tif |
| | | Fraction of OME Core prescribed metadata fields that can be retreived from the file header | | | | | | | |
| **<Image> Sub-Element** | **Nr. of fields per element** | | | | | | | | |
| **Experiment** | **4** | | - | - | - | - | - | - | - |
| **Microbeam Manipulation** | **4** | | - | - | - | - | - | - | - |
| **Image** | **7** | | 43% | 29% | 57 - 71% | 57% | 57% | - | 43% |
| **Imaging Environment** | **5** | | - | - | - | - | 20% | - | - |
| **Objective Settings** | **4** | | 50% | - | 25% | 50% | 50% | - | - |
| **Light Source Settings** | **3** | | - | - | - | - | 67% | - | - |
| **Filter Set** | **1** | | - | - | 0 - 100% | - | - | - | - |
| **Light Path** | **3** | | - | - | - | 33% | - | - | - |
| **Detector Settings** | **9** | | 11 22% | - | 22 - 33% | 22% | 22% | - | - |
| **Stage Label** | **4** | | - | - | 100% | - | - | - | - |
| **Pixel** | **15** | | 87% | 93% | 87 - 93% | 87% | 93% | - | 93% |
| **Plane** | **9** | | 89% | - | 67 - 89% | 56% | 89% | - | - |
| **Channel** | **13** | | 38% | 23% | 62 -69% | 31% | 31% | - | 31% |
| **ALL** | **81** | | **37 - 41%** | **24%** | **53 - 58%** | **40%** | **47%** | - | **27%** |

### Supplemental Table III

**Supplemental Table III: List of Microscopes documented using Micro-Meta App**

| Nr. | Manufacturer | Model | Figure | Tier | Type | Illumination | Facility Name | Department and Institution | URL | References |
|---|---|---|---|---|---|---|---|---|---|---|
| 1 | Carl Zeiss Microscopy | Axio Observer Z1 (with LSM 710 scanhead) | Supp. Fig. 2 | 1 | Compound _Inverted | Confocal (Laser Scanning) | Centre for Cell Imaging (CCI) | University of Liverpool | https://cci.liv.ac.uk/equipment_710.html | Upton et al., 2020 |
| 2 | Carl Zeiss Microscopy | Axio Observer (Axiovert 200M) | Supp. Fig. 3 | 2 | Compound _Inverted | Epifluorescence | Advanced BioImaging Facility (ABIF). | McGill University | https://www.mcgill.ca/abif/equipment/axiovert-1 | Kiepas et al., 2020 |
| 3 | Carl Zeiss Microscopy | Axio Observer Z1 (with Spinning Disk) | Supp. Fig. 4 | 2 | Compound _Inverted | Confocal (Spinning Disk) | Imagerie Cellulaire; Quality Control managed by Miacellavie (https://miacellavie.com/) | Centre de recherche du Centre Hospitalier Université de Montréal (CR CHUM), University of Montreal | https://www.chumontreal.qc.ca/crchum/plateformes-et-services (the web site is for all core facilities, not specifically for the core facility hosting this microscope) | Pilliod et al., 2020 |
| 4 | Carl Zeiss Microscopy | Axio Imager Z2 (with Apotome) | Supp. Fig. 5 | 2 | Compound _Upright | Epifluorescence | Bioimaging Unit | Newcastle University | https://www.ncl.ac.uk/bioimaging/ | Watson et al., 2020 |
| 5 | Carl Zeiss Microscopy | Axio Observer Z1 | Supp. Fig. 6 | 2 | Compound _Inverted | Epifluorescnce | Life Imaging Center (LIC) | Centre for Integrative Signalling Analysis (CISA), University of Freiburg | https://miap.eu/equipments/sd-i-abl/ | Hannibal et al., 2020 |
| 6 | Leica Microsystems | DMI6000 | Supp. Fig. 7 | 2 | Compound _Inverted | Epifluorescnce | IMAG'IC Confocal Microscopy Facility | Institut Cochin, CNRS, INSERM, Université de Paris | https://www.institutcochin.fr/core_facilities/confocal-microscopy/cochin-imaging-photonic-microscopy/organigram_team/10054/view | Dumas et al., 2015; Jubrail et al., 2020 |
| 7 | Leica Microsystems | DM5550B | Supp. Fig. 8 | 2 | Compound _Upright | Epifluorescence | Bioimaging Unit | Edwardson Building on the Campus for Ageing and Vitality, Newcastle University | https://www.ncl.ac.uk/bioimaging/equipment/leica-dm5500/#overview | da Silva et al., 2019; Nelson et al., 2012 |
| 8 | Leica Microsystems | DMI8-CS (with TCS SP8 STED 3X) | Supp. Fig. 9 | 2 | Compound _Inverted | Confocal (Laser Scanning) | Center for Advanced imaging (CAi) | School of Mathematics/Natural Sciences, Heinrich-Heine-Universität Düsseldorf | https://www.cai.hhu.de/en/equipment/super-resolution-microscopy/leica-tcs-sp8-sted-3x | Singer et al., 2017; Hänsch et al., 2020 |
| 9 | Nikon Instruments | Eclipse Ti | Supp. Fig. 10 | 2 | Compound _Inverted | Epifluorescence | Advanced Imaging Center (AIC) | Janelia Research Campus, Howard Hughes Medical Institute | https://www.janelia.org/support-team/light-microscopy/equipment | Abdelfattah et al., 2019; Qian et al., 2019; Grimm et al., 2020 |
| 10 | Nikon Instruments | Eclipse Ti-E | Supp. Fig. 11 | 2 | Compound _Inverted | Epifluorescence | Light Microscopy Facility (IALS-LIF) | Institute for Applied Life Sciences, University of Massachusetts at Amherst | https://www.umass.edu/ials/light-microscopy | Fernandez et al., 2020 |
| 11 | Nikon Instruments/Coleman laboratory (customized) | TIRF HILO Epifluorescence light Microscope (THEM)/ Eclipse Ti | Supp. Fig. 12 | 2 | Compound _Inverted | Epifluorescence | Coleman laboratory | Anatomy and Structural Biology Department, The Albert Einstein College of Medicine | https://einsteinmed.org/faculty/12252/robert-coleman/ | Drosopoulos et al., 2020 |
| 12 | Nikon Instruments | Eclipse Ti (with Andor Dragon Fly Spinning Disk) | Supp. Fig. 13 | 2 | Compound _Inverted | Confocal (Spinning Disk) | Montpellier Resources Imagerie | Centre de Recherche de Biologie cellulaire de Montpellier (MRI-CRBM), CNRS, Univerity of Montpellier | https://www.mri.cnrs.fr/en/optical-imaging/our-facilities/mri-crbm.html | Ayala-Nunez et al., 2019 |
| 13 | Nikon Instruments | Eclipse Ti2 | Supp. Fig. 14 | 2 | Compound _Inverted | Epifluorescnce | Neuroscience Center Microscopy Core | Neuroscience Center, University of North Carolina | https://www.med.unc.edu/neuroscience/core-facilities/neuro-microscopy/ | Aghajanian et al., 2021 |
| 14 | Nikon Instruments | Eclipse Ti2 | Supp. Fig. 15 | 2 | Compound _Inverted | Epifluorescnce | Microscopy Resources on the North Quad (MicRoN) | Harvard Medical School | https://micron.hms.harvard.edu/ | Lim and Bernhardt 2019; Lim et al., 2019 |
| 15 | Olympus/Biomedical Imaging Group (customized) | TIRF Epifluorescence Structured light Microscope (TESM)/IX71 | Figure 5 | 3 | Compound _Inverted | Epifluorescnce | Biomedical Imaging Group | Program in Molecular Medicine, University of Massachusetts Medical School | https://trello.com/b/BQ8zCcQC/tirf-epi-fluorescence-structured-light-microscope | Navaroli et al., 2012 |
| 16 | Olympus/Computer Vision Laboratory (customized) | 3D BrightField Scanner/IX71 | Supp. Fig. 16 | 3 | Compound _Inverted | Brightfield | Laboratorio Nacional de Microscopia Avanzada (LNMA) and Computer Vision Laboratory of the Institute of Biotechnology | Universidad Nacional Autonoma de Mexico (UNAM) | https://lnma.unam.mx/wp/ | Pimentel et al., 2012; Silva-Villalobos et al., 2014 |

### SUPPLEMENTAL REFERENCES

Abdelfattah, A.S., T. Kawashima, A. Singh, O. Novak, H. Liu, Y. Shuai, Y.-C. Huang, L. Campagnola, S.C. Seeman, J. Yu, J. Zheng, J.B. Grimm, R. Patel, J. Friedrich, B.D. Mensh, L. Paninski, J.J. Macklin, G.J. Murphy, K. Podgorski, B.-J. Lin, T.-W. Chen, G.C. Turner, Z. Liu, M. Koyama, K. Svoboda, M.B. Ahrens, L.D. Lavis, and E.R. Schreiter. 2019. Bright and photostable chemigenetic indicators for extended in vivo voltage imaging. *Science*. 365:699–704. doi:10.1126/science.aav6416.

Aghajanian, A., H. Zhang, B.K. Buckley, E.S. Wittchen, W.Y. Ma, and J.E. Faber. 2021. Decreased inspired oxygen stimulates de novo formation of coronary collaterals in adult heart. *J. Mol. Cell. Cardiol.* 150:1–11. doi:10.1016/j.yjmcc.2020.09.015.

Ayala-Nunez, N.V., G. Follain, F. Delalande, A. Hirschler, E. Partiot, G.L. Hale, B.C. Bollweg, J. Roels, M. Chazal, F. Bakoa, M. Carocci, S. Bourdoulous, O. Faklaris, S.R. Zaki, A. Eckly, B. Uring-Lambert, F. Doussau, S. Cianferani, C. Carapito, F.M.J. Jacobs, N. Jouvenet, J.G. Goetz, and R. Gaudin. 2019. Zika virus enhances monocyte adhesion and transmigration favoring viral dissemination to neural cells. *Nat. Commun.* 10:4430. doi:10.1038/s41467-019-12408-x.

Drosopoulos, W.C., D.A. Vierra, C.A. Kenworthy, R.A. Coleman, and C.L. Schildkraut. 2020. Dynamic Assembly and Disassembly of the Human DNA Polymerase δ Holoenzyme on the Genome In Vivo. *Cell Rep.* 30:1329–1341.e5. doi:10.1016/j.celrep.2019.12.101.

Dumas, A., G. Lê-Bury, F. Marie-Anaïs, F. Herit, J. Mazzolini, T. Guilbert, P. Bourdoncle, D.G. Russell, S. Benichou, A. Zahraoui, and F. Niedergang. 2015. The HIV-1 protein Vpr impairs phagosome maturation by controlling microtubule-dependent trafficking. *J. Cell Biol.* 211:359–372. doi:10.1083/jcb.201503124.

Fernandez, A., C.A. Zentner, M. Shivrayan, E. Samson, S. Savagatrup, J. Zhuang, T.M. Swager, and S. Thayumanavan. 2020. Programmable Emulsions via Nucleophile-Induced Covalent Surfactant Modifications. *Chem. Mater.* 32:4663–4671. doi:10.1021/acs.chemmater.0c01107.

Goldberg, I.G., C. Allan, J.-M. Burel, D. Creager, A. Falconi, H. Hochheiser, J. Johnston, J. Mellen, P.K. Sorger, and J.R. Swedlow. 2005. The Open Microscopy Environment (OME) Data Model and XML file: open tools for informatics and quantitative analysis in biological imaging. *Genome Biol.* 6:R47.

Grimm, J.B., A.N. Tkachuk, L. Xie, H. Choi, B. Mohar, N. Falco, K. Schaefer, R. Patel, Q. Zheng, Z. Liu, J. Lippincott-Schwartz, T.A. Brown, and L.D. Lavis. 2020. A general method to optimize and functionalize red-shifted rhodamine dyes. *Nat. Methods*. 17:815–821. doi:10.1038/s41592-020-0909-6.

Hammer, M., M. Huisman, A. Rigano, U. Boehm, J.J. Chambers, N. Gaudreault, J.A. Pimentel, D. Sudar, P. Bajcsy, C.M. Brown, A.D. Corbett, O. Faklaris, J. Lacoste, A. Laude, G. Nelson, R. Nitschke, A.J. North, R. Gopinathan, F. Farzam, C. Smith, D. Grunwald, and C. Strambio-De-Castillia. 2021. Towards community-driven metadata standards for light microscopy: tiered specifications extending the OME model. *BioRxiv*. doi:10.1101/2021.04.25.441198v1.

Hannibal, L., J. Theimer, V. Wingert, K. Klotz, I. Bierschenk, R. Nitschke, U. Spiekerkoetter, and S.C. Grünert. 2020. Metabolic Profiling in Human Fibroblasts Enables Subtype Clustering in Glycogen Storage Disease. *Front. Endocrinol.* . 11:579981. doi:10.3389/fendo.2020.579981.

Hänsch, S., D. Spona, G. Murra, K. Köhrer, A. Subtil, A.R. Furtado, S.F. Lichtenthaler, B. Dislich, K. Mölleken, and J.H. Hegemann. 2020. Chlamydia-induced curvature of the host-cell plasma membrane is required for infection. *Proc. Natl. Acad. Sci. U. S. A.* 117:2634–2644. doi:10.1073/pnas.1911528117.

Huisman, M., M. Hammer, A. Rigano, U. Boehm, J.J. Chambers, N. Gaudreault, J.A. Pimentel, D. Sudar, P. Bajcsy, C.M. Brown, A.D. Corbett, O. Faklaris, J. Lacoste, A. Laude, G. Nelson, R. Nitschke, A.J. North, D. Grunwald, and C. Strambio-De-Castillia. 2021. A perspective on Microscopy Metadata: data provenance and quality control. *arXiv [q-bio.QM]*.

Jubrail, J., K. Africano-Gomez, F. Herit, A. Mularski, P. Bourdoncle, L. Oberg, E. Israelsson, P.-R. Burgel, G. Mayer, D.M. Cunoosamy, N. Kurian, and F. Niedergang. 2020. Arpin is critical for phagocytosis in macrophages and is targeted by human rhinovirus. *EMBO Rep.* 21:e47963. doi:10.15252/embr.201947963.

Kiepas, A., E. Voorand, F. Mubaid, P.M. Siegel, and C.M. Brown. 2020. Optimizing live-cell fluorescence imaging conditions to minimize phototoxicity. *J. Cell Sci.* 133. doi:10.1242/jcs.242834.

Lim, H.C., and T.G. Bernhardt. 2019. A PopZ-linked apical recruitment assay for studying protein-protein interactions in the bacterial cell envelope. *Mol. Microbiol.* 112:1757–1768. doi:10.1111/mmi.14391.

Lim, H.C., J.W. Sher, F.P. Rodriguez-Rivera, C. Fumeaux, C.R. Bertozzi, and T.G. Bernhardt. 2019. Identification of new components of the RipC-FtsEX cell separation pathway of Corynebacterineae. *PLoS Genet.* 15:e1008284. doi:10.1371/journal.pgen.1008284.

Linkert, M., C.T. Rueden, C. Allan, J.-M. Burel, W. Moore, A. Patterson, B. Loranger, J. Moore, C. Neves, D. MacDonald, A. Tarkowska, C. Sticco, E. Hill, M. Rossner, K.W. Eliceiri, and J.R. Swedlow. 2010. Metadata matters: access to image data in the real world. *J. Cell Biol.* 189:777–782.

Moore, J., C. Allan, S. Besson, J.-M. Burel, E. Diel, D. Gault, K. Kozlowski, D. Lindner, M. Linkert, T. Manz, W. Moore, C. Tischer, and J.R. Swedlow. 2021. OME-NGFF: scalable format strategies for interoperable bioimaging data. *bioRxiv*. 2021.03.31.437929. doi:10.1101/2021.03.31.437929.

Navaroli, D.M., K.D. Bellve, C. Standley, L.M. Lifshitz, J. Cardia, D. Lambright, D. Leonard, K.E. Fogarty, and S. Corvera. 2012. Rabenosyn-5 defines the fate of the transferrin receptor following clathrin-mediated endocytosis. *Proceedings of the National Academy of Sciences*. 109:E471–80. doi:10.1073/pnas.1115495109.

Nelson, G., J. Wordsworth, C. Wang, D. Jurk, C. Lawless, C. Martin-Ruiz, and T. von Zglinicki. 2012. A senescent cell bystander effect: senescence-induced senescence. *Aging Cell*. 11:345–349. doi:10.1111/j.1474-9726.2012.00795.x.

Pilliod, J., A. Desjardins, C. Pernègre, H. Jamann, C. Larochelle, E.A. Fon, and N. Leclerc. 2020. Clearance of intracellular tau protein from neuronal cells via VAMP8-induced secretion. *J. Biol. Chem.* 295:17827–17841. doi:10.1074/jbc.RA120.013553.

Pimentel, J.A., J. Carneiro, A. Darszon, and G. Corkidi. 2012. A segmentation algorithm for automated tracking of fast swimming unlabelled cells in three dimensions. *J. Microsc.* 245:72–81. doi:10.1111/j.1365-2818.2011.03545.x.

Qian, Y., K.D. Piatkevich, B. Mc Larney, A.S. Abdelfattah, S. Mehta, M.H. Murdock, S. Gottschalk, R.S. Molina, W. Zhang, Y. Chen, J. Wu, M. Drobizhev, T.E. Hughes, J. Zhang, E.R. Schreiter, S. Shoham, D. Razansky, E.S. Boyden, and R.E. Campbell. 2019. A genetically encoded near-infrared fluorescent calcium ion indicator. *Nat. Methods*. 16:171–174. doi:10.1038/s41592-018-0294-6.

Rigano, A., A. Balashov, S. Ehmsen, S.U. Ozturk, B. Alver, and C. Strambio-De-Castillia. 2021a. Micro-Meta App - React. Github - https://github.com/WU-BIMAC.

Rigano, A., A. Balashov, S. Ehmsen, S.U. Ozturk, B. Alver, and C. Strambio-De-Castillia. 2021b. Micro-Meta App - Electron. Github - https://github.com/WU-BIMAC.

Rigano, A., U. Boehm, J.J. Chambers, N. Gaudreault, A.J. North, J.A. Pimentel, D. Sudar, P. Bajcsy, C.M. Brown, A.D. Corbett, O. Faklaris, J. Lacoste, A. Laude, G. Nelson, R. Nitschke, D. Grunwald, and C. Strambio-De-Castillia. 2021c. 4DN-BINA-OME (NBO) Tiered Microscopy Metadata Specifications - v2.01. https://github.com/WU-BIMAC.

Rigano, A., S.U. Öztürk, S. Ehmsen, A. Cosolo, A. Balashov, B. Alver, and C. Strambio-De-Castillia. 2021d. Micro-Meta App – 4DN Data Portal.

Rigano, A., and C. Strambio-De-Castillia. 2021a. 4DN Microscopy Metadata Reader. Github - https://github.com/WU-BIMAC.

Rigano, A., and C. Strambio-De-Castillia. 2021b. 4DN Metadata Schema XSD to JSON Converter. Github - https://github.com/WU-BIMAC.

da Silva, P.F.L., M. Ogrodnik, O. Kucheryavenko, J. Glibert, S. Miwa, K. Cameron, A. Ishaq, G. Saretzki, S. Nagaraja-Grellscheid, G. Nelson, and T. von Zglinicki. 2019. The bystander effect contributes to the accumulation of senescent cells in vivo. *Aging Cell*. 18:e12848. doi:10.1111/acel.12848.

Silva-Villalobos, F., J.A. Pimentel, A. Darszon, and G. Corkidi. 2014. Imaging of the 3D dynamics of flagellar beating in human sperm. *Conf. Proc. IEEE Eng. Med. Biol. Soc.* 2014:190–193. doi:10.1109/EMBC.2014.6943561.

Singer, A., G. Poschmann, C. Mühlich, C. Valadez-Cano, S. Hänsch, V. Hüren, S.A. Rensing, K. Stühler, and E.C.M. Nowack. 2017. Massive Protein Import into the Early-Evolutionary-Stage Photosynthetic Organelle of the Amoeba Paulinella chromatophora. *Curr. Biol.* 27:2763–2773.e5. doi:10.1016/j.cub.2017.08.010.

The Apache Software Foundation. 2018. Xerces2 Java XML Parser 2.12.1 Release.

Upton, R.L., Z. Davies-Manifold, M. Marcello, K. Arnold, and C.R. Crick. 2020. A general formulation approach for the fabrication of water repellent materials: how composition can impact resilience and functionality. *Mol. Syst. Des. Eng.* 5:477–483. doi:10.1039/C9ME00144A.

W3C. 2003. W3C Java XML bindings libraries.

Watson, N.A., T.N. Cartwright, C. Lawless, M. Cámara-Donoso, O. Sen, K. Sako, T. Hirota, H. Kimura, and J.M.G. Higgins. 2020. Kinase inhibition profiles as a tool to identify kinases for specific phosphorylation sites. *Nat. Commun.* 11:1684. doi:10.1038/s41467-020-15428-0.

Wilkinson, M.D., M. Dumontier, I.J.J. Aalbersberg, G. Appleton, M. Axton, A. Baak, N. Blomberg, J.-W. Boiten, L.B. da Silva Santos, P.E. Bourne, J. Bouwman, A.J. Brookes, T. Clark, M. Crosas, I. Dillo, O. Dumon, S. Edmunds, C.T. Evelo, R. Finkers, A. Gonzalez-Beltran, A.J.G. Gray, P. Groth, C. Goble, J.S. Grethe, J. Heringa, P.A.C. 't Hoen, R. Hooft, T. Kuhn, R. Kok, J. Kok, S.J. Lusher, M.E. Martone, A. Mons, A.L. Packer, B. Persson, P. Rocca-Serra, M. Roos, R. van Schaik, S.-A. Sansone, E. Schultes, T. Sengstag, T. Slater, G. Strawn, M.A. Swertz, M. Thompson, J.

van der Lei, E. van Mulligen, J. Velterop, A. Waagmeester, P. Wittenburg, K. Wolstencroft, J. Zhao, and B. Mons. 2016. The FAIR Guiding Principles for scientific data management and stewardship. *Sci Data*. 3:160018. doi:10.1038/sdata.2016.18.
